# A putative origin of insect chemosensory receptors in the last common eukaryotic ancestor

**DOI:** 10.1101/2020.08.24.264408

**Authors:** Richard Benton, Christophe Dessimoz, David Moi

## Abstract

The insect chemosensory repertoires of Gustatory Receptors (GRs) and Odorant Receptors (ORs) together represent one of the largest families of ligand-gated ion channels. Previous analyses have identified homologous “Gustatory Receptor-Like (GRL)” proteins across Animalia, but the evolutionary origin of this novel class of ion channels is unknown. We describe a survey of unicellular eukaryotic genomes for GRLs, identifying several candidates in fungi, protists and algae that contain many structural features characteristic of animal GRLs. The existence of these proteins in unicellular eukaryotes, together with *ab initio* protein structure predictions, supports homology between GRLs and a large family of uncharacterised plant proteins containing the DUF3537 domain. Together, this evidence suggests an origin of this protein family in the last common eukaryotic ancestor.

## Introduction

The insect chemosensory receptor superfamily, comprising Gustatory Receptors (GRs) and Odorant Receptors (ORs) form a critical molecular interface between the diverse chemical signals in the environment and neural activity patterns that evoke adaptive behavioural responses (Rytz, et al. 2013; Benton 2015; Joseph and Carlson 2015; van Giesen and Garrity 2017; Robertson 2019). Most insect genomes encode dozens to hundreds of different, often species-specific, GRs and/or ORs. Detailed analyses, in particular in *Drosophila melanogaster,* indicate that the vast majority of these are likely to be expressed in, and define the chemical response properties of, distinct subpopulations of peripheral sensory neurons (Vosshall and Stocker 2007; Joseph and Carlson 2015; Scott 2018; Chen and Dahanukar 2020).

Insect GRs and ORs – the latter having derived from an ancestral GR (Robertson, et al. 2003; Robertson 2019) – contain seven transmembrane (TM) domains (Clyne, et al. 1999; Vosshall, et al. 1999; Clyne, et al. 2000; Scott, et al. 2001). In contrast to vertebrate olfactory and taste receptors, which belong to the G protein-coupled receptor (GPCR) superfamily of seven TM domain proteins (Yarmolinsky, et al. 2009; Glezer and Malnic 2019), insect ORs and GRs have the opposite topology, with an intracellular N-terminus (Benton, et al. 2006; Lundin, et al. 2007). Functional analyses of these proteins in heterologous expression systems indicate that they form ligand-gated ion channels (Sato, et al. 2008; Wicher, et al. 2008; Sato, et al. 2011; Butterwick, et al. 2018). Insect ORs assemble into heteromeric (probably tetrameric) complexes composed of a tuning OR, which recognises odour ligands, and a universal co-receptor, ORCO, which is critical for complex assembly, subcellular trafficking, and – together with the tuning OR – forms the ion conduction pore (Larsson, et al. 2004; Benton, et al. 2006; Sato, et al. 2008; Wicher, et al. 2008; Butterwick, et al. 2018). Some GRs are also likely to function in multimeric complexes of one or more different subunits (Joseph and Carlson 2015; Scott 2018). A cryogenic electron microscopy (cryo-EM) structure of an ORCO homotetramer (Butterwick, et al. 2018) – which can conduct ions itself upon stimulation with artificial ligands (Jones, et al. 2011) – demonstrated that this receptor adopts a novel fold unrelated to any known family of ion channels. Analysis of amino acid conservation across the OR repertoire (Butterwick, et al. 2018) and *de novo* structure predictions of tuning ORs, guided by patterns of amino acid co-evolution (Hopf, et al. 2015), suggest that this fold is globally similar for all ORs (and potentially GRs).

Surveys for GR-like (GRL) proteins beyond insects have identified members of this family across Protostomia (including in Annelida, Nematoda, and Mollusca) as well as in a limited number of Deuterostomia (including Echinodermata and Hemichordata, but not Chordata) (Robertson 2015; Saina, et al. 2015; Eyun, et al. 2017). Several non-bilaterian animals have recognisable GRLs, including Cnidaria and Placozoa (Nordstrom, et al. 2011; Robertson 2015; Saina, et al. 2015; Eyun, et al. 2017). Although only sparse expression and functional information exists outside Insecta, several lines of evidence indicate that members of this superfamily have roles beyond chemosensation. For example, two homologues in *Caenorhabditis elegans* function in light detection (Edwards, et al. 2008), either as a photoreceptor (Gong, et al. 2016), or through recognition of cellular chemical products produced upon light exposure (Bhatla and Horvitz 2015). Another *C. elegans* homologue functions in motoneurons to control egg-laying (Moresco and Koelle 2004). GRLs in the sea anemone *Nematostella vectensis* and the purple sea urchin*, Strongylocentrotus purpuratus* are expressed early during development (Saina, et al. 2015), and one of the *N. vectensis* proteins may have a role in apical body patterning (Saina, et al. 2015).

Although this family of established (or presumed) ion channels represents one of the largest and most functionally diverse in nature, its evolutionary origin remains unknown. We previously proposed potential homology between GRLs and a family of uncharacterised plant proteins containing the Domain of Unknown Function 3537 (DUF3537), based upon their predicted seven TM domains and intracellular N-termini (Benton 2015). However, this proposal was questioned (Robertson 2019) because DUF3537 proteins lack other features that characterise animal GRLs, such as a motif in TM7 and conserved introns near the 3’ end of the corresponding gene (Robertson 2015; Saina, et al. 2015; Robertson 2019). Moreover, if insect ORs/GRs and plant DUF3537 proteins were derived from a common ancestor, we might expect to find related proteins encoded in the genomes of unicellular eukaryotes. This study aimed to profit from the wealth of genomic information now available to investigate the potential existence of GRL homologues in such species.

## Results

### Screening and assessment of candidate GRLs in unicellular eukaryotes

We used diverse members of the animal GRL and plant DUF3537 families as sequence queries in BLAST searches of protein and genomic sequence databases of unicellular organisms (see Methods). Significant hits were subjected to further assessment to exclude spurious similarities, retaining those that fulfilled most or all of the following criteria: (i) reciprocal BLAST using a candidate sequence as query identified a known GRL or DUF3537 member as a top hit (or no significant similarity to other protein families); (ii) ~350-500 amino acids long, similar to GRLs/DUF3537 proteins; (iii) predicted seven TMDs; (iv) intracellular N-terminus; (v) longer intracellular loops than extracellular loops (a feature of GRLs (Otaki and Yamamoto 2003; Robertson 2015)). These analyses identified 17 sequences from Fungi, Protista and unicellular Plantae (Table 1 and Figure 1), described in more detail below.

**Table 1.**
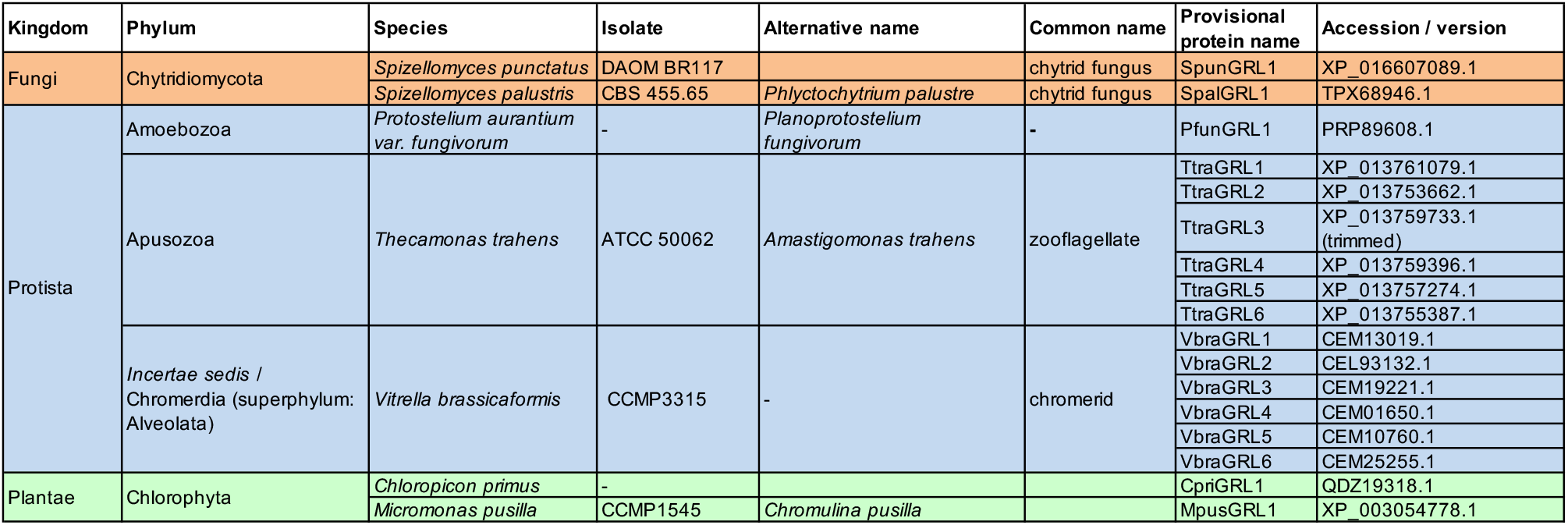
Candidate GRLs in unicellular eukaryotes. Protein sequences are provided in Data S1. Protein nomenclature is provisional and does not imply orthology between species.

| Kingdom | Phylum | Species | Isolate | Alternative name | Common name | Provisional protein name | Accession / version |
| --- | --- | --- | --- | --- | --- | --- | --- |
| Fungi | Chytridiomycota | *Spizellomyces punctatus* | DAOM BR117 |  | chytrid fungus | SpunGRL1 | XP_016607089.1 |
|  |  | *Spizellomyces palustris* | CBS 455.65 | *Phlyctochytrium palustre* | chytrid fungus | SpalGRL1 | TPX68946.1 |
| Protista | Amoebozoa | *Protostelium aurantium* var. *fungivorum* | - | *Planoprotostelium fungivorum* | - | PfunGRL1 | PRP89608.1 |
|  | Apusozoa | *Thecamonas trahens* | ATCC 50062 | *Amastigomonas trahens* | zooflagellate | TtraGRL1 | XP_013761079.1 |
|  |  |  |  |  |  | TtraGRL2 | XP_013753662.1 |
|  |  |  |  |  |  | TtraGRL3 | XP_013759733.1 (trimmed) |
|  |  |  |  |  |  | TtraGRL4 | XP_013759396.1 |
|  |  |  |  |  |  | TtraGRL5 | XP_013757274.1 |
|  |  |  |  |  |  | TtraGRL6 | XP_013755387.1 |
|  | *Incertae sedis* / Chromeridia (superphylum: Alveolata) | *Vitrella brassicaformis* | CCMP3315 | - | chromerid | VbraGRL1 | CEM13019.1 |
|  |  |  |  |  |  | VbraGRL2 | CEL93132.1 |
|  |  |  |  |  |  | VbraGRL3 | CEM19221.1 |
|  |  |  |  |  |  | VbraGRL4 | CEM01650.1 |
|  |  |  |  |  |  | VbraGRL5 | CEM10760.1 |
|  |  |  |  |  |  | VbraGRL6 | CEM25255.1 |
| Plantae | Chlorophyta | *Chloropicon primus* | - |  |  | CpriGRL1 | QDZ19318.1 |
|  |  | *Micromonas pusilla* | CCMP1545 | *Chromulina pusilla* |  | MpusGRL1 | XP_003054778.1 |

**Figure 1.**
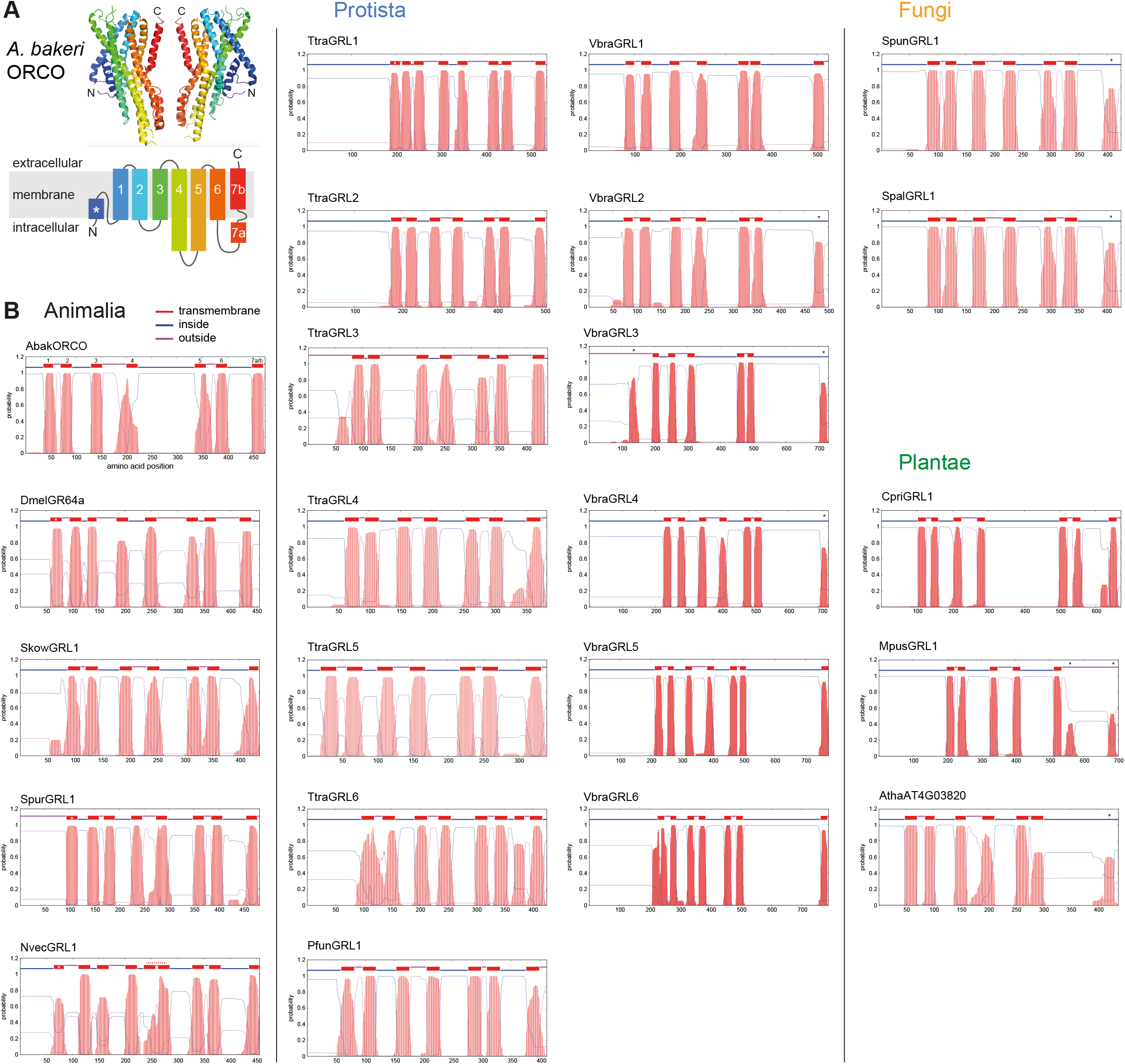
Transmembrane topology predictions of GRLs. (A) Top: cryo-EM structure of *Apocrypta bakeri* ORCO (AbakORCO) (PDB 6C70 (Butterwick, et al. 2018)); only two subunits of the homotetrameric structure are visualised. Bottom: Schematic of the membrane topology of AbakORCO (adapted from (Butterwick, et al. 2018)), coloured as in the cryo-EM structure. The white asterisk marks a helical segment that forms part of a membrane re-entrant loop in the N-terminal region. TM domain 7 is divided into a cytoplasmic segment (7a) and a membrane-spanning segment (7b). (B) TM domain and topology predictions of the previously described and newly recognised GRLs and DUF3537 proteins (Dmel, *Drosophila melanogaster*; Skow, *Saccoglossus kowalevskii*; Spur, *Strongylocentrotus purpuratus*; Nvec, *Nematostella vectensis*; Atha, *Arabidopsis thaliana*; see Table 1 for other species abbreviations and sequence accessions). Each plot represents the posterior probabilities of transmembrane helix and inside/outside cellular location along the protein sequence, adapted from the output of TMHMM Server v2 (Krogh, et al. 2001). In several sequences an extra transmembrane segment near the N-terminus is predicted (marked by a white asterisk in the N-best prediction above the plot); this may represent the re-entrant loop helical region observed in ORCO, rather than a transmembrane region; in at least one case (SpurGRL1) the prediction of this region as a TMD, leads to atypical (and presumably incorrect) N-terminal outside orientation of this protein. Conversely, in a few proteins individual transmembrane domains are not predicted (notably TM7, black asterisks above the N-best plot), which is likely due simply to subthreshold predictions for helices in these regions. In NvecGRL1, the long TM4 helix (which projects into the intracellular space in ORCO (Butterwick, et al. 2018)) is mis-predicated as two TM domains (dashed red line). Similar membrane topology predictions for unicellular species’ GRLs were obtained using TOPCONs (Data S2).

To further assess these candidate GRL homologues, we used their sequences to build and compare Hidden Markov Models (HMMs) using HHblits (Remmert, et al. 2011), a remote homology detection tools that is more sensitive than BLAST (Steinegger, et al. 2019) (see Methods). We constructed HMMs for candidate GRLs, as well as a representative set of animal GRLs and DUF3537 proteins. Each HMM was used as a query to perform all-versus-all alignments. A similarity matrix comparing these alignments was compiled by parsing the probabilities of each alignment (Figure S1). As expected, alignments of HMMs seeded by animal and plant proteins each form clusters of high probability similarity, although we also detected similarity between these clusters, indicative of homology. Importantly, HMMs of all new candidate sequences display significant similarity to those of multiple animal and/or plant proteins. Some candidates clustered more closely with the animal sequences, while others displayed similar probabilities with both animal and plant proteins, an observation consistent with phylogenetic analyses presented below.

We also examined the candidate homologues for the only known primary sequence feature of animal GRLs, a short motif located in the C-terminal half of TM7: (T/S)Yhhhhh(Q/K/E)(F/L/M), where h denotes a hydrophobic amino acid (Robertson 2015). This motif is diagnostic, but not definitive: many insect tuning ORs (as well as some GRs/GRLs) have divergent amino acids at some or all four positions (Scott, et al. 2001; Robertson 2019). Structural and functional analyses of a subset of residues of this motif in ORCO indicates that the TY residues form part of the interaction interface between subunits (Figure 2A), and that their mutation is detrimental to function in some, but not all, combinations of subunits (Nakagawa, et al. 2012; Butterwick, et al. 2018). The terminal L residue of the motif is part of the channel gate (Figure 2A), and its mutation alters ion permeation selectivity (Butterwick, et al. 2018). These observations suggest that divergence from the GRL motif in a given protein could be compatible with it still functioning as an ion channel with different complex assembly and biophysical properties. Bearing these observations in mind, candidate GRLs were inspected for this motif, as described below (Figure 2B). Finally, the corresponding genes were examined for the existence of conserved 3’ introns of the animal genes (Robertson 2015; Saina, et al. 2015); this analysis was ultimately uninformative because of the scarcity of introns in most of these organisms (Roy and Gilbert 2006) (Figure 2B).

**Figure 2.**
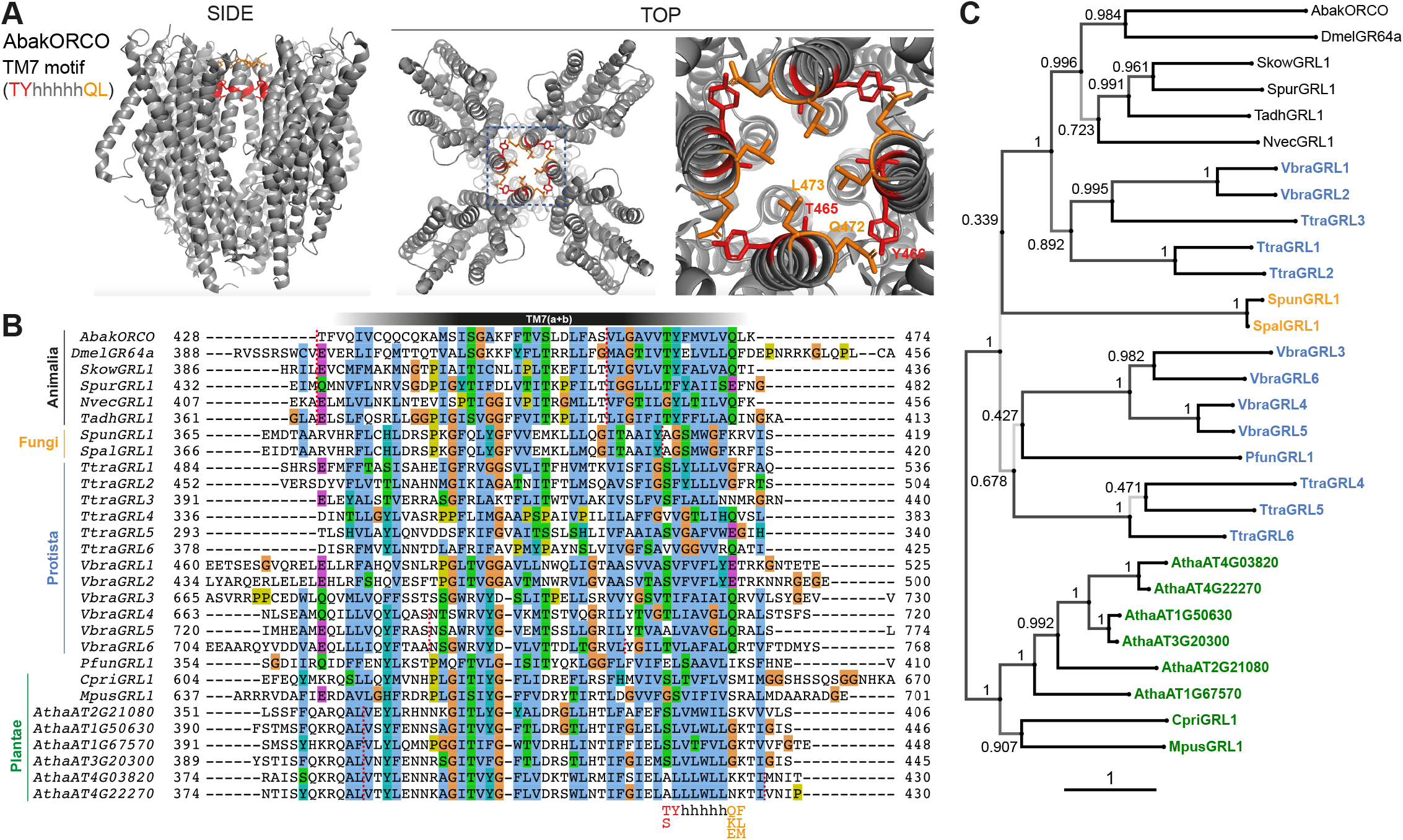
Conservation and divergence in TM7 features and GRL phylogeny. (A) Side and top views of the cryo-EM structure of the ORCO homotetramer (Butterwick, et al. 2018), in which the TM7 motif amino acid side chains are shown in stick format and coloured red or orange. The region in the dashed blue box, representing the extracellular entrance to the ion channel pore, is shown in a magnified view on the far right. (B) Multiple sequence alignment of the C-terminal region (encompassing TM7) of unicellular eukaryotic GRLs and selected animal GRLs and plant DUF3537 proteins. Tadh, *Trichoplax adhaerens*; other species abbreviations are defined in Figure 1 and Table 1. The TM7 motif consensus amino acids (and conservative substitutions) are indicated below the alignment; h indicates a hydrophobic amino acid. Red dashed lines on the alignment indicate positions of predicted introns within the corresponding transcripts. Intron locations are generally conserved within sequences from different Kingdoms, but not between Kingdoms; many Protista sequences do not have introns in this region. (C) Maximum likelihood phylogenetic tree of unicellular eukaryotic GRLs, and selected animal GRLs and plant DUF3537 proteins, with aBayes branch support values. Although the tree is represented as rooted, the rooting is highly uncertain. Protein labels are in black for animals, orange for fungi, blue for protists and green for plants. The scale bar represents one substitution per site.

### Candidate GRLs from unicellular eukaryotes

The fungal kingdom is thought to be the closest relative to Animalia (Baldauf and Palmer 1993). A single candidate GRL was identified in two fungi, *Spizellomyces punctatus* (Russ, et al. 2016) and *Spizellomyces palustris* (van de Vossenberg, et al. 2019). The fungal proteins exhibit the secondary structural features of GRLs (Figure 1), but only one of the four positions of the TM7 motif is conserved (Figure 2B). A 3’ intron is not in the same position of the characteristic last intron of animal GRL genes (Figure 2B). Both of these species are chytrids, an early diverging lineage of fungi that retains some features of the last common opisthokont ancestor of animals and fungi (Medina and Buchler 2020). Chytrids have diverse lifestyles, but are notable for their reproduction via zoospores, which use a motile cilium to swim or crawl.

Taxonomic classification of many single-celled eukaryotes remains unresolved, and we use the term Protista to cover all unicellular species that are neither Fungi nor Plantae. Three such species (of several dozen sequenced protists) were found to encode GRLs. The marine gliding zooflagellate *Thecamonas trahens* (Cavalier-Smith and Chao 2010; Howe, et al. 2020) has six candidate proteins. Beyond secondary structural similarity (Figure 1), the proteins have up to three conserved residues of the TM7 motif (if counting a Y→F substitution in the second position as conservative, as observed in some animal GRLs (e.g., SpurGRL1)) (Figure 2B). The chromerid *Vitrella brassicaformis* (Woo, et al. 2015), a free-living, non-parasitic photosynthetic protist, also has six GRLs, most of which have two conserved positions within the TM7 motif. Finally, the amoebozoan *Protostelium aurantium var. fungivorum* (Hillmann, et al. 2018), has a single GRL. Protosteloid amoeba differ from dictyosteliids by producing simple fruiting bodies with only one or few single stalked spores.

DUF3537 domain proteins are widely (and possibly universally) encoded in higher plant genomes, typically comprising small families of 4-12 members (Benton 2015). Single proteins were also found in unicellular plants (“green algae”), including the marine microalgae *Chloropicon primus* (Lemieux, et al. 2019) and *Micromonas pusilla* (Worden, et al. 2009; van Baren, et al. 2016). As for higher plant sequences, the TM7 motif is largely unrecognisable. The relationship between DUF3537 proteins and the GRL superfamily may have been overlooked earlier because protein alignments are impeded by the longer length of several intracellular loops (IL) of the plant proteins, notably, IL3 (as noted previously (Benton 2015)), in all DUF3537 proteins, and IL2 in the green algal proteins (~200 residues in the *C. primus* homologue). We note that ORCO is also distinguished from other insect ORs and GRs by an additional ~60-70 amino acids in IL2 (Benton, et al. 2006), a region that contributes to channel regulation (Mukunda, et al. 2014; Bahk and Jones 2016).

### Phylogenetic analysis of candidate GRLs

The candidate GRLs are extremely divergent in primary sequence: pairwise alignment of the new proposed family members, together with representative animal and plant proteins, reveal as little as 10% amino acid identity. While this divergence does not preclude their definition as family members – insect OR families themselves have an average of only ~20% amino acid identity (Butterwick, et al. 2018) – it makes it difficult to infer homology from sequence alone, and hinders confident multiprotein alignment. Indeed, an alignment of these 17 sequences, together with selected animal GRLs and plant DUF3537 proteins highlights the absence of any universally conserved residues, beyond the hydrophobic regions predicted to be TM domains (Figure S2). These TM regions are most confidently aligned in the C-terminal half of the protein, with extreme divergence in loop lengths fragmenting the N-terminal half (Figure S2). This pattern of conservation along the protein length is characteristic of insect OR/GR families (Robertson 2015; Saina, et al. 2015; Robertson 2019), which might reflect the role of the N-terminal half in ligand-recognition (and commensurate higher divergence between proteins) and the C-terminal half in mediating subunit interactions and forming the ion channel pore (Butterwick, et al. 2018).

To gain an initial idea of the phylogeny of these proteins, we inferred a maximum likelihood tree (Figure 2C). As expected, the animal and plant/algal proteins form distinct clades. The sets of GRLs of *V. brassicaformis* and *T. trahens* GRLs segregate into two distinct lineages, one of which (comprising VbraGRL1/2 and TtraGRL1/2/3) is more closely related to animal GRLs, while the others are more distantly related; this distinction matches observations from the alignments of HMMs derived from these sequences (Figure S1). Low branch support does not allow for a confident placement of the fungal GRLs; they could group with either of the two protist lineages. All of these observations are consistently held if the tree is inferred from trimmed alignments (Figure S3).

### Common three-dimensional structural predictions of GRLs and DUF3537 proteins

To obtain further evidence supporting the homology of these diverse proteins, we performed *ab initio* structure predictions of animal GRLs (including AbakORCO as control), plant DUF3537 proteins and unicellular eukaryotic GRLs, using the transform-restrained Rosetta (trRosetta) algorithm (Yang, et al. 2020). Most query sequences successfully seeded a multi-sequence alignment to permit extraction of co-evolutionary couplings, generation of inter-residue contact maps and prediction of three-dimensional models (Figure 3A-B, Table S1 and Data S7). The contact maps predicted consistent patterns of anti-parallel packing of TM helices (Figure 3A and Data S7). Concordantly, the top-predicted models bore high similarity to the ORCO cryo-EM structure (Figure 3B and Table S1). As expected, this resemblance was particularly obvious in the transmembrane core of these models (Figure 3B). Furthermore, the highly consistent helical packing of GRLs and DUF3537 proteins is fundamentally different to an unrelated seven TM protein, the Adiponectin Receptor 1, which – despite sharing the same membrane orientation as GRLs – displays an arrangement of helices that is convergent with that of GPCRs (Hopf, et al. 2015; Vasiliauskaite-Brooks, et al. 2017). We confirmed these observations by quantitative comparisons of structures (Figure 3C). Importantly, while protein models of unicellular eukaryotic GRLs were derived from multi-sequence alignments containing large numbers of animal GRLs, the models of the plant proteins used information extracted from alignments of only other DUF3537 family members. This analysis argues that, despite their high sequence divergence, animal and protist GRLs and plant DUF3537 proteins adopt a common three-dimensional architecture.

**Figure 3.**
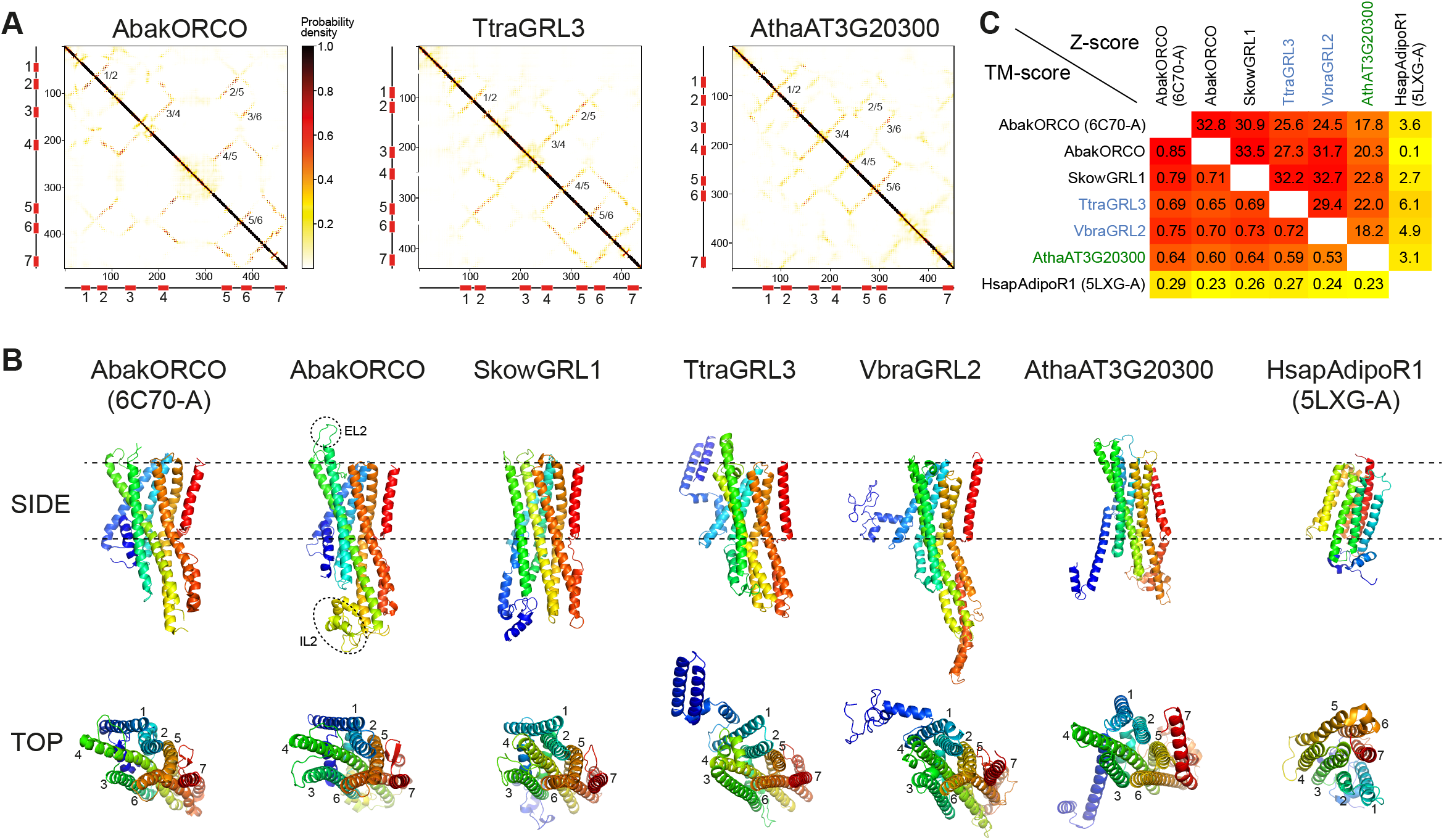
*Ab initio* structural predictions of GRLs and DUF3537 proteins. (A) Inter-residue contact maps from trRosetta analysis of the indicated proteins. The axes represent the indices along the primary sequence; the positions of the predicted TM domains are shown in the schematics. The representation is mirror-symmetric along the diagonal; in one half “lines” of contacts anti-parallel to the diagonal of the map support the existence anti-parallel alpha-helical transmembrane packing arrangements. Most pairs of predicated anti-parallel TMs are conserved across the proteins, despite variation in the length of loops between TM domains, supporting a globally similar packing of TM helices. The complete output of tsRosetta analyses for these and other proteins is provided in Table S1 and Data S7. (B) Side and top views of experimentally-determined (AbakORCO (PDB 6C70 chain A) and *Homo sapiens* Adiponectin Receptor 1 (HsapAdipoR1; PDB 5LXG chain A (Vasiliauskaite-Brooks, et al. 2017)) or the top trRosetta protein model of GRL and DUF3537 proteins. All GRL/DUF3537 proteins have a similar predicted global packing of TM domains (which is particularly evident in the top view in which the seven TM domains are labelled), despite variation in length of the loops and N-terminal regions (coloured in dark blue). By contrast, HsapAdipoR1 has a fundamentally different arrangement of TM domains. The dashed ovals on the AbakORCO model highlight the extracellular loop 2 (EL2) and intracellular loop 2 (IL2) regions that were not visualised in the ORCO cryo-EM structure (Butterwick, et al. 2018). (C) Quantitative pairwise comparisons of the structures shown in (B) using TM-align (Zhang and Skolnick 2005) and DALI (Holm and Rosenstrom 2010)). TM-scores of 0.0-0.30 indicate random structural similarity; TM-scores of 0.5-1.00 indicate that the two proteins adopt generally the same fold. DALI Z-scores of <2 indicate spurious similarity. The two half-matrices are coloured using different scales.

## Discussion

Claims of evolutionary relationships between proteins whose sequence identity resides in the “twilight zone” must be made with caution (Rost 1999). Nevertheless, our primary, secondary and tertiary structural analyses together support the hypothesis that animal ORs/GRs/GRLs, newly-recognised unicellular eukaryotic GRLs, and plant DUF3537 proteins are homologous. The extremely sparse phylogenetic distribution of these genes in unicellular eukaryotes suggests that they have been independently lost in many lineages, although we may have missed distant homologues despite our efforts to perform exhaustive searches. It is also possible that some homologues were acquired by lateral gene transfer, as has been proposed to explain the patchy phylogenetic distribution of microbial rhodopsins (Gavelis, et al. 2017). The global conservation of the structural features of unicellular eukaryotic GRLs and DUF3537 proteins with insect chemosensory receptors argues that they might also be ligand-gated ion channels. It is possible, if not likely, that the proteins fulfil very different physiological roles in these species, as has been suggested in non-Bilateria (Saina, et al. 2015). While functional studies remain a future challenge, the recognition of their existence across Animalia, Plantae, Fungi and Protista provides evidence that this protein family originated in the last common eukaryotic ancestor, 1.5-2 billion years ago (Hedges, et al. 2006). No sequences bearing resemblance to GRLs were found, so far, in Bacteria or Archaea. Future analysis of additional genomic data will help to update the present survey and refine (or refute) the current evolutionary model.

## Materials and Methods

Candidate unicellular GRL sequences were retrieved by searches with PSI-BLAST (Altschul, et al. 1997) and DELTA-BLAST (Boratyn, et al. 2012) using a wide range of animal GRLs (Pfam 7tm_6 (PF02949) and 7tm_7 domain (PF08395)) and plant DUF3537 proteins (PF12056) as queries, retrieving sequences that had a query coverage of >50% and a significant E-value (<0.005). After exclusion of obvious spurious similarities (see criteria in the main text), identified hits were used as queries in further BLAST searches, as well as TBLASTN searches (Gertz, et al. 2006) of the corresponding genomes. Protein topology analysis was performed with TMHMM Server v2.0 (Krogh, et al. 2001) and TOPCONS (Bernsel, et al. 2009). HMMs were constructed with HHblits with default parameters (Remmert, et al. 2011), using three iterations over the Uniclust30 database (Mirdita, et al. 2017; Steinegger and Soding 2017). The results of these iterative searches were examined to verify that no additional candidates GRL sequences had been identified (Data S3). The HMMs resulting from the iterative searches were aligned pairwise using HHblits to obtain a matrix of homology probabilities (code provided in Data S4). The multiprotein alignment was built with MAFFT v7.310 (option “linsi”) (Katoh and Standley 2013), and a maximum likelihood tree was inferred with IQTree v.2.0.6 (Minh, et al. 2020) with aBayes branch support values (Anisimova, et al. 2011). Alignment trimming for Figure S3 was performed with trimAl (option “gappyout”) (Capella-Gutierrez, et al. 2009). Raw untrimmed and trimmed sequence alignments are provided in Data S5-6. Alignments were visualised in Jalview 2.9.0b2 (Waterhouse, et al. 2009) and trees in phylo.io (Robinson, et al. 2016). *Ab initio* protein structure prediction was performed using trRosetta (Yang, et al. 2020), and structural similarities of the resulting models assessed with TM-align (Zhang and Skolnick 2005) and DALI (Holm and Rosenstrom 2010).The ORCO structure (PDB 6C70) (Butterwick, et al. 2018) and models were visualised in MacPyMol.

## Acknowledgements

We thank members of the Benton lab for discussions. Research in R.B.’s laboratory is supported by the University of Lausanne, European Research Council Consolidator and Advanced Grants (615094 and 833548), the Swiss National Science Foundation and the Novartis Foundation for medical-biological Research. D.M. and C.D. were supported by Swiss National Science Foundation Grant 183723.

## Supplementary Information

**Figure S1.**
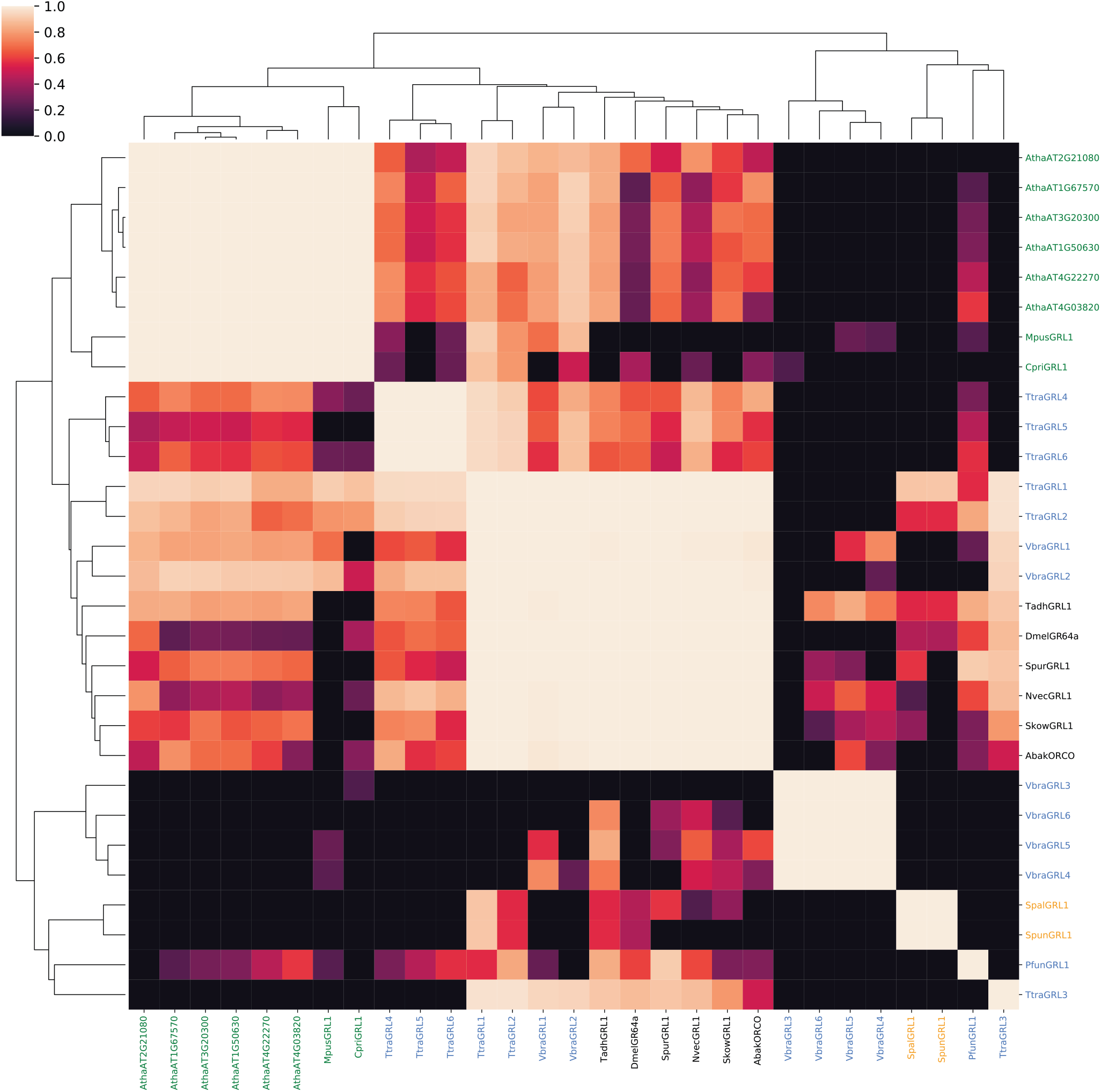
Probabilities of alignments of HMMs of known and candidate GRLs and DUF3537 proteins. A probability similarity matrix representing the quality of pairwise alignments of the HMMs constructed from the protein sequences indicated in the corresponding rows and columns. The similarity matrix was clustered over its rows and columns using UPGMA hierarchical clustering.

**Figure S2.**
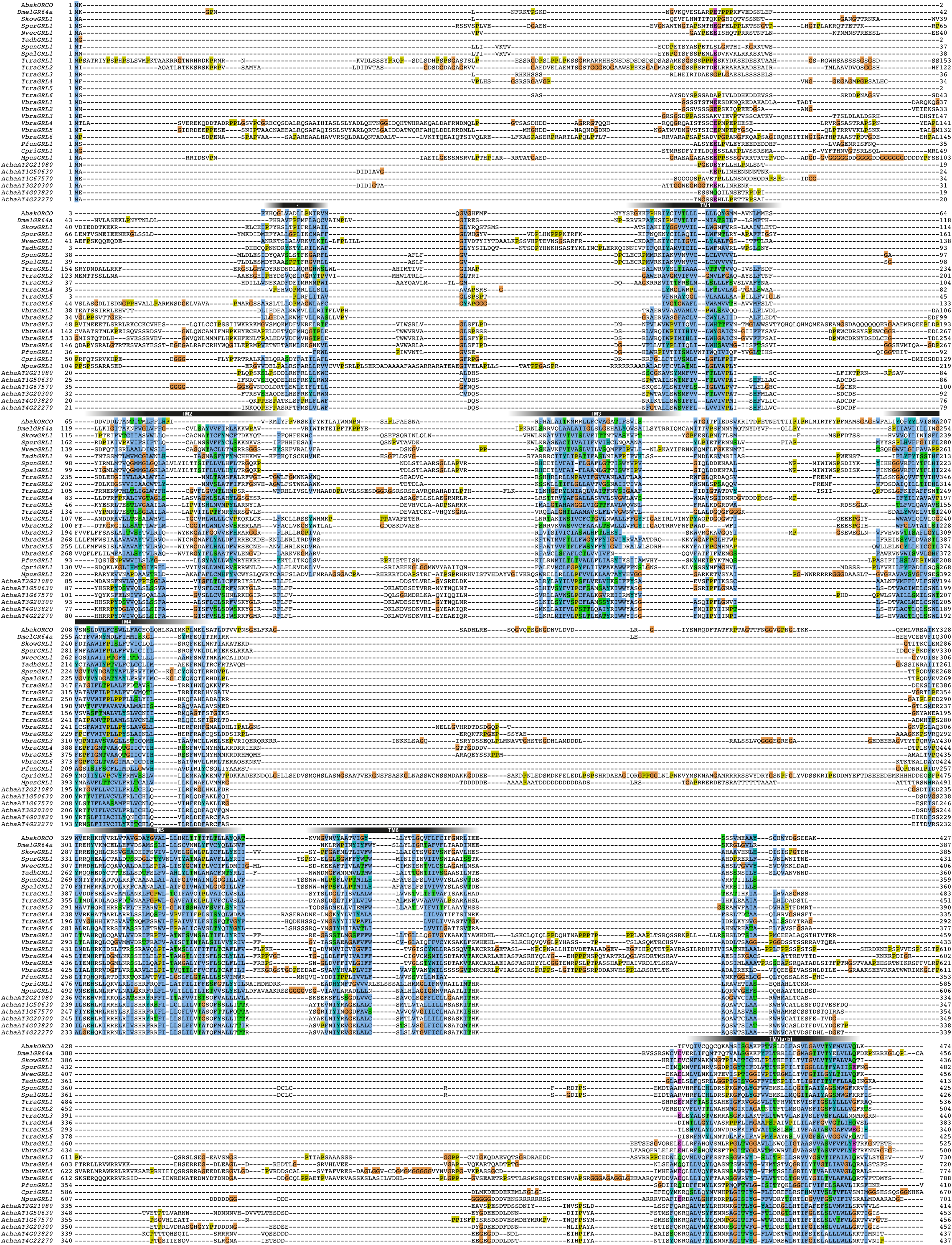
Alignment of GRL superfamily members. Multiple sequence alignment of the GRL proteins illustrated in Figure 1. The approximate positions of the TM domains and the N-terminal re-entrant loop (asterisk) are indicated above the alignment.

**Figure S3.**
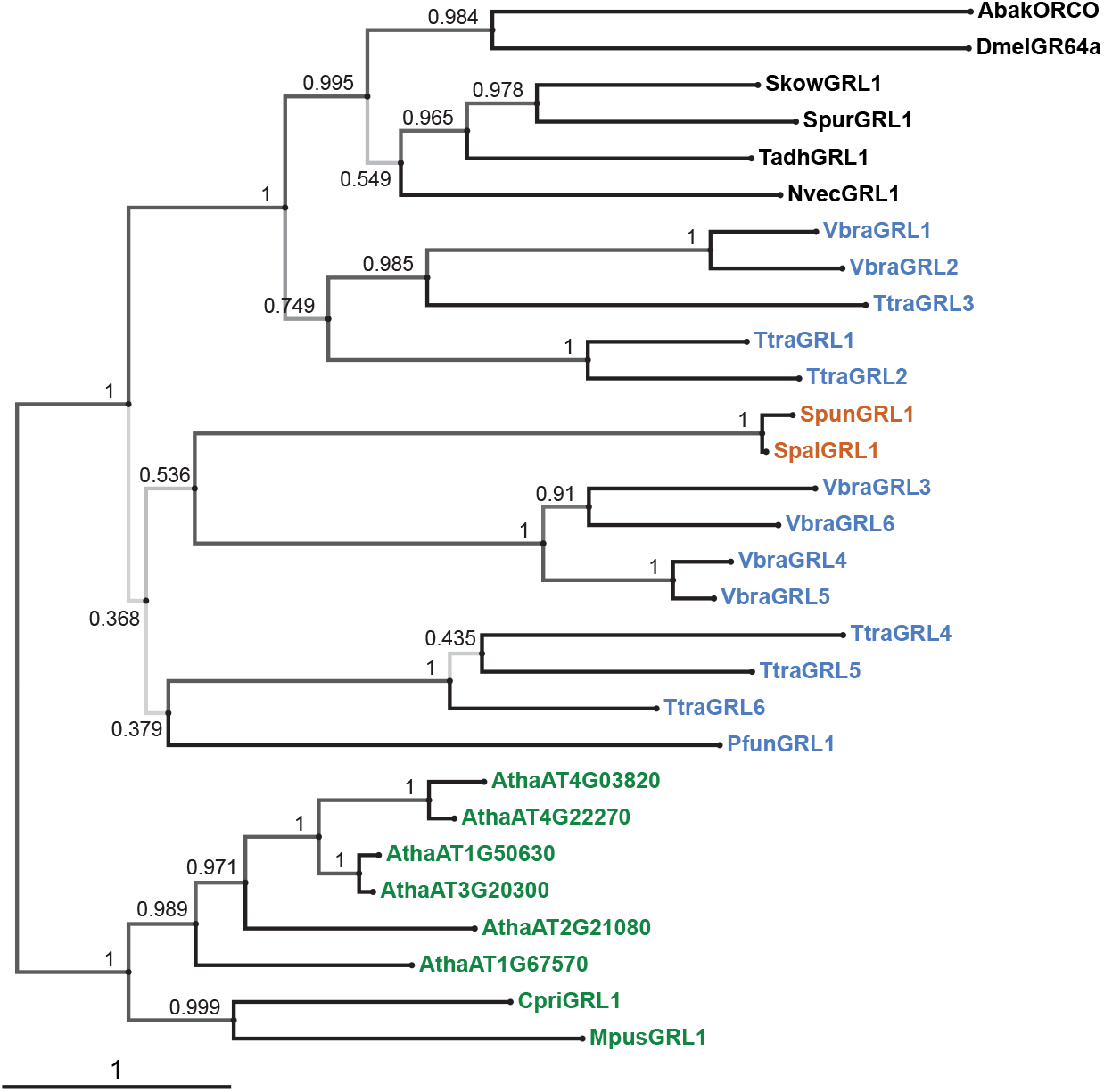
Phylogenetic tree derived from a trimmed multiprotein alignment. Maximum likelihood phylogenetic tree of unicellular eukaryotic GRLs, and selected animal GRLs and plant DUF3537 proteins, with aBayes branch support values. Although the tree is represented as rooted, the rooting is highly uncertain. Protein labels are in black for animals, orange for fungi, blue for protists and green for plants. The scale bar represents one substitution per site.

**Table S1.** *Ab initio* protein modelling of GRLs and DUF3537 proteins. Results from trRosetta analysis with the indicated protein queries. Multiple sequence alignments (MSA) were built automatically with the indicated numbers of proteins. In several cases, insufficient sequences were aligned, leading to inadequate data for prediction of inter-residue contacts. For these proteins, the “estimated TM-scores” of the models (Yang, et al. 2020) are commensurately low (scores < 0.17 are likely to reflect spurious protein structural models (Zhang and Skolnick 2004)) and further analysis was not pursued. For the other proteins, the top hit and corresponding Z-score from Dali searches of the full Protein Data Bank (PDB) with the top-predicted model are shown. For almost all models, individual chains of ORCO homotetramer (A-D) were retrieved as top-hits with a much higher Z-score than the next (non-ORCO) hit. TtraGRL4 and TtraGRL5 models were built using MSAs containing only a subset of plant DUF3537 proteins; although the estimated TM-scores were above the threshold, the retrieved Dali top-hits did not, by contrast, have stand-out Z-scores. Full output of tsRosetta analyses and Dali searches are provided in Data S7.

| Protein | # sequences<br>in MSA | Estimated<br>TM-score | Dali top-hit<br>(full PDB) | Z-score |
| --- | --- | --- | --- | --- |
| AbakORCO | 4209 | 0.641 | ORCO-A | 32.8 |
| DmelGR64a | 1196 | 0.539 | ORCO-D | 26.2 |
| DmelOR22a | 1962 | 0.669 | ORCO-C | 30 |
| SkowGRL1 | 2776 | 0.547 | ORCO-A | 30.9 |
| SpurGRL1 | 646 | 0.495 | ORCO-D | 24.6 |
| NvecGRL1 | 894 | 0.361 | ORCO-C | 18.1 |
| TadhGRL1 | 1598 | 0.485 | ORCO-A | 19.7 |
| SpunGRL1 | 1 | 0.111 | - | - |
| SpalGRL1 | 2 | 0.111 | - | - |
| PfunGRL1 | 1 | 0.1 | - | - |
| TtraGRL1 | 6248 | 0.296 | ORCO-B | 21.7 |
| TtraGRL2 | 2603 | 0.458 | ORCO-A | 24.3 |
| TtraGRL3 | 1239 | 0.48 | ORCO-B | 24.6 |
| TtraGRL4 | 127 | 0.262 | DIABLO (6JX6-D) | 14.8 |
| TtraGRL5 | 222 | 0.242 | PLECTIN (5J1G-A) | 11.5 |
| TtraGRL6 | 312 | 0.167 | - | - |
| VbraGRL1 | 2628 | 0.461 | ORCO-A | 28.7 |
| VbraGRL2 | 3010 | 0.504 | ORCO-A | 25.6 |
| VbraGRL3 | 3 | 0.1 | - | - |
| VbraGRL4 | 2 | 0.1 | - | - |
| VbraGRL5 | 2 | 0.1 | - | - |
| VbraGRL6 | 3 | 0.1 | - | - |
| CpriGRL1 | 171 | 0.106 | - | - |
| MpusGRL1 | 1 | 0.1 | - | - |
| AthaAT2G21080 | 289 | 0.277 | ORCO-B | 17.1 |
| AthaAT4G03820 | 311 | 0.238 | ORCO-C | 11.2 |
| AthaAT4G22270 | 312 | 0.218 | ORCO-C | 13.4 |
| AthaAT1G50630 | 309 | 0.238 | ORCO-B | 16.7 |
| AthaAT1G67570 | 291 | 0.262 | ORCO-C | 17.3 |
| AthaAT3G20300 | 311 | 0.283 | ORCO-B | 18 |

**Data S1. Protein sequences of candidate unicellular eukaryotic GRLs.**

Provisional protein nomenclature (as used in the figures) are indicated in the header of each sequence. Note that these names do not imply orthology between species. Manual corrections to sequences are also noted in the header (e.g., in TtraGRL3 a large C-terminal extension was removed as this is likely due to a mispredicted gene fusion.

**Data S2. TOPCONs analysis output of candidate unicellular eukaryotic GRLs.**

**Data S3. Sequences retrieved through the HMM searches.** Each row contains a separate hit, with identifier, probability, length, query, and score.

**Data S4. Code to generate Figure S1 (matrix of pairwise HMM alignment probabilities).** The Python code is provided as a Jupyter notebook in HTML format, and includes the specific arguments used to run the HHsuite tools.

**Data S5. Multiprotein alignment of GRLs and DUF3537 proteins.**

**Data S6. Trimmed multiprotein alignment of GRLs and DUF3537 proteins.**

**Data S7. tsRosetta analysis output and Dali hits of candidate unicellular eukaryotic GRLs and selected animal GRLs and plant DUF3537 proteins.** Data from these analyses are summarised in Table S1; selected contact maps and models are shown in Figure 3A-B.

*Available as download from:*

https://www.dropbox.com/t/jxQwXqUVGh4Ls4AK
https://www.dropbox.com/t/ujRCuNM9EOJ2ngcW

