## Supplementary material for "A putative origin of insect chemosensory receptors in the last common eukaryotic ancestor": Data S2: nicetop.html

|  |  |
| --- | --- |
|  | 1                                           41 |
| Seq. | MTLLIVKTVE CDPETSYASP ETLSLGRTHI KGRKTWSMLD LESIDYQAVS |
| TOPCONS | iiiiiiiiii iiiiiiiiii iiiiiiiiii iiiiiiiiii iiiiiiiiii |
| OCTOPUS | iiiiiiiiii iiiiiiiiii iiiiiiiiii iiiiiiiiii iiiiiiiiii |
| Philius | iiiiiiiiii iiiiiiiiii iiiiiiiiii iiiiiiiiii iiiiiiiiii |
| PolyPhobius | iiiiiiiiii iiiiiiiiii iiiiiiiiii iiiiiiiiii iiiiiiiiii |
| SCAMPI | iiiiiiiiii iiiiiiiiii iiiiiiiiii iiiiiiiiii iiiiiiiiii |
| SPOCTOPUS | iiiiiiiiii iiiiiiiiii iiiiiiiiii iiiiiiiiii iiiiiiiiii |
| PDB-homology |  |
|  | 51                                          91 |
| Seq. | LSTFKGARFL AFLFGVDPCL ECRPMMRKIA KVVNVVCLCL VVVVLGAYIR |
| TOPCONS | iiiiiiiiii iiiiiiiiii iiiiiiiiii iMMMMMMMMM MMMMMMMMMM |
| OCTOPUS | iiiiiiiiii iiiiiiiiii iiiiiiiiii iMMMMMMMMM MMMMMMMMMM |
| Philius | iiiiiiiiii iiiiiiiiii iiiiiiiiii iMMMMMMMMM MMMMMMMMMM |
| PolyPhobius | iiiiiiiiii iiiiiiiiii iiiiiiiiii iMMMMMMMMM MMMMMMMMMM |
| SCAMPI | iiiiiiiiii iiiiiiiiii iiiiiiiiii iMMMMMMMMM MMMMMMMMMM |
| SPOCTOPUS | iiiiiiiiii iiiiiiiiii iiiiiiiiii iMMMMMMMMM MMMMMMMMMM |
| PDB-homology |  |
|  | 101                                         141 |
| Seq. | MLMTVQGMMG LGQLALVLYI LTTSIFLLVL HFLTRGQKPN DLSTLAARSG |
| TOPCONS | MMoooooooo MMMMMMMMMM MMMMMMMMMM Miiiiiiiii iiiiiiiiii |
| OCTOPUS | MMoooooooo MMMMMMMMMM MMMMMMMMMM Miiiiiiiii iiiiiiiiii |
| Philius | MMoooooooo MMMMMMMMMM MMMMMMMMMM MMMiiiiiii iiiiiiiiii |
| PolyPhobius | MMoooooooo MMMMMMMMMM MMMMMMMMMM MMMiiiiiii iiiiiiiiii |
| SCAMPI | MMoooooooo oooMMMMMMM MMMMMMMMMM MMMMiiiiii iiiiiiiiii |
| SPOCTOPUS | MMoooooooo MMMMMMMMMM MMMMMMMMMM Miiiiiiiii iiiiiiiiii |
| PDB-homology |  |
|  | 151                                         191 |
| Seq. | LLAPVRRHEE TLVFAIFLGA FLGTTISWYV PVGIQLDDEN AALWPMIWIW |
| TOPCONS | iiiiiiiiiM MMMMMMMMMM MMMMMMMMMM oooooooooo oooooooooo |
| OCTOPUS | iiiiiiiiiM MMMMMMMMMM MMMMMMMMMM oooooooooo oooooooooo |
| Philius | iiiiiiiiii MMMMMMMMMM MMMMMMMMMM MMoooooooo oooooooooo |
| PolyPhobius | iiiiiiiiii iMMMMMMMMM MMMMMMMMMM oooooooooo oooooooooo |
| SCAMPI | iiiiiiiiiM MMMMMMMMMM MMMMMMMMMM oooooooooo oooooooooo |
| SPOCTOPUS | iiiiiiiiiM MMMMMMMMMM MMMMMMMMMM oooooooooo oooooooooo |
| PDB-homology |  |
|  | 201                                         241 |
| Seq. | SPSDIYKYIH HGGVRFLYTF LHIVGVTVYD GATYAFLFRV YIMCKGLCYQ |
| TOPCONS | oooooooooo oooooMMMMM MMMMMMMMMM MMMMMMiiii iiiiiiiiii |
| OCTOPUS | oooooooooo oooooMMMMM MMMMMMMMMM MMMMMMiiii iiiiiiiiii |
| Philius | oooooooooo oooooooMMM MMMMMMMMMM MMMMMMMMii iiiiiiiiii |
| PolyPhobius | oooooooooo oooooMMMMM MMMMMMMMMM MMMMMMMMii iiiiiiiiii |
| SCAMPI | oooooooooo oooMMMMMMM MMMMMMMMMM MMMMiiiiii iiiiiiiiii |
| SPOCTOPUS | oooooooooo oooooMMMMM MMMMMMMMMM MMMMMMiiii iiiiiiiiii |
| PDB-homology |  |
|  | 251                                         291 |
| Seq. | WQTLRRDVPL TTPQDVEEFM TYFRKADTQL RKFCAANALA IAIFGIVHAI |
| TOPCONS | iiiiiiiiii iiiiiiiiii iiiiiiiiii iiMMMMMMMM MMMMMMMMMM |
| OCTOPUS | iiiiiiiiii iiiiiiiiii iiiiiiiiii iiiiMMMMMM MMMMMMMMMM |
| Philius | iiiiiiiiii iiiiiiiiii iiiiiiiiii iiMMMMMMMM MMMMMMMMMM |
| PolyPhobius | iiiiiiiiii iiiiiiiiii iiiiiiiiii iiMMMMMMMM MMMMMMMMMM |
| SCAMPI | iiiiiiiiii iiiiiiiiii iiiiiiiiii iiMMMMMMMM MMMMMMMMMM |
| SPOCTOPUS | iiiiiiiiii iiiiiiiiii iiiiiiiiii iiiiMMMMMM MMMMMMMMMM |
| PDB-homology |  |
|  | 301                                         341 |
| Seq. | NLGDGILLVF TSSNWNLPSF LVPTMILHAF ATSLFSLVIV VYSLASVTDE |
| TOPCONS | MMMooooooo oooooooooo ooooMMMMMM MMMMMMMMMM MMMMMiiiii |
| OCTOPUS | MMMMMooooo oooooooooo oooooMMMMM MMMMMMMMMM MMMMMMiiii |
| Philius | MMoooooooo oooooooooo oooMMMMMMM MMMMMMMMMM MMMMMiiiii |
| PolyPhobius | MMoooooooo oooooooooo MMMMMMMMMM MMMMMMMMMM MMMMMiiiii |
| SCAMPI | MMMooooooo oooooooooo ooooMMMMMM MMMMMMMMMM MMMMMiiiii |
| SPOCTOPUS | MMMMMooooo oooooooooo oooooMMMMM MMMMMMMMMM MMMMMMiiii |
| PDB-homology |  |
|  | 351                                         391 |
| Seq. | VRRTIILLSD CLCRFRDTPS EMDTAARVHR FLCHLDRSPK GFQLYGFVVE |
| TOPCONS | iiiiiiiiii iiiiiiiiii iiiiiiiiii iiiiiiiiii iiiiMMMMMM |
| OCTOPUS | iiiiiiiiii iiiiiiiiii iiiiiiiiii iiiiiiiiii iiiiMMMMMM |
| Philius | iiiiiiiiii iiiiiiiiii iiiiiiiiii iiiiiiiiii iiiiiiiiii |
| PolyPhobius | iiiiiiiiii iiiiiiiiii iiiiiiiiii iiiiiiiiii iiiiMMMMMM |
| SCAMPI | iiiiiiiiii iiiiiiiiii iiiiiiiiii iiiiiiiiii iiiiiiiiii |
| SPOCTOPUS | iiiiiiiiii iiiiiiiiii iiiiiiiiii iiiiiiiiii iiiiMMMMMM |
| PDB-homology |  |

|  |  |
| --- | --- |
|  | 401                   421 |
| Seq. | MKLLLQGITA AIYAGSMWGF KRVIS |
| TOPCONS | MMMMMMMMMM MMMMMooooo ooooo |
| OCTOPUS | MMMMMMMMMM MMMMMooooo ooooo |
| Philius | iiMMMMMMMM MMMMMMMooo ooooo |
| PolyPhobius | MMMMMMMMMM MMMMMMMMMo ooooo |
| SCAMPI | iiMMMMMMMM MMMMMMMMMM MMMoo |
| SPOCTOPUS | MMMMMMMMMM MMMMMooooo ooooo |
| PDB-homology |  |
