## Supplementary material for "A putative origin of insect chemosensory receptors in the last common eukaryotic ancestor": Data S2: nicetop.html

|  |  |
| --- | --- |
|  | 1                                           41 |
| Seq. | MNLTIVRTVE YNPGTSFSSP ENLEDVKVKL RKKSKTWSTL DLESMDYRAA |
| TOPCONS | iiiiiiiiii iiiiiiiiii iiiiiiiiii iiiiiiiiii iiiiiiiiii |
| OCTOPUS | iiiiiiiiii iiiiiiiiii iiiiiiiiii iiiiiiiiii iiiiiiiiii |
| Philius | iiiiiiiiii iiiiiiiiii iiiiiiiiii iiiiiiiiii iiiiiiiiii |
| PolyPhobius | iiiiiiiiii iiiiiiiiii iiiiiiiiii iiiiiiiiii iiiiiiiiii |
| SCAMPI | iiiiiiiiii iiiiiiiiii iiiiiiiiii iiiiiiiiii iiiiiiiiii |
| SPOCTOPUS | iiiiiiiiii iiiiiiiiii iiiiiiiiii iiiiiiiiii iiiiiiiiii |
| PDB-homology |  |
|  | 51                                          91 |
| Seq. | SPPTFRGVRL LASLFGVDPC LECRPMVRKI AKVVNVVCLC MVVVVLSGYI |
| TOPCONS | iiiiiiiiii iiiiiiiiii iiiiiiiiii iiMMMMMMMM MMMMMMMMMM |
| OCTOPUS | iiiiiiiiii iiiiiiiiii iiiiiiiiii iiMMMMMMMM MMMMMMMMMM |
| Philius | iiiiiiiiii iiiiiiiiii iiiiiiiiii iiMMMMMMMM MMMMMMMMMM |
| PolyPhobius | iiiiiiiiii iiiiiiiiii iiiiiiiiii iiMMMMMMMM MMMMMMMMMM |
| SCAMPI | iiiiiiiiii iiiiiiiiii iiiiiiiiii iiMMMMMMMM MMMMMMMMMM |
| SPOCTOPUS | iiiiiiiiii iiiiiiiiii iiiiiiiiii iiMMMMMMMM MMMMMMMMMM |
| PDB-homology |  |
|  | 101                                         141 |
| Seq. | GMLMTVQGMM GLGKLALVLY ILTTSIFLLV LHYLTRGQKP TDLSTLAARS |
| TOPCONS | MMMooooooo oMMMMMMMMM MMMMMMMMMM MMiiiiiiii iiiiiiiiii |
| OCTOPUS | MMMooooooo MMMMMMMMMM MMMMMMMMMM Miiiiiiiii iiiiiiiiii |
| Philius | MMMooooooo oMMMMMMMMM MMMMMMMMMM MMMMiiiiii iiiiiiiiii |
| PolyPhobius | MMMooooooo oooMMMMMMM MMMMMMMMMM MMMMiiiiii iiiiiiiiii |
| SCAMPI | MMMooooooo ooooMMMMMM MMMMMMMMMM MMMMMiiiii iiiiiiiiii |
| SPOCTOPUS | MMMooooooo MMMMMMMMMM MMMMMMMMMM Miiiiiiiii iiiiiiiiii |
| PDB-homology |  |
|  | 151                                         191 |
| Seq. | GLLAPVRRHE ETLVFAIFLA AMLGTTISWY IPVGIQLDDE NTALWPMIWI |
| TOPCONS | iiiiiiiiii MMMMMMMMMM MMMMMMMMMM Mooooooooo oooooooooo |
| OCTOPUS | iiiiiiiiii MMMMMMMMMM MMMMMMMMMM Mooooooooo oooooooooo |
| Philius | iiiiiiiiii iMMMMMMMMM MMMMMMMMMM MMMooooooo oooooooooo |
| PolyPhobius | iiiiiiiiii iiMMMMMMMM MMMMMMMMMM Mooooooooo oooooooooo |
| SCAMPI | iiiiiiiiii MMMMMMMMMM MMMMMMMMMM Mooooooooo oooooooooo |
| SPOCTOPUS | iiiiiiiiii MMMMMMMMMM MMMMMMMMMM Mooooooooo oooooooooo |
| PDB-homology |  |
|  | 201                                         241 |
| Seq. | WSPSDISKFI NNGGVRFFYT FLHIVGVTVY DGATYAYLFR VYIMCKGLCY |
| TOPCONS | oooooooooo ooooooMMMM MMMMMMMMMM MMMMMMMiii iiiiiiiiii |
| OCTOPUS | oooooooooo ooooooMMMM MMMMMMMMMM MMMMMMMiii iiiiiiiiii |
| Philius | oooooooooo ooooooooMM MMMMMMMMMM MMMMMMMMMi iiiiiiiiii |
| PolyPhobius | oooooooooo ooooooMMMM MMMMMMMMMM MMMMMMMMii iiiiiiiiii |
| SCAMPI | oooooooooo ooooMMMMMM MMMMMMMMMM MMMMMiiiii iiiiiiiiii |
| SPOCTOPUS | oooooooooo oooMMMMMMM MMMMMMMMMM MMMMMMMMMM MMMMiiiiii |
| PDB-homology |  |
|  | 251                                         291 |
| Seq. | QWQTLRHDLP LTTPQDVEVF MTHFRKADTQ LRKFCAANAL AIAIFGIVHA |
| TOPCONS | iiiiiiiiii iiiiiiiiii iiiiiiiiii iiiMMMMMMM MMMMMMMMMM |
| OCTOPUS | iiiiiiiiii iiiiiiiiii iiiiiiiiii iiiiiMMMMM MMMMMMMMMM |
| Philius | iiiiiiiiii iiiiiiiiii iiiiiiiiii iiiMMMMMMM MMMMMMMMMM |
| PolyPhobius | iiiiiiiiii iiiiiiiiii iiiiiiiiii iiiMMMMMMM MMMMMMMMMM |
| SCAMPI | iiiiiiiiii iiiiiiiiii iiiiiiiiii iiiMMMMMMM MMMMMMMMMM |
| SPOCTOPUS | iiiiiiiiii iiiiiiiiii iiiiiiiiii iiiiiMMMMM MMMMMMMMMM |
| PDB-homology |  |
|  | 301                                         341 |
| Seq. | INLGDGILLV FTSSNWNFPS FLLPTMILHA FATSLFSIVI VVYSLASVTD |
| TOPCONS | MMMMoooooo oooooooooo oooooMMMMM MMMMMMMMMM MMMMMMiiii |
| OCTOPUS | MMMMMMoooo oooooooooo ooooooMMMM MMMMMMMMMM MMMMMMMiii |
| Philius | MMMooooooo oooooooooo ooooMMMMMM MMMMMMMMMM MMMMMMiiii |
| PolyPhobius | MMMooooooo oooooooooo oMMMMMMMMM MMMMMMMMMM MMMMMMiiii |
| SCAMPI | MMMMoooooo oooooooooo oooooMMMMM MMMMMMMMMM MMMMMMiiii |
| SPOCTOPUS | MMMMMMoooo oooooooooo ooooooMMMM MMMMMMMMMM MMMMMMMiii |
| PDB-homology |  |
|  | 351                                         391 |
| Seq. | EVRRSIILLS DCLCRSGDIP SEIDTAARVH RFLCHLDRSP KGFQLYGFVV |
| TOPCONS | iiiiiiiiii iiiiiiiiii iiiiiiiiii iiiiiiiiii iiiiiMMMMM |
| OCTOPUS | iiiiiiiiii iiiiiiiiii iiiiiiiiii iiiiiiiiii iiiiiMMMMM |
| Philius | iiiiiiiiii iiiiiiiiii iiiiiiiiii iiiiiiiiii iiiiiiiiii |
| PolyPhobius | iiiiiiiiii iiiiiiiiii iiiiiiiiii iiiiiiiiii iiiiiMMMMM |
| SCAMPI | iiiiiiiiii iiiiiiiiii iiiiiiiiii iiiiiiiiii iiiiiiiiii |
| SPOCTOPUS | iiiiiiiiii iiiiiiiiii iiiiiiiiii iiiiiiiiii iiiiiMMMMM |
| PDB-homology |  |

|  |  |
| --- | --- |
|  | 401                   421 |
| Seq. | EMKLLMQGIT AAIYAGSMWG FKRFIS |
| TOPCONS | MMMMMMMMMM MMMMMMoooo oooooo |
| OCTOPUS | MMMMMMMMMM MMMMMMoooo oooooo |
| Philius | iiiMMMMMMM MMMMMMMMoo oooooo |
| PolyPhobius | MMMMMMMMMM MMMMMMMMMM oooooo |
| SCAMPI | iiiMMMMMMM MMMMMMMMMM MMMMoo |
| SPOCTOPUS | MMMMMMMMMM MMMMMMoooo oooooo |
| PDB-homology |  |
