## Supplementary material for "A putative origin of insect chemosensory receptors in the last common eukaryotic ancestor": Data S2: nicetop.html

|  |  |
| --- | --- |
|  | 1                                           41 |
| Seq. | MPSATRIYPS PHPSLSVMPK TAAKRRGTNR HRDKPRNRNK VDLSSSYPRQ |
| TOPCONS | iiiiiiiiii iiiiiiiiii iiiiiiiiii iiiiiiiiii iiiiiiiiii |
| OCTOPUS | iiiiiiiiii iiiiiiiiii iiiiiiiiii iiiiiiiiii iiiiiiiiii |
| Philius | iiiiiiiiii iiiiiiiiii iiiiiiiiii iiiiiiiiii iiiiiiiiii |
| PolyPhobius | oooooooooo oooooooooo oooooooooo oooooooooo oooooooooo |
| SCAMPI | oooooooooo oooooooooo oooooooooo oooooooooo oooooooooo |
| SPOCTOPUS | iiiiiiiiii iiiiiiiiii iiiiiiiiii iiiiiiiiii iiiiiiiiii |
| PDB-homology |  |
|  | 51                                          91 |
| Seq. | PSDLSDHPSP SGASTSLPES SRGDPSLPPL PKSSGRRARR HHSNSDSDSD |
| TOPCONS | iiiiiiiiii iiiiiiiiii iiiiiiiiii iiiiiiiiii iiiiiiiiii |
| OCTOPUS | iiiiiiiiii iiiiiiiiii iiiiiiiiii iiiiiiiiii iiiiiiiiii |
| Philius | iiiiiiiiii iiiiiiiiii iiiiiiiiii iiiiiiiiii iiiiiiiiii |
| PolyPhobius | oooooooooo oooooooooo oooooooooo oooooooooo oooooooooo |
| SCAMPI | oooooooooo oooooooooo oooooooooo oooooooooo oooooooooo |
| SPOCTOPUS | iiiiiiiiii iiiiiiiiii iiiiiiiiii iiiiiiiiii iiiiiiiiii |
| PDB-homology |  |
|  | 101                                         141 |
| Seq. | SDSDSASAME SGSSSPPPES KYDKSEDESK TAAHGSRQWH SASSSSGSGS |
| TOPCONS | iiiiiiiiii iiiiiiiiii iiiiiiiiii iiiiiiiiii iiiiiiiiii |
| OCTOPUS | iiiiiiiiii iiiiiiiiii iiiiiiiiii iiiiiiiiii iiiiiiiiii |
| Philius | iiiiiiiiii iiiiiiiiii iiiiiiiiii iiiiiiiiii iiiiiiiiii |
| PolyPhobius | oooooooooo oooooooooo oooooooooo oooooooooo oooooooooo |
| SCAMPI | oooooooooo oooooooooo oooooooooo oooooooooo oooooooooo |
| SPOCTOPUS | iiiiiiiiii iiiiiiiiii iiiiiiiiii iiiiiiiiii iiiiiiiiii |
| PDB-homology |  |
|  | 151                                         191 |
| Seq. | DSSSRYDNDA LLRKFERGSL GMVDYRNDND LMQRGHWSLW LAHIMTIVFG |
| TOPCONS | iiiiiiiiii iiiiiiiiii iiiiiiiiii iiiiiiiiii iiiiiiiiii |
| OCTOPUS | iiiiiiiiii iiiiiiiiii iiiiiiiiii iiiiiiiiii iiiiiiiiii |
| Philius | iiiiiiiiii iiiiiiiiii iiiiiiiiii iiiiiMMMMM MMMMMMMMMM |
| PolyPhobius | oooooooooo oooooooooo oooooooooo oooooMMMMM MMMMMMMMMM |
| SCAMPI | oooooooooo oooooooooo oooooooooo ooooooMMMM MMMMMMMMMM |
| SPOCTOPUS | iiiiiiiiii iiiiiiiiii iiiiiiiiii iiiiiiiiii iiiiiiiiii |
| PDB-homology |  |
|  | 201                                         241 |
| Seq. | INAPSALWRV YSLTIAAAVT TVTVVQIVSL FSDFSDLEHG IIVLLAALWY |
| TOPCONS | iiiiiiiiiM MMMMMMMMMM MMMMMMMMMM oooooooooM MMMMMMMMMM |
| OCTOPUS | iiiiiiiiMM MMMMMMMMMM MMMMMMMMMo ooooooooMM MMMMMMMMMM |
| Philius | MMMMMMMooo MMMMMMMMMM MMMMMMMMMM MMiiiiiiiM MMMMMMMMMM |
| PolyPhobius | MMMMiiiiiM MMMMMMMMMM MMMMMMMMMM Mooooooooo MMMMMMMMMM |
| SCAMPI | MMMMMMMiiM MMMMMMMMMM MMMMMMMMMM oooooooooo oooMMMMMMM |
| SPOCTOPUS | iiiiiiiiMM MMMMMMMMMM MMMMMMMMMo ooooooooMM MMMMMMMMMM |
| PDB-homology |  |
|  | 251                                         291 |
| Seq. | LLNMLTAWSF ALRFWGHTGW LFGMWKAMWQ SEADVCRSHH RLRLLMPAVI |
| TOPCONS | MMMMMMMMMM iiiiiiiiii iiiiiiiiii iiiiiiiiii iMMMMMMMMM |
| OCTOPUS | MMMMMMMMMi iiiiiiiiii iiiiiiiiii iiiiiiiiiM MMMMMMMMMM |
| Philius | MMMMMMMoMM MMMMMMMMMM MMMiiiiiii iiiiiiiiii iiiMMMMMMM |
| PolyPhobius | MMMMMMMMMM MMMMMMiiii iiiiiiiiii iiiiiiiiii iiiMMMMMMM |
| SCAMPI | MMMMMMMMMM MMMMiiiiii iiiiiiiiii iiiiiiiiii iMMMMMMMMM |
| SPOCTOPUS | MMMMMMMMMi iiiiiiiiii iiiiiiiiii iiiiiiiiiM MMMMMMMMMM |
| PDB-homology |  |
|  | 301                                         341 |
| Seq. | FGVVANLALT LAARWFGLQR DAQSVFREMF VLSSNGAQVV FTVIHVFATG |
| TOPCONS | MMMMMMMMMM MMoooooooo oooooooooo oooooooooM MMMMMMMMMM |
| OCTOPUS | MMMMMMMMMM oooooooooo oooooooooo oooooooooM MMMMMMMMMM |
| Philius | MMMMMMMMMM MMMMMMoooo oooooooooo ooooooooMM MMMMMMMMMM |
| PolyPhobius | MMMMMMMMMM MMMMMMMooo oooooooooo ooooooMMMM MMMMMMMMMM |
| SCAMPI | MMMMMMMMMM MMoooooooo oooooooooo oooooooooo ooooooMMMM |
| SPOCTOPUS | MMMMMMMMMM oooooooooo oooooooooo oooooooooM MMMMMMMMMM |
| PDB-homology |  |
|  | 351                                         391 |
| Seq. | IFLTPLALFF DTAVSLTRRI HWLQKKVFRD EKSLTELVDD FSELSVHAML |
| TOPCONS | MMMMMMMMMM iiiiiiiiii iiiiiiiiii iiiiiiiiii iiiiiiiiii |
| OCTOPUS | MMMMMMMMMM iiiiiiiiii iiiiiiiiii iiiiiiiiii iiiiiiiiii |
| Philius | MMMMMMMMMM MMMiiiiiii iiiiiiiiii iiiiiiiiii iiiiiiiiii |
| PolyPhobius | MMMMMMMMMM iiiiiiiiii iiiiiiiiii iiiiiiiiii iiiiiiiiii |
| SCAMPI | MMMMMMMMMM MMMMMMMiii iiiiiiiiii iiiiiiiiii iiiiiiiiii |
| SPOCTOPUS | MMMMMMMMMM iiiiiiiiii iiiiiiiiii iiiiiiiiii iiiiiiiiii |
| PDB-homology |  |
|  | 401                                         441 |
| Seq. | ANKLFGPWLT CIFAVQIPLV AICLVSLLSY TSLDGLTISV LALWLVVNTV |
| TOPCONS | iiiiiiMMMM MMMMMMMMMM MMMMMMMooo oooooooMMM MMMMMMMMMM |
| OCTOPUS | iiiiiiMMMM MMMMMMMMMM MMMMMMMooo oooooooMMM MMMMMMMMMM |
| Philius | iiiiMMMMMM MMMMMMMMMM MMMMMMMMMM oooooooooM MMMMMMMMMM |
| PolyPhobius | iiiiMMMMMM MMMMMMMMMM MMMMMMMMMM oooooMMMMM MMMMMMMMMM |
| SCAMPI | iiiiiiiMMM MMMMMMMMMM MMMMMMMMoo oooooooooM MMMMMMMMMM |
| SPOCTOPUS | iiiiiiMMMM MMMMMMMMMM MMMMMMMooo oooooooMMM MMMMMMMMMM |
| PDB-homology |  |
|  | 451                                         491 |
| Seq. | LTFTVAFISA KLHADTEATI HKIALHVASG RSSSHRSEFM FFTASISAHE |
| TOPCONS | MMMMMMMMii iiiiiiiiii iiiiiiiiii iiiiiiiiii iiMMMMMMMM |
| OCTOPUS | MMMMMMMMii iiiiiiiiii iiiiiiiiii iiiiiiiiii iiiiiiiiii |
| Philius | MMMMMMMMMM iiiiiiiiii iiiiiiiiii iiiiiiiiMM MMMMMMMMMM |
| PolyPhobius | MMMMMMMMMM iiiiiiiiii iiiiiiiiii iiiiiiiiMM MMMMMMMMMM |
| SCAMPI | MMMMMMMMMM iiiiiiiiii iiiiiiiiii iiiiiiiiii iiiiiiiiii |
| SPOCTOPUS | MMMMMMMMii iiiiiiiiii iiiiiiiiii iiiiiiiiii iiiiiiiiii |
| PDB-homology |  |

|  |  |
| --- | --- |
|  | 501                              531 |
| Seq. | IGFRVGGSVL ITFHVMTKVI SFIGSLYLLL VGFRAQ |
| TOPCONS | MMMMMMMMMM MMMoMMMMMM MMMMMMMMMM MMMMMi |
| OCTOPUS | iiiiMMMMMM MMMMMMMMMo MMMMMMMMMM MMMMMi |
| Philius | MMMooooooM MMMMMMMMMM MMMMMMMMMM MMMiii |
| PolyPhobius | MMMMMoooMM MMMMMMMMMM MMMMMMMMMM MMMiii |
| SCAMPI | iiiiiiiiii iiiiMMMMMM MMMMMMMMMM MMMMMo |
| SPOCTOPUS | iiiiMMMMMM MMMMMMMMMM MMMMMMMMMM MMMMMo |
| PDB-homology |  |
