## Supplementary material for "A putative origin of insect chemosensory receptors in the last common eukaryotic ancestor": Data S2: nicetop.html

|  |  |
| --- | --- |
|  | 1                                           41 |
| Seq. | MIAQATLRTK KSRSKPRPVS AMYALDIDVT ASGDSDGDAG AGRVVGAEDV |
| TOPCONS | iiiiiiiiii iiiiiiiiii iiiiiiiiii iiiiiiiiii iiiiiiiiii |
| OCTOPUS | iiiiiiiiii iiiiiiiiii iiiiiiiiii iiiiiiiiii iiiiiiiiii |
| Philius | iiiiiiiiii iiiiiiiiii iiiiiiiiii iiiiiiiiii iiiiiiiiii |
| PolyPhobius | iiiiiiiiii iiiiiiiiii iiiiiiiiii iiiiiiiiii iiiiiiiiii |
| SCAMPI | oooooooooo oooooooooo oooooooooo oooooooooo oooooooooo |
| SPOCTOPUS | iiiiiiiiii iiiiiiiiii iiiiiiiiii iiiiiiiiii iiiiiiiiii |
| PDB-homology |  |
|  | 51                                          91 |
| Seq. | ELAEMTSGST GGGEQGAAWS PEKSGAGMAS PQSGSSPSRT DELRRARARA |
| TOPCONS | iiiiiiiiii iiiiiiiiii iiiiiiiiii iiiiiiiiii iiiiiiiiii |
| OCTOPUS | iiiiiiiiii iiiiiiiiii iiiiiiiiii iiiiiiiiii iiiiiiiiii |
| Philius | iiiiiiiiii iiiiiiiiii iiiiiiiiii iiiiiiiiii iiiiiiiiii |
| PolyPhobius | iiiiiiiiii iiiiiiiiii iiiiiiiiii iiiiiiiiii iiiiiiiiii |
| SCAMPI | oooooooooo oooooooooo oooooooooo oooooooooo oooooooooo |
| SPOCTOPUS | iiiiiiiiii iiiiiiiiii iiiiiiiiii iiiiiiiiii iiiiiiiiii |
| PDB-homology |  |
|  | 101                                         141 |
| Seq. | DSEAIRTMLA AVDVQSGGSH HFMEMTTSSL LNAAAEEGHI PHYDSVQSLR |
| TOPCONS | iiiiiiiiii iiiiiiiiii iiiiiiiiii iiiiiiiiii iiiiiiiiii |
| OCTOPUS | iiiiiiiiii iiiiiiiiii iiiiiiiiii iiiiiiiiii iiiiiiiiii |
| Philius | iiiiiiiiii iiiiiiiiii iiiiiiiiii iiiiiiiiii iiiiiiiiii |
| PolyPhobius | iiiiiiiiii iiiiiiiiii iiiiiiiiii iiiiiiiiii iiiiiiiiii |
| SCAMPI | oooooooooo oooooooooo oooooooooo oooooooooo oooooooooo |
| SPOCTOPUS | iiiiiiiiii iiiiiiiiii iiiiiiiiii iiiiiiiiii iiiiiiiiii |
| PDB-homology |  |
|  | 151                                         191 |
| Seq. | GRYTPVVIMH WLTRLLGLTR IGAWERAVGM VIFVVGMVLF GAQAYSLFTN |
| TOPCONS | iiiiiiiiii iiiiiiiiii iiiiiiMMMM MMMMMMMMMM MMMMMMMooo |
| OCTOPUS | iiiiiiiiii iiiiiiiiii iiiiiMMMMM MMMMMMMMMM MMMMMMoooo |
| Philius | iiiiiiiiii iiiiiiiiii iiiiiiMMMM MMMMMMMMMM MMMMMMMooo |
| PolyPhobius | iiiiiiiiii iiiiiiiiii iiiiiiMMMM MMMMMMMMMM MMMMMMMMoo |
| SCAMPI | oooMMMMMMM MMMMMMMMMM MMMMiiMMMM MMMMMMMMMM MMMMMMMooo |
| SPOCTOPUS | iiiiiiiiii iiiiiiiiii iiiiiMMMMM MMMMMMMMMM MMMMMMoooo |
| PDB-homology |  |
|  | 201                                         241 |
| Seq. | FTDLKHGSVV IIAACWTLYN MVSLATFIFR FGNVHAGWLF STWKAAWKTE |
| TOPCONS | ooooooooMM MMMMMMMMMM MMMMMMMMMi iiiiiiiiii iiiiiiiiii |
| OCTOPUS | ooooooMMMM MMMMMMMMMM MMMMMMMiii iiiiiiiiii iiiiiiiiii |
| Philius | oooooooooM MMMMMMMMMM MMMMMMMMMM MMMMiiiiii iiiiiiiiii |
| PolyPhobius | ooooooooMM MMMMMMMMMM MMMMMMMMMM Miiiiiiiii iiiiiiiiii |
| SCAMPI | oooooooooo oMMMMMMMMM MMMMMMMMMM MMiiiiiiii iiiiiiiiii |
| SPOCTOPUS | ooooooMMMM MMMMMMMMMM MMMMMMMiii iiiiiiiiii iiiiiiiiii |
| PDB-homology |  |
|  | 251                                         291 |
| Seq. | TDLAKSSWLL TIFLGLLAVT VGSNAIITIV ARWSNLSPSA QQVFREMFVF |
| TOPCONS | iiiiiiiiMM MMMMMMMMMM MMMMMMMMMo oooooooooo oooooooooo |
| OCTOPUS | iiiiiiiiMM MMMMMMMMMM MMMMMMMMMo oooooooooo oooooooooo |
| Philius | iiiiiiiMMM MMMMMMMMMM MMMMMMMMMM Mooooooooo oooooooooo |
| PolyPhobius | iiiiiiiMMM MMMMMMMMMM MMMMMMMMMM MMMMoooooo oooooooooo |
| SCAMPI | iiiiiiiiiM MMMMMMMMMM MMMMMMMMMM oooooooooo oooooooooo |
| SPOCTOPUS | iiiiiiiiMM MMMMMMMMMM MMMMMMMMMo oooooooooo oooooooooo |
| PDB-homology |  |
|  | 301                                         341 |
| Seq. | DSDAIQILLT IDHLVATAVF ILPIALFVDV MQTLTRRIER LHYSAIVVGR |
| TOPCONS | ooooooooMM MMMMMMMMMM MMMMMMMMMi iiiiiiiiii iiiiiiiiii |
| OCTOPUS | ooooooooMM MMMMMMMMMM MMMMMMMMMi iiiiiiiiii iiiiiiiiii |
| Philius | ooooooooMM MMMMMMMMMM MMMMMMMMMM iiiiiiiiii iiiiiiiiii |
| PolyPhobius | oooMMMMMMM MMMMMMMMMM MMMMMMMMii iiiiiiiiii iiiiiiiiii |
| SCAMPI | oooooooooo oooMMMMMMM MMMMMMMMMM MMMMiiiiii iiiiiiiiii |
| SPOCTOPUS | ooooooooMM MMMMMMMMMM MMMMMMMMMi iiiiiiiiii iiiiiiiiii |
| PDB-homology |  |
|  | 351                                         391 |
| Seq. | TLPELTMDLK DLAQSFDTVN AAFGPWLGAV FAIELPLIVF CLVSLLDYAS |
| TOPCONS | iiiiiiiiii iiiiiiiiii iiiiMMMMMM MMMMMMMMMM MMMMMooooo |
| OCTOPUS | iiiiiiiiii iiiiiiiiii iiiiMMMMMM MMMMMMMMMM MMMMMooooo |
| Philius | iiiiiiiiii iiiiiiiiii iiMMMMMMMM MMMMMMMMMM MMMMMMoooo |
| PolyPhobius | iiiiiiiiii iiiiiiiiii iiiMMMMMMM MMMMMMMMMM MMMMMMMMoo |
| SCAMPI | iiiiiiiiii iiiiiiiiii iMMMMMMMMM MMMMMMMMMM MMoooooooo |
| SPOCTOPUS | iiiiiiiiii iiiiiiiiii iiiiMMMMMM MMMMMMMMMM MMMMMooooo |
| PDB-homology |  |
|  | 401                                         441 |
| Seq. | LDGLTIFILV IWALMNMVVA AVVALPSARA HSLIHKLEAA IALHLDADST |
| TOPCONS | oooMMMMMMM MMMMMMMMMM MMMMiiiiii iiiiiiiiii iiiiiiiiii |
| OCTOPUS | oooMMMMMMM MMMMMMMMMM MMMMiiiiii iiiiiiiiii iiiiiiiiii |
| Philius | oooMMMMMMM MMMMMMMMMM MMMMMMiiii iiiiiiiiii iiiiiiiiii |
| PolyPhobius | oooMMMMMMM MMMMMMMMMM MMMMMMiiii iiiiiiiiii iiiiiiiiii |
| SCAMPI | oooooooMMM MMMMMMMMMM MMMMMMMMii iiiiiiiiii iiiiiiiiii |
| SPOCTOPUS | oooMMMMMMM MMMMMMMMMM MMMMiiiiii iiiiiiiiii iiiiiiiiii |
| PDB-homology |  |
|  | 451                                         491 |
| Seq. | LVERSDYVFL VTTLNAHNMG IKIAGATNIT FTLMSQAVSF IGSFYLLLVG |
| TOPCONS | iiiiiiiiii iiiiiiiiii iiiiiiiiii MMMMMMMMMM MMMMMMMMMM |
| OCTOPUS | iiiiiiiiii iiiiiiiiii iiiiiiiiii MMMMMMMMMM MMMMMMMMMM |
| Philius | iiiiiiiiii iiiiiiiiii iiiiiiiiMM MMMMMMMMMM MMMMMMMMMo |
| PolyPhobius | iiiiiiiiii iiiiiiiiii iiiiiiiMMM MMMMMMMMMM MMMMMMMMMM |
| SCAMPI | iiiiiiiiii iiiiiiiiii iiiiiiiiii iiMMMMMMMM MMMMMMMMMM |
| SPOCTOPUS | iiiiiiiiii iiiiiiiiii iiiiiiiiiM MMMMMMMMMM MMMMMMMMMM |
| PDB-homology |  |

|  |  |
| --- | --- |
|  | 501 |
| Seq. | FRTS |
| TOPCONS | Mooo |
| OCTOPUS | Mooo |
| Philius | oooo |
| PolyPhobius | Mooo |
| SCAMPI | MMMo |
| SPOCTOPUS | oooo |
| PDB-homology |  |
