## Supplementary material for "A putative origin of insect chemosensory receptors in the last common eukaryotic ancestor": Data S2: nicetop.html

|  |  |
| --- | --- |
|  | 1                                           41 |
| Seq. | MRLRHKHSSS RLHEIRTDAE SGPLGAESLS SSSELSHDIL LVNEKADFDE |
| TOPCONS | iiiiiiiiii iiiiiiiiii iiiiiiiiii iiiiiiiiii iiiiiiiiii |
| OCTOPUS | iiiiiiiiii iiiiiiiiii iiiiiiiiii iiiiiiiiii iiiiiiiiii |
| Philius | iiiiiiiiii iiiiiiiiii iiiiiiiiii iiiiiiiiii iiiiiiiiii |
| PolyPhobius | iiiiiiiiii iiiiiiiiii iiiiiiiiii iiiiiiiiii iiiiiiiiii |
| SCAMPI | iiiiiiiiii iiiiiiiiii iiiiiiiiii iiiiiiiiii iiiiiiiiii |
| SPOCTOPUS | iiiiiiiiii iiiiiiiiii iiiiiiiiii iiiiiiiiii iiiiiiiiii |
| PDB-homology |  |
|  | 51                                          91 |
| Seq. | IMRNMPWIAA YQAVLMTLGM DQAAGKRRSV ITTFRSLMLS LLLFSVSLVA |
| TOPCONS | iiiiiiiiii iiiiiiiiii iiiiiiiiii iiMMMMMMMM MMMMMMMMMM |
| OCTOPUS | iiiiiiiiii iiiiiiiiii iiiiiiiiii iMMMMMMMMM MMMMMMMMMM |
| Philius | iiiiiiiiii iiiiiiiiii iiiiiiiiii iiMMMMMMMM MMMMMMMMMM |
| PolyPhobius | iiiiiiiiii iiiiiiiiii iiiiiiiiii iiiiiMMMMM MMMMMMMMMM |
| SCAMPI | iiiiiiiiii iiiiiiiiii iiiiiiiiii iiiiiMMMMM MMMMMMMMMM |
| SPOCTOPUS | iiiiiiiiii iiiiiiiiii iiiiiiiiii iMMMMMMMMM MMMMMMMMMM |
| PDB-homology |  |
|  | 101                                         141 |
| Seq. | FYNATRNERW TMLTLIGLWY FHCGVFLGVM TLHMPSRWFR HLIVSLVVHA |
| TOPCONS | MMMooooooo MMMMMMMMMM MMMMMMMMMM Miiiiiiiii iiiiiiiiii |
| OCTOPUS | MMoooooooo MMMMMMMMMM MMMMMMMMMM Miiiiiiiii iiiiiiiiii |
| Philius | MMMMoooooo oMMMMMMMMM MMMMMMMMMM MMMiiiiiii iiiiiiiiii |
| PolyPhobius | MMMMoooooM MMMMMMMMMM MMMMMMMMMM MMiiiiiiii iiiiiiiiii |
| SCAMPI | MMMMMMoooo oMMMMMMMMM MMMMMMMMMM MMiiiiiiii iiiiiiiiii |
| SPOCTOPUS | MMoooooooo MMMMMMMMMM MMMMMMMMMM Miiiiiiiii iiiiiiiiii |
| PDB-homology |  |
|  | 151                                         191 |
| Seq. | ADDPLVSLSS ESDGGRGSRS RSEAVRQRAH ADILPTHFLK ILNHGFRYLM |
| TOPCONS | iiiiiiiiii iiiiiiiiii iiiiiiiiii iiiiiiiiii iiiiiiiMMM |
| OCTOPUS | iiiiiiiiii iiiiiiiiii iiiiiiiiii iiiiiiiiii iiiiiiMMMM |
| Philius | iiiiiiiiii iiiiiiiiii iiiiiiiiii iiiiiiiiii iiiiiiiiMM |
| PolyPhobius | iiiiiiiiii iiiiiiiiii iiiiiiiiii iiiiiiiiii iiiiiiiMMM |
| SCAMPI | iiiiiiiiii iiiiiiiiii iiiiiiiiii iiiiiiiiii iiiiiiiiMM |
| SPOCTOPUS | iiiiiiiiii iiiiiiiiii iiiiiiiiii iiiiiiiiii iiiiiiMMMM |
| PDB-homology |  |
|  | 201                                         241 |
| Seq. | IAQLVAITFN VSIGAAFFID IGTATDVYIS IIQPFDSLGF KIFAFFSNTV |
| TOPCONS | MMMMMMMMMM MMMMMMMMoo oooooooooo oooooooooo oooooooMMM |
| OCTOPUS | MMMMMMMMMM MMMMMMMooo oooooooooo oooooooooo oooooooMMM |
| Philius | MMMMMMMMMM MMMMMMMMMo oooooooooo oooooooooo oMMMMMMMMM |
| PolyPhobius | MMMMMMMMMM MMMMMMMMMo oooooooooo oooooooooo oMMMMMMMMM |
| SCAMPI | MMMMMMMMMM MMMMMMMMMo oooooooooo oooooooooo oooooooooM |
| SPOCTOPUS | MMMMMMMMMM MMMMMMMooo oooooooooo oooooooooo oooooooMMM |
| PDB-homology |  |
|  | 251                                         291 |
| Seq. | ATVVWIFPLP PFLLSLYILH KQFAHLADAI RRGAIPLPED MAVTHQRIHR |
| TOPCONS | MMMMMMMMMM MMMMMMMMii iiiiiiiiii iiiiiiiiii iiiiiiiiii |
| OCTOPUS | MMMMMMMMMM MMMMMMMMii iiiiiiiiii iiiiiiiiii iiiiiiiiii |
| Philius | MMMMMMMMMM MMMMMMiiii iiiiiiiiii iiiiiiiiii iiiiiiiiii |
| PolyPhobius | MMMMMMMMMM MMMMMMMiii iiiiiiiiii iiiiiiiiii iiiiiiiiii |
| SCAMPI | MMMMMMMMMM MMMMMMMMMM iiiiiiiiii iiiiiiiiii iiiiiiiiii |
| SPOCTOPUS | MMMMMMMMMM MMMMMMMMii iiiiiiiiii iiiiiiiiii iiiiiiiiii |
| PDB-homology |  |
|  | 301                                         341 |
| Seq. | RSVFLTHFAR WPIGLNISSH AVFSVFLSYR IVTQDSADME LLVIEMFWLL |
| TOPCONS | iiiiiiiiMM MMMMMMMMMM MMMMMMMMMo oooooooooo oooooMMMMM |
| OCTOPUS | iiiiiiiiii iiiiiiiiii iiiiiiiiii iiiiiiiiii iiMMMMMMMM |
| Philius | iiiiiiMMMM MMMMMMMMMM MMMMMMMMMo oooooooooo oooooMMMMM |
| PolyPhobius | iiiiiiiMMM MMMMMMMMMM MMMMMMMMMo oooooooooo oooooMMMMM |
| SCAMPI | iiiiiiiiii MMMMMMMMMM MMMMMMMMMM Mooooooooo oooooMMMMM |
| SPOCTOPUS | iiiiiiiiii iiiiiiiiii iiiiiiiiii iiiiiiiiii iiMMMMMMMM |
| PDB-homology |  |
|  | 351                                         391 |
| Seq. | AATLVIFVIT FLAAYVHSSG SEAMVFAIAD VHAPTFDMRT ELEYALSTVE |
| TOPCONS | MMMMMMMMMM MMMMMMiiii iiiiiiiiii iiiiiiiiii iiiiiiiiii |
| OCTOPUS | MMMMMMMMMM MMMooooooo oooooooooo oooooooooo oooooooooo |
| Philius | MMMMMMMMMM MMMMMMiiii iiiiiiiiii iiiiiiiiii iiiiiiiiii |
| PolyPhobius | MMMMMMMMMM MMMMMMiiii iiiiiiiiii iiiiiiiiii iiiiiiiiii |
| SCAMPI | MMMMMMMMMM MMMMMMiiii iiiiiiiiii iiiiiiiiii iiiiiiiiii |
| SPOCTOPUS | MMMMMMMMMM MMMooooooo oooooooooo oooooooooo oooooooooo |
| PDB-homology |  |

|  |  |
| --- | --- |
|  | 401                              431 |
| Seq. | RRASGFRLAK TFLITWTVLA KIVSLFVSFL ALLNNMRGRN |
| TOPCONS | iiiiiiiiii iiMMMMMMMM MMMMMMMMMM MMMooooooo |
| OCTOPUS | oooooooooo MMMMMMMMMM MMMMMMMMMM Miiiiiiiii |
| Philius | iiiiiiiiii iMMMMMMMMM MMMMMMMMMM MMMooooooo |
| PolyPhobius | iiiiiiiiii MMMMMMMMMM MMMMMMMMMM MMMooooooo |
| SCAMPI | iiiiiiiiii iiMMMMMMMM MMMMMMMMMM MMMooooooo |
| SPOCTOPUS | oooooooooo MMMMMMMMMM MMMMMMMMMM Miiiiiiiii |
| PDB-homology |  |
