## Supplementary material for "A putative origin of insect chemosensory receptors in the last common eukaryotic ancestor": Data S2: nicetop.html

|  |  |
| --- | --- |
|  | 1                                           41 |
| Seq. | MFVPLHSGSR SRGAVGPVNG GEGAGMPGPS ALHCVPEHVQ PMRLLLSLLA |
| TOPCONS | iiiiiiiiii iiiiiiiiii iiiiiiiiii iiiiiiiiii iiiiiiiiii |
| OCTOPUS | iiiiiiiiii iiiiiiiiii iiiiiiiiii iiiiiiiiii iiiiiiiiii |
| Philius | iiiiiiiiii iiiiiiiiii iiiiiiiiii iiiiiiiiii iiiiiiiiii |
| PolyPhobius | iiiiiiiiii iiiiiiiiii iiiiiiiiii iiiiiiiiii iiiiiiiiii |
| SCAMPI | iiiiiiiiii iiiiiiiiii iiiiiiiiii iiiiiiiiii iiiiiiiiii |
| SPOCTOPUS | iiiiiiiiii iiiiiiiiii iiiiiiiiii iiiiiiiiii iiiiiiiiii |
| PDB-homology |  |
|  | 51                                          91 |
| Seq. | VAPSRSGLYR GLWRPLLFTL VLAGTGALSW AALLDTRFPK ALIVGTAGLS |
| TOPCONS | iiiiiiiiii iMMMMMMMMM MMMMMMMMMM MMoooooooo MMMMMMMMMM |
| OCTOPUS | iiiiiiiiii iMMMMMMMMM MMMMMMMMMM MMoooooooo MMMMMMMMMM |
| Philius | iiiiiiiiii iMMMMMMMMM MMMMMMMMMM MMMMoooooo MMMMMMMMMM |
| PolyPhobius | iiiiiiiiii iiMMMMMMMM MMMMMMMMMM MMMMoooooM MMMMMMMMMM |
| SCAMPI | iiiiiiiiii iiiiMMMMMM MMMMMMMMMM MMMMMooooo MMMMMMMMMM |
| SPOCTOPUS | iiiiiiiiii iMMMMMMMMM MMMMMMMMMM MMoooooooo MMMMMMMMMM |
| PDB-homology |  |
|  | 101                                         141 |
| Seq. | LASVAGWLSL AHYLGSGHSR ASAGFLLSLA EGRMRLRLRS TTRVYAFGAL |
| TOPCONS | MMMMMMMMMM Miiiiiiiii iiiiiiiiii iiiiiiiiii iiiMMMMMMM |
| OCTOPUS | MMMMMMMMMM Miiiiiiiii iiiiiiiiii iiiiiiiiii iMMMMMMMMM |
| Philius | MMMMMMMMMM MMMMiiiiii iiiiiiiiii iiiiiiiiii iiiMMMMMMM |
| PolyPhobius | MMMMMMMMMM MMiiiiiiii iiiiiiiiii iiiiiiiiii iiiMMMMMMM |
| SCAMPI | MMMMMMMMMM Miiiiiiiii iiiiiiiiii iiiiiiiiii iiiMMMMMMM |
| SPOCTOPUS | MMMMMMMMMM Miiiiiiiii iiiiiiiiii iiiiiiiiii iMMMMMMMMM |
| PDB-homology |  |
|  | 151                                         191 |
| Seq. | LASVVLSVGW LAVSWMAVSG IDNNDGVDDD PDSSMPIFYI LYAVAVPVNV |
| TOPCONS | MMMMMMMMMM MMMMoooooo oooooooooo oooooooooo MMMMMMMMMM |
| OCTOPUS | MMMMMMMMMM MMoooooooo oooooooooo oooooooooo MMMMMMMMMM |
| Philius | MMMMMMMMMM MMMMoooooo oooooooooo ooooooooMM MMMMMMMMMM |
| PolyPhobius | MMMMMMMMMM MMMMMMMooo oooooooooo ooooooMMMM MMMMMMMMMM |
| SCAMPI | MMMMMMMMMM MMMMoooooo oooooooooo ooooooMMMM MMMMMMMMMM |
| SPOCTOPUS | MMMMMMMMMM MMoooooooo oooooooooo oooooooooo MMMMMMMMMM |
| PDB-homology |  |
|  | 201                                         241 |
| Seq. | TVFVFAVAVA ALMAHISRAQ VAALAVSMED GTLSMERVVR KHATMARLAR |
| TOPCONS | MMMMMMMMMM Miiiiiiiii iiiiiiiiii iiiiiiiiii iiiiiiiiii |
| OCTOPUS | MMMMMMMMMM Miiiiiiiii iiiiiiiiii iiiiiiiiii iiiiiiiiii |
| Philius | MMMMMMMMMM MMiiiiiiii iiiiiiiiii iiiiiiiiii iiiiiiiiii |
| PolyPhobius | MMMMMMMMMM MMiiiiiiii iiiiiiiiii iiiiiiiiii iiiiiiiiii |
| SCAMPI | MMMMMMMiii iiiiiiiiii iiiiiiiiii iiiiiiiiii iiiiiiiiii |
| SPOCTOPUS | MMMMMMMMMM Miiiiiiiii iiiiiiiiii iiiiiiiiii iiiiiiiiii |
| PDB-homology |  |
|  | 251                                         291 |
| Seq. | RLSSLMQSFV VPVFIIFPLS ISYQLWDAAR ASERADNELN GKYYLVIYAL |
| TOPCONS | iiiiMMMMMM MMMMMMMMMM MMMMMooooo oooooooooo ooMMMMMMMM |
| OCTOPUS | iiiiiMMMMM MMMMMMMMMM MMMMMMoooo oooooooooo ooMMMMMMMM |
| Philius | iiiMMMMMMM MMMMMMMMMM MMMooooooo oooooooooo oooMMMMMMM |
| PolyPhobius | iiiMMMMMMM MMMMMMMMMM MMMooooooo oooooooooo ooMMMMMMMM |
| SCAMPI | iiiiMMMMMM MMMMMMMMMM MMMMMooooo oooooooooo oMMMMMMMMM |
| SPOCTOPUS | iiiiiMMMMM MMMMMMMMMM MMMMMMoooo oooooooooo ooMMMMMMMM |
| PDB-homology |  |
|  | 301                                         341 |
| Seq. | ALILVATIFT SINRKFSSLT DTAAQLHRVG SHSFTDINTL LGYLVASRPP |
| TOPCONS | MMMMMMMMMM MMMiiiiiii iiiiiiiiii iiiiiiiiii iiiiiiiiii |
| OCTOPUS | MMMMMMMMMM MMMiiiiiii iiiiiiiiii iiiiiiiiii iiiiiiiiii |
| Philius | MMMMMMMMMM MMMiiiiiii iiiiiiiiii iiiMMMMMMM MMMMMMMMMM |
| PolyPhobius | MMMMMMMMMM MMiiiiiiii iiiiiiiiii iiiiiiiiii iiiiiiiiii |
| SCAMPI | MMMMMMMMMM MMiiiiiiii iiiiiiiiii iiiiiiiiii iiiiiiiiii |
| SPOCTOPUS | MMMMMMMMMM MMMiiiiiii iiiiiiiiii iiiiiiiiii iiiiiiiiii |
| PDB-homology |  |

|  |  |
| --- | --- |
|  | 351                              381 |
| Seq. | FLIMGAAPSP AIVPILILAF FGVVGTLIHQ VSL |
| TOPCONS | iiiiiiiiMM MMMMMMMMMM MMMMMMMMMo ooo |
| OCTOPUS | iiiiiiiiii MMMMMMMMMM MMMMMMMMMM Moo |
| Philius | MMoooMMMMM MMMMMMMMMM MMMMMMMMMM Mii |
| PolyPhobius | iMMMMMMMMM MMMMMMMMMM MMMMMMMMMo ooo |
| SCAMPI | iiiiiiiMMM MMMMMMMMMM MMMMMMMMoo ooo |
| SPOCTOPUS | iiiiiiiiii MMMMMMMMMM MMMMMMMMMM Moo |
| PDB-homology |  |
