## Supplementary material for "A putative origin of insect chemosensory receptors in the last common eukaryotic ancestor": Data S2: nicetop.html

|  |  |
| --- | --- |
|  | 1                                           41 |
| Seq. | MEPLRFLITA VGLSPSPTVF NRAYQGLVLA ALLAAPILLF VIGLNKYESR |
| TOPCONS | oooooooooo oooooooooo ooMMMMMMMM MMMMMMMMMM MMMiiiiiii |
| OCTOPUS | oooooooooo oooooooooo ooMMMMMMMM MMMMMMMMMM MMMiiiiiii |
| Philius | oooooooooo oooooooooo ooMMMMMMMM MMMMMMMMMM MMMMiiiiii |
| PolyPhobius | oooooooooo oooooooooo oooMMMMMMM MMMMMMMMMM MMMMiiiiii |
| SCAMPI | iiiiiiiiii iiiiiiiiii iiiiiiMMMM MMMMMMMMMM MMMMMMMooo |
| SPOCTOPUS | oooooooooo oooooooooo ooMMMMMMMM MMMMMMMMMM MMMiiiiiii |
| PDB-homology |  |
|  | 51                                          91 |
| Seq. | LTESTLVGAA ILALIPVISH LMVMPYLARN YIADEVHVCL AADPSARKKI |
| TOPCONS | iiiMMMMMMM MMMMMMMMMM MMMMoooooo oooooooooo oooooooooo |
| OCTOPUS | iiiMMMMMMM MMMMMMMMMM MMMMoooooo oooooooooo oooooooooo |
| Philius | iiiMMMMMMM MMMMMMMMMM MMMMoooooo oooooooooo oooooooooo |
| PolyPhobius | iiiMMMMMMM MMMMMMMMMM MMMMMMMooM MMMMMMMMMM MMMMiiiiii |
| SCAMPI | oooMMMMMMM MMMMMMMMMM MMMMiiiiii iiiiiiiiii iiiiiiiiii |
| SPOCTOPUS | iiiMMMMMMM MMMMMMMMMM MMMMoooooo oooooooooo oooooooooo |
| PDB-homology |  |
|  | 101                                         141 |
| Seq. | MALGTAHAWG GLVIGTTFAV LWQVNLFRAL KDGRDWARDS SGLKTLFVLQ |
| TOPCONS | ooooMMMMMM MMMMMMMMMM MMMMMiiiii iMMMMMMMMM MMMMMMMMMM |
| OCTOPUS | oooMMMMMMM MMMMMMMMMM MMMMiiiiii iiiiiiiiii iiiMMMMMMM |
| Philius | ooooMMMMMM MMMMMMMMMM MMMMMMiiii iiiiiiiiii iiiiiMMMMM |
| PolyPhobius | iiiiMMMMMM MMMMMMMMMM MMMMMMMMMo oooooooooo ooooMMMMMM |
| SCAMPI | iiiiiMMMMM MMMMMMMMMM MMMMMMoooo oooooooooo ooooMMMMMM |
| SPOCTOPUS | ooooMMMMMM MMMMMMMMMM MMMMMiiiii iiiiiiiiii iiiMMMMMMM |
| PDB-homology |  |
|  | 151                                         191 |
| Seq. | AVAVSVSVAS FTMALVLYSL VCNIIRMQAG TFSTGIKSGK YANEAIVYGH |
| TOPCONS | MMoMMMMMMM MMMMMMMMMM MMMMiiiiii iiiiiiiiii iiiiiiiiii |
| OCTOPUS | MMMMMMMMoM MMMMMMMMMM MMMMiiiiii iiiiiiiiii iiiiiiiiii |
| Philius | MMMMMMMMMM MMMMMMMMMM Mooooooooo oooooooooo oooooooooo |
| PolyPhobius | MMMMMMMMMM MMMMMMMMMM Miiiiiiiii iiiiiiiiii iiiiiiiiii |
| SCAMPI | MMMMMMMMMM MMMMMiiiii iiiiiiiiii iiiiiiiiii iiiiiiiiii |
| SPOCTOPUS | MMMMMMMMoM MMMMMMMMMM MMMMiiiiii iiiiiiiiii iiiiiiiiii |
| PDB-homology |  |
|  | 201                                         241 |
| Seq. | HHIKTSVAVT NQMFSRWIFP AIVVTLFSIS FQTVGYFHQD KHSSQYNGAY |
| TOPCONS | iiiiiiiiii iiMMMMMMMM MMMMMMMMMM MMMooooooo ooooooooMM |
| OCTOPUS | iiiiiiiiii iiMMMMMMMM MMMMMMMMMM MMMooooooo ooooooooMM |
| Philius | oooooooooo ooooooMMMM MMMMMMMMMM MMMMMMMiii iiiiiiiMMM |
| PolyPhobius | iiiiiiiiii iiMMMMMMMM MMMMMMMMMM MMMMMMMooo ooooooooMM |
| SCAMPI | iiiiiiiiii iiMMMMMMMM MMMMMMMMMM MMMooooooo ooooooooMM |
| SPOCTOPUS | iiiiiiiiii iiMMMMMMMM MMMMMMMMMM MMMooooooo ooooooooMM |
| PDB-homology |  |
|  | 251                                         291 |
| Seq. | YIIVPAFLLV FLIVVASTVT GKIDRLKYNV GALLSSDYTR AGTLSHVLAY |
| TOPCONS | MMMMMMMMMM MMMMMMMMMi iiiiiiiiii iiiiiiiiii iiiiiiiiii |
| OCTOPUS | MMMMMMMMMM MMMMMMMMMi iiiiiiiiii iiiiiiiiii iiiiiiiiii |
| Philius | MMMMMMMMMM MMMMMMMMMM oooooooooo oooooooooo oooooooooo |
| PolyPhobius | MMMMMMMMMM MMMMMMMMMi iiiiiiiiii iiMMMMMMMM MMMMMMMooo |
| SCAMPI | MMMMMMMMMM MMMMMMMMMi iiiiiiiiii iiiiiiiiii iiiiiiiiii |
| SPOCTOPUS | MMMMMMMMMM MMMMMMMMMi iiiiiiiiii iiiiiiiiii iiiiiiiiii |
| PDB-homology |  |

|  |  |
| --- | --- |
|  | 301                              331 |
| Seq. | LQNVDDSFKI FGVAITSSLS HLIVFAAIAS VGAFVWEGIH |
| TOPCONS | iiiiiiiiii iiiiiiiMMM MMMMMMMMMM MMMMMMMMoo |
| OCTOPUS | iiiiiiiiii iiiiiiiMMM MMMMMMMMMM MMMMMMMMoo |
| Philius | oooooooooo MMMMMMMMMM MMMMMMMMMM MMMMMMiiii |
| PolyPhobius | oooooooooM MMMMMMMMMM MMMMMMMMMM MMMMMiiiii |
| SCAMPI | iiiiiiiiii iiiiiiiMMM MMMMMMMMMM MMMMMMMMoo |
| SPOCTOPUS | iiiiiiiiii iiiiiiiMMM MMMMMMMMMM MMMMMMMMoo |
| PDB-homology |  |
