## Supplementary material for "A putative origin of insect chemosensory receptors in the last common eukaryotic ancestor": Data S2: nicetop.html

|  |  |
| --- | --- |
|  | 1                                           41 |
| Seq. | MSSASAYSDY SPSSADAPIV LDDHKDDEVS SSRDDPNAGS VSDVSLASGD |
| TOPCONS | iiiiiiiiii iiiiiiiiii iiiiiiiiii iiiiiiiiii iiiiiiiiii |
| OCTOPUS | oooooooooo oooooooooo oooooooooo oooooooooo oooooooooo |
| Philius | iiiiiiiiii iiiiiiiiii iiiiiiiiii iiiiiiiiii iiiiiiiiii |
| PolyPhobius | iiiiiiiiii iiiiiiiiii iiiiiiiiii iiiiiiiiii iiiiiiiiii |
| SCAMPI | oooooooooo oooooooooo oooooooooo oooooooooo oooooooooo |
| SPOCTOPUS | iiiiiiiiii iiiiiiiiii iiiiiiiiii iiiiiiiiii iiiiiiiiii |
| PDB-homology |  |
|  | 51                                          91 |
| Seq. | LISDNGPPHV ALLPARMNSD GELVAVAPMA RGSSARSLTL LQPMAGWLAF |
| TOPCONS | iiiiiiiiii iiiiiiiiii iiiiiiiiii iiiiiiiiii iiiiiiiiii |
| OCTOPUS | oooooooooo oooooooooo oooooooooo oooooooooo oooooooooo |
| Philius | iiiiiiiiii iiiiiiiiii iiiiiiiiii iiiiiiiiii iiiMMMMMMM |
| PolyPhobius | iiiiiiiiii iiiiiiiiii iiiiiiiiii iiiiiiiiiM MMMMMMMMMM |
| SCAMPI | oooooooooo oooooooooo oooooooooo oooooooooo oooMMMMMMM |
| SPOCTOPUS | iiiiiiiiii iiiiiiiiii iiiiiiiiii iiiiiiiiii iiiiiiiiii |
| PDB-homology |  |
|  | 101                                         141 |
| Seq. | CGYAPGGGIV GTGWAFLVWA MVVTHAVVMF SLTVYPMRLT ESTLSGLAIL |
| TOPCONS | iiiiiiiiii MMMMMMMMMM MMMMMMMMMM Mooooooooo oMMMMMMMMM |
| OCTOPUS | oooooooooo MMMMMMMMMM MMMMMMMMMM Miiiiiiiii iMMMMMMMMM |
| Philius | MMMMMMMMMM MMMMMMMMMM MMMooooooo ooooooooMM MMMMMMMMMM |
| PolyPhobius | MMMMoMMMMM MMMMMMMMMM iMMMMMMMMM MMMMMMoooo ooMMMMMMMM |
| SCAMPI | MMMMMMMMMM MMMMiMMMMM MMMMMMMMMM MMMMMMoooo oMMMMMMMMM |
| SPOCTOPUS | iiiiiiiiii MMMMMMMMMM MMMMMMMMMM Mooooooooo oMMMMMMMMM |
| PDB-homology |  |
|  | 151                                         191 |
| Seq. | ALAPVITLPT FNRYMRSGVL EDEVATCKYV HQYSGRAVRQ LALGHTLAAM |
| TOPCONS | MMMMMMMMMM MMiiiiiiii iiiiiiiiii iiiiiiiiii iiMMMMMMMM |
| OCTOPUS | MMMMMMMMMM MMoooooooo oooooooooo oooooooooo ooMMMMMMMM |
| Philius | MMMMMMMMii iiiiiiiiii iiiiiiiiii iiiiiiiiii iiMMMMMMMM |
| PolyPhobius | MMMMMMMMMM Miiiiiiiii iiiiiiiiii iiiiiiiiii iMMMMMMMMM |
| SCAMPI | MMMMMMMMMM MMiiiiiiii iiiiiiiiii iiiiiiiiii iiiiiMMMMM |
| SPOCTOPUS | MMMMMMMMMM MMiiiiiiii iiiiiiiiii iiiiiiiiii iiMMMMMMMM |
| PDB-homology |  |
|  | 201                                         241 |
| Seq. | AVSLFLVVGW NVTLGLALDG QGFGGGKSDA LYWVELVTSI FAIPAMVTPL |
| TOPCONS | MMMMMMMMMM MMMooooooo oooooooooo oooooMMMMM MMMMMMMMMM |
| OCTOPUS | MMMMMMMMMM MMMiiiiiii iiiiiiiiii MMMMMMMMMM MMMMMoMMMM |
| Philius | MMMMMMMMMM MMMMoooooo oooooooooo oooooMMMMM MMMMMMMMMM |
| PolyPhobius | MMMMMMMMMM MMMooooooo oooooooooM MMMMMMMMMM MMMMMMMMMM |
| SCAMPI | MMMMMMMMMM MMMMMMoooo oooooooooo oooooMMMMM MMMMMMMMMM |
| SPOCTOPUS | MMMMMMMMMM MMMooooooo oooooooooo ooooMMMMMM MMMMMMMMMM |
| PDB-homology |  |
|  | 251                                         291 |
| Seq. | AMLSLVCNLH RLQCLSFIGR LTDADMHIPS ALRLHQAIRR SIKASSRLFT |
| TOPCONS | MMMMMMiiii iiiiiiiiii iiiiiiiiii iiiiiiiiii iiiiiiiMMM |
| OCTOPUS | MMMMMMMMMM Miiiiiiiii iiiiiiiiii iiiiiiiiii iiiiiiiMMM |
| Philius | MMMMMMMMMi iiiiiiiiii iiiiiiiiii iiiiiiiiii iiiiiiiMMM |
| PolyPhobius | MMMMMMiiii iiiiiiiiii iiiiiiiiii iiiiiiiiii iiiiiiiiMM |
| SCAMPI | MMMMMMiiii iiiiiiiiii iiiiiiiiii iiiiiiiiii iiiiiiiMMM |
| SPOCTOPUS | MMMMMiiiii iiiiiiiiii iiiiiiiiii iiiiiiiiii iiiiiiiMMM |
| PDB-homology |  |
|  | 301                                         341 |
| Seq. | RWVFPAALIC AVSLTYQIYG YLLSHSSSRQ YNGIYHIIAV TLILVVILGA |
| TOPCONS | MMMMMMMMMM MMMMMMMMoo oooooooooo oooMMMMMMM MMMMMMMMMM |
| OCTOPUS | MMMMMMMMMM MMMMMMMMoo oooooooooo oooMMMMMMM MMMMMMMMMM |
| Philius | MMMMMMMMMM MMMMMMMMMo oooooooooo oooMMMMMMM MMMMMMMMMM |
| PolyPhobius | MMMMMMMMMM MMMMMMMMMM Mooooooooo oooMMMMMMM MMMMMMMMMM |
| SCAMPI | MMMMMMMMMM MMMMMMMMoo oooooooooo oooMMMMMMM MMMMMMMMMM |
| SPOCTOPUS | MMMMMMMMMM MMMMMMMMoo oooooooooo oooMMMMMMM MMMMMMMMMM |
| PDB-homology |  |
|  | 351                                         391 |
| Seq. | TTSVTRAAHN VRHATAELYA VSFGSAADIS RFMVYLNNTD LAFRIFAVPM |
| TOPCONS | MMMMiiiiii iiiiiiiiii iiiiiiiiii iiiiiiiiii iiiiiiiiii |
| OCTOPUS | MMMMiiiiii iiiiiiiiii iiiiiiiiii iiiiiiiiii iiiiiiiiii |
| Philius | MMMMMiiiii iiiiMMMMMM MMMMMMMMMM MMMMMooooo oooooooooo |
| PolyPhobius | MMMMiiiiii iiiiiMMMMM MMMMMMMMMM MMMMMMoooo oooooooooo |
| SCAMPI | MMMMiiiiii iiiiiiiiii iiiiiiiiii iiiiiiiiii iiiiiiiiii |
| SPOCTOPUS | MMMMiiiiii iiiiiiiiii iiiiiiiiii iiiiiiiiii iiiiiiiiii |
| PDB-homology |  |

|  |  |
| --- | --- |
|  | 401                   421 |
| Seq. | YPAYNSLVIV GFSAVVGGVV RQATI |
| TOPCONS | MMMMMMMMMM MMMMMMMMMM Moooo |
| OCTOPUS | iiMMMMMMMM MMMMMMMMMM MMMoo |
| Philius | MMMMMMMMMM MMMMMMMMMM iiiii |
| PolyPhobius | MMMMMMMMMM MMMMMMMMMM iiiii |
| SCAMPI | MMMMMMMMMM MMMMMMMMMM Moooo |
| SPOCTOPUS | iiMMMMMMMM MMMMMMMMMM MMMoo |
| PDB-homology |  |
