## Supplementary material for "A putative origin of insect chemosensory receptors in the last common eukaryotic ancestor": Data S2: nicetop.html

|  |  |
| --- | --- |
|  | 1                                           41 |
| Seq. | MSALSYEELP VLEYREEDED DHFLVAGENR ISFNQFRWLP IWVNTLGVSE |
| TOPCONS | oooooooooo oooooooooo oooooooooo oooooooooo oooooooooo |
| OCTOPUS |  |
| Philius | iiiiiiiiii iiiiiiiiii iiiiiiiiii iiiiiiiiii iiiiiiiiii |
| PolyPhobius | iiiiiiiiii iiiiiiiiii iiiiiiiiii iiiiiiiiii iiiiiiiiii |
| SCAMPI |  |
| SPOCTOPUS | oooooooooo oooooooooo oooooooooo oooooooooo oooooooooo |
| PDB-homology |  |
|  | 51                                          91 |
| Seq. | DGHLWRPIVT IISMLVWTIW QIVIQFYVGL TIHSQIGGTE ITIQSIGNPV |
| TOPCONS | oooooooooo oooooooooo oooooooooo oooooooooo ooooooooMM |
| OCTOPUS |  |
| Philius | iiiiiiiMMM MMMMMMMMMM MMMMMMMMMo oooooooooo ooooooooMM |
| PolyPhobius | iiiiiiiMMM MMMMMMMMMM MMMMMMMMMM oooooooooo ooooooooMM |
| SCAMPI |  |
| SPOCTOPUS | ooooooMMMM MMMMMMMMMM MMMMMMMooo oooooooooo ooooooooMM |
| PDB-homology |  |
|  | 101                                         141 |
| Seq. | WVILSLYGLL SYAYLLWYMR YHKRHYKTLI TKQLSVEPKI ETEISNKILR |
| TOPCONS | MMMMMMMMMM MMMMMMMMMi iiiiiiiiii iiiiiiiiii iiiiiiiiii |
| OCTOPUS |  |
| Philius | MMMMMMMMMM MMMMMMMMii iiiiiiiiii iiiiiiiiii iiiiiiiiii |
| PolyPhobius | MMMMMMMMMM MMMMMMMMii iiiiiiiiii iiiiiiiiii iiiiiiiiii |
| SCAMPI |  |
| SPOCTOPUS | MMMMMMMMMM MMMMMMMMMo oooooooooo oooooooooo oooooooooo |
| PDB-homology |  |
|  | 151                                         191 |
| Seq. | NMILTLTVAL LFLIASYIKS IIAMVHYSKY HEIPYKTMQS LPASIFILHS |
| TOPCONS | MMMMMMMMMM MMMMMMMMMM Mooooooooo oooooooooo oooooooooo |
| OCTOPUS |  |
| Philius | iMMMMMMMMM MMMMMMMMMM MMMooooooo oooooooooo oooooooooo |
| PolyPhobius | iiMMMMMMMM MMMMMMMMMM MMMMMMoooo oooooooooo oooooooooo |
| SCAMPI |  |
| SPOCTOPUS | MMMMMMMMMM MMMMMMMMMM Mooooooooo oooooooooo oooooooooo |
| PDB-homology |  |
|  | 201                                         241 |
| Seq. | LVEPIMWFAG SISIFSCFLI MDLLGWVHAV DIRIYCSQLI YAARSTGQPD |
| TOPCONS | oooooooooo oooooooooo oooooooooo oooooooooo oooooooooo |
| OCTOPUS |  |
| Philius | ooooMMMMMM MMMMMMMMMM MMMMMMiiii iiiiiiiiii iiiiiiiiii |
| PolyPhobius | ooooMMMMMM MMMMMMMMMM MMMMMMiiii iiiiiiiiii iiiiiiiiii |
| SCAMPI |  |
| SPOCTOPUS | MMMMMMMMMM MMMMMMMMMM MMMMMMMMMM Mooooooooo oooooooooo |
| PDB-homology |  |
|  | 251                                         291 |
| Seq. | NRIPIDVLIT QHQRLRTDIK KTQKLWTPVI LGSLFLGTAL LLSSVIMWRM |
| TOPCONS | oooooooooo oooooooooo oooooooooo oooooooooo oooooooooo |
| OCTOPUS |  |
| Philius | iiiiiiiiii iiiiiiiiii iiiiMMMMMM MMMMMMMMMM MMMMMMMMoo |
| PolyPhobius | iiiiiiiiii iiiiiiiiii iiiiMMMMMM MMMMMMMMMM MMMMMMMMoo |
| SCAMPI |  |
| SPOCTOPUS | oooooooooo oooooooooo oooooMMMMM MMMMMMMMMM MMMMMMoooo |
| PDB-homology |  |
|  | 301                                         341 |
| Seq. | NQVQYDDITP PLIVVLFLVV SSVYLLNIAY VNKFGGELCG QLYQSSALKS |
| TOPCONS | oooooooooo oooooooooo oooooooooo oooooooooo oooooooooo |
| OCTOPUS |  |
| Philius | oooooooMMM MMMMMMMMMM MMMMMMMMMM Miiiiiiiii iiiiiiiiii |
| PolyPhobius | oooooooooM MMMMMMMMMM MMMMMMMMMM iiiiiiiiii iiiiiiiiii |
| SCAMPI |  |
| SPOCTOPUS | oooooooooo MMMMMMMMMM MMMMMMMMMM Mooooooooo oooooooooo |
| PDB-homology |  |
|  | 351                                         391 |
| Seq. | PHCSGDIIRQ IDFFENYLKS TPMQFTVLGI SITYQKLGGF LFVIFELSAA |
| TOPCONS | oooooooooo oooooooooo oooooooooo oooooooooo oooooooooo |
| OCTOPUS |  |
| Philius | iiiiiiiiii iiiiiiiiii iiiiiiiiii iiiiiiMMMM MMMMMMMMMM |
| PolyPhobius | iiiiiiiiii iiiiiiiiii iiiiiiiiii iiiiiiMMMM MMMMMMMMMM |
| SCAMPI |  |
| SPOCTOPUS | oooooooooo oooooooooo oooooooooo MMMMMMMMMM MMMMMMMMMM |
| PDB-homology |  |

|  |  |
| --- | --- |
|  | 401 |
| Seq. | VLIKSFHNEV |
| TOPCONS | oooooooooo |
| OCTOPUS |  |
| Philius | MMMooooooo |
| PolyPhobius | MMMMMMoooo |
| SCAMPI |  |
| SPOCTOPUS | Mooooooooo |
| PDB-homology |  |
