## Supplementary figures and images for "A putative origin of insect chemosensory receptors in the last common eukaryotic ancestor"

### topcons.large.png

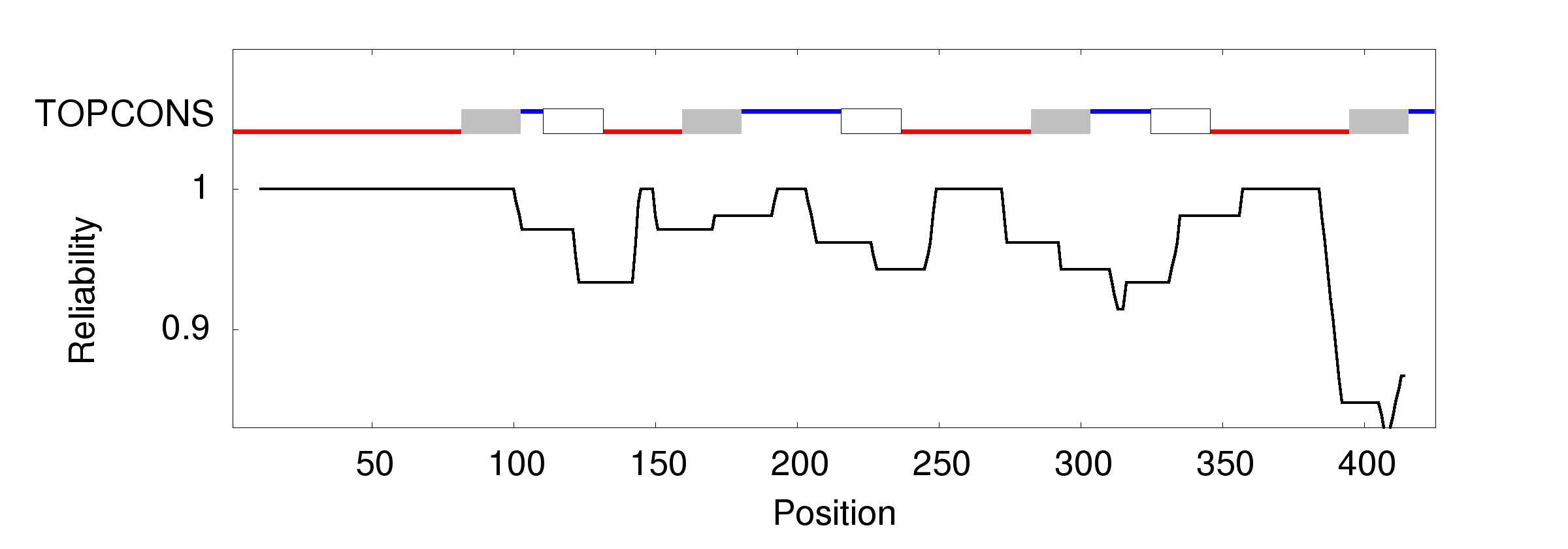

### topcons.large.png

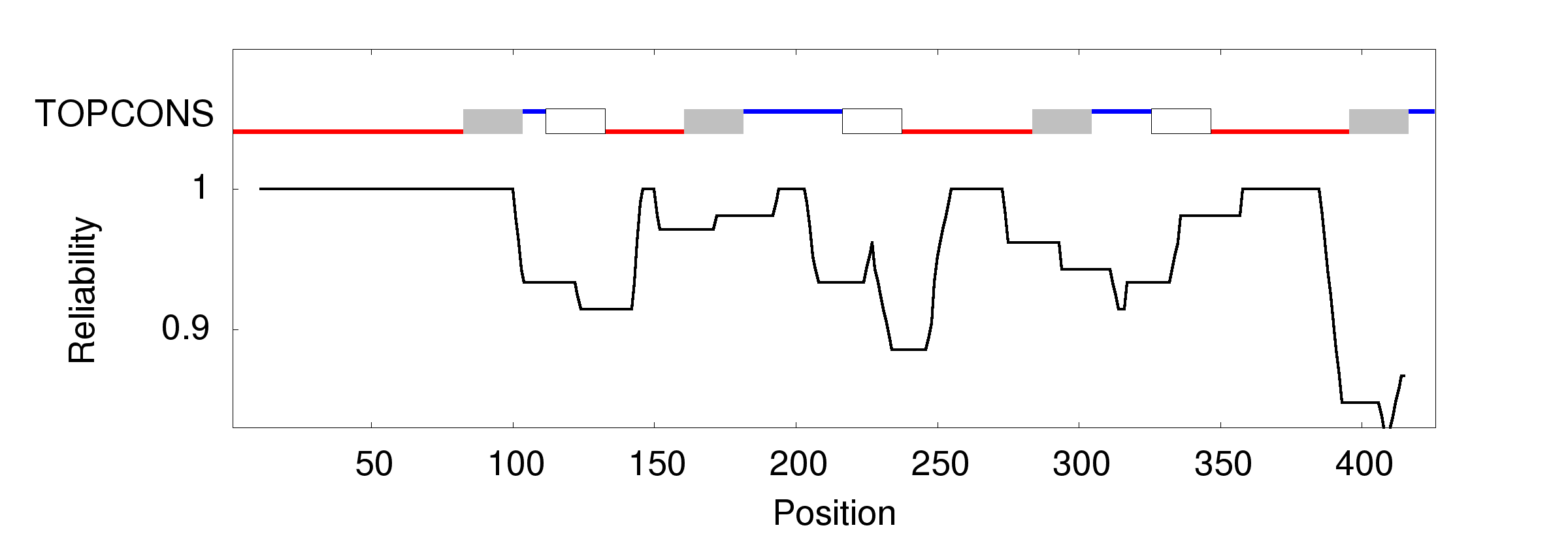

### topcons.large.png

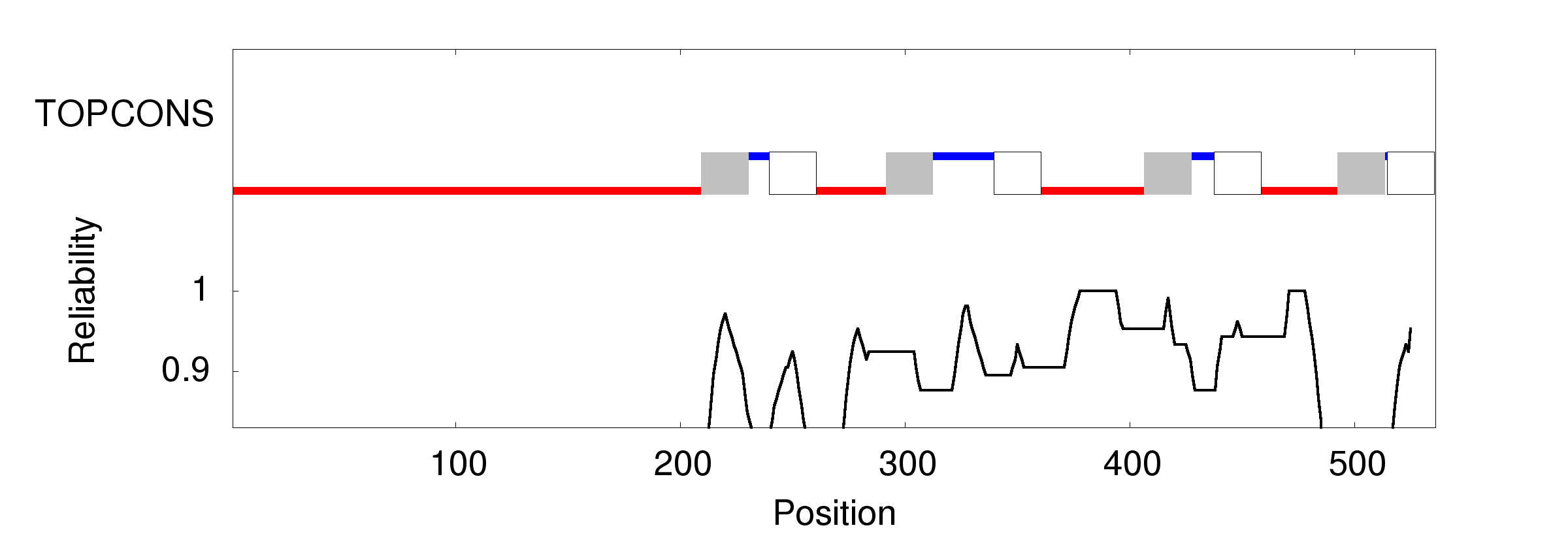

### topcons.large.png

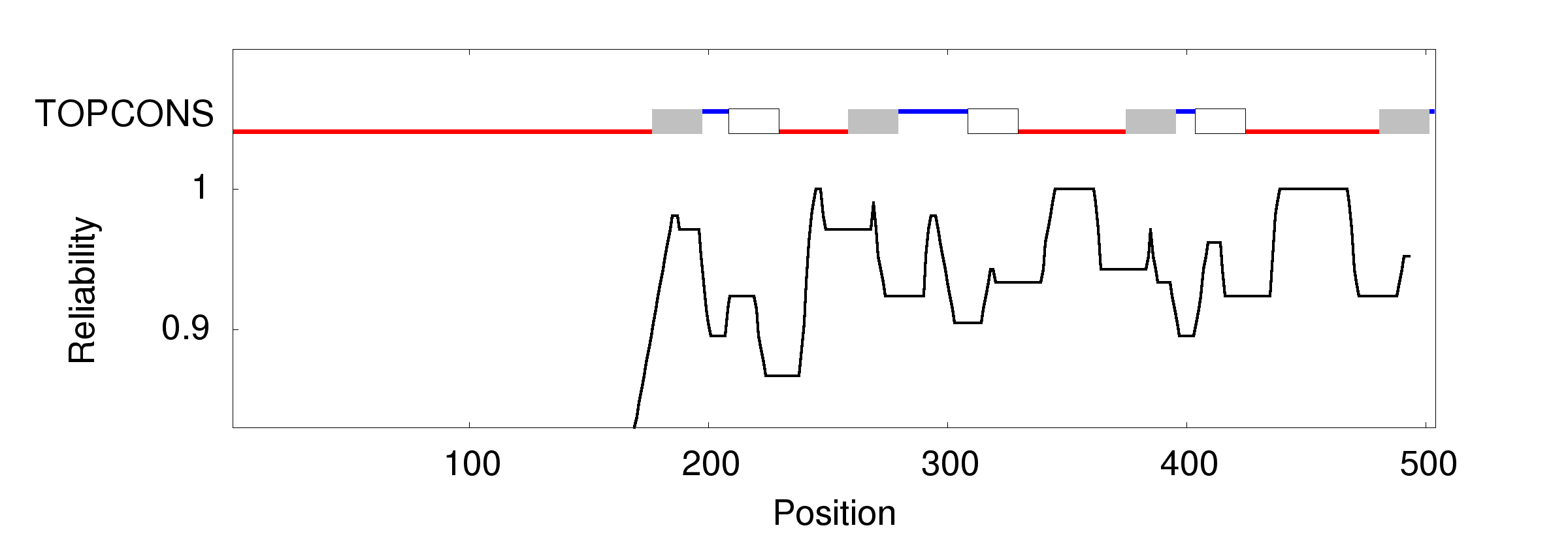

### topcons.large.png

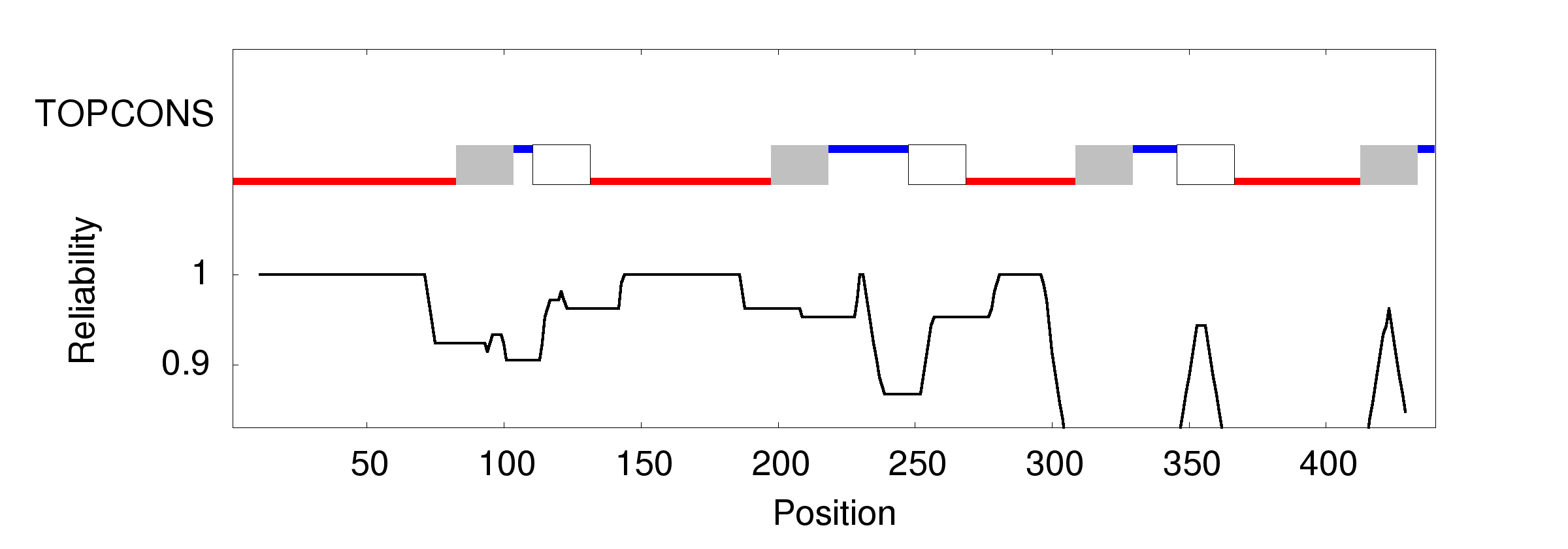

### topcons.large.png

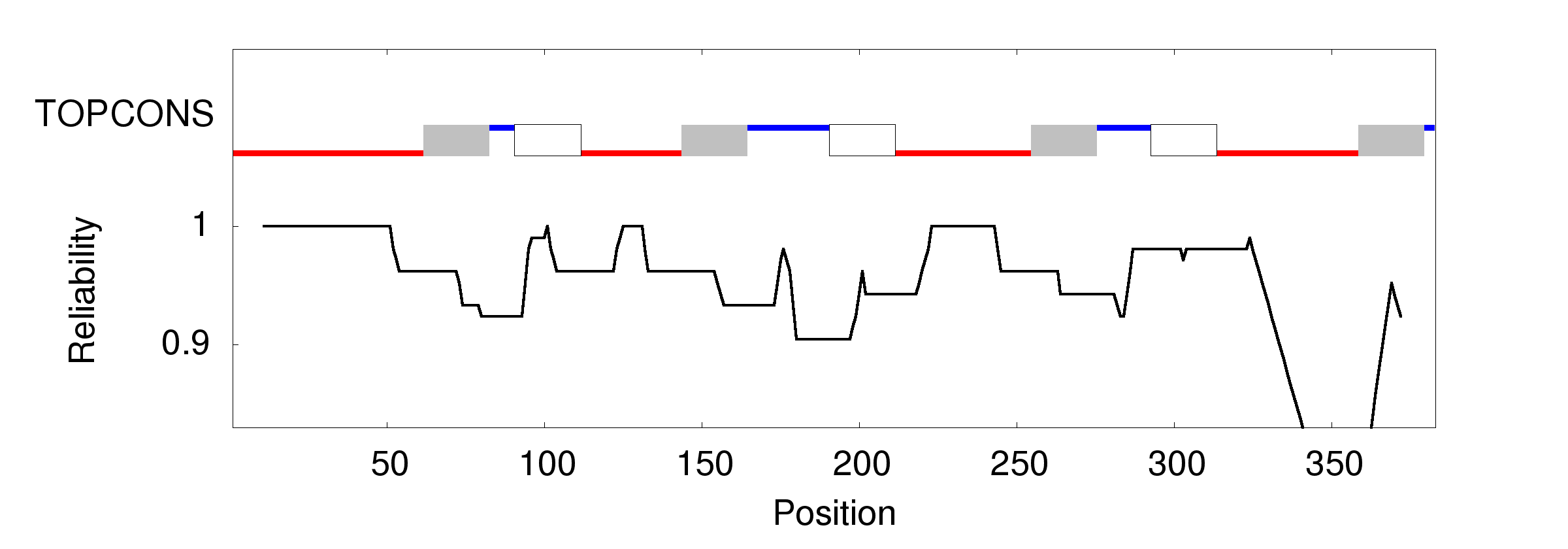

### topcons.large.png

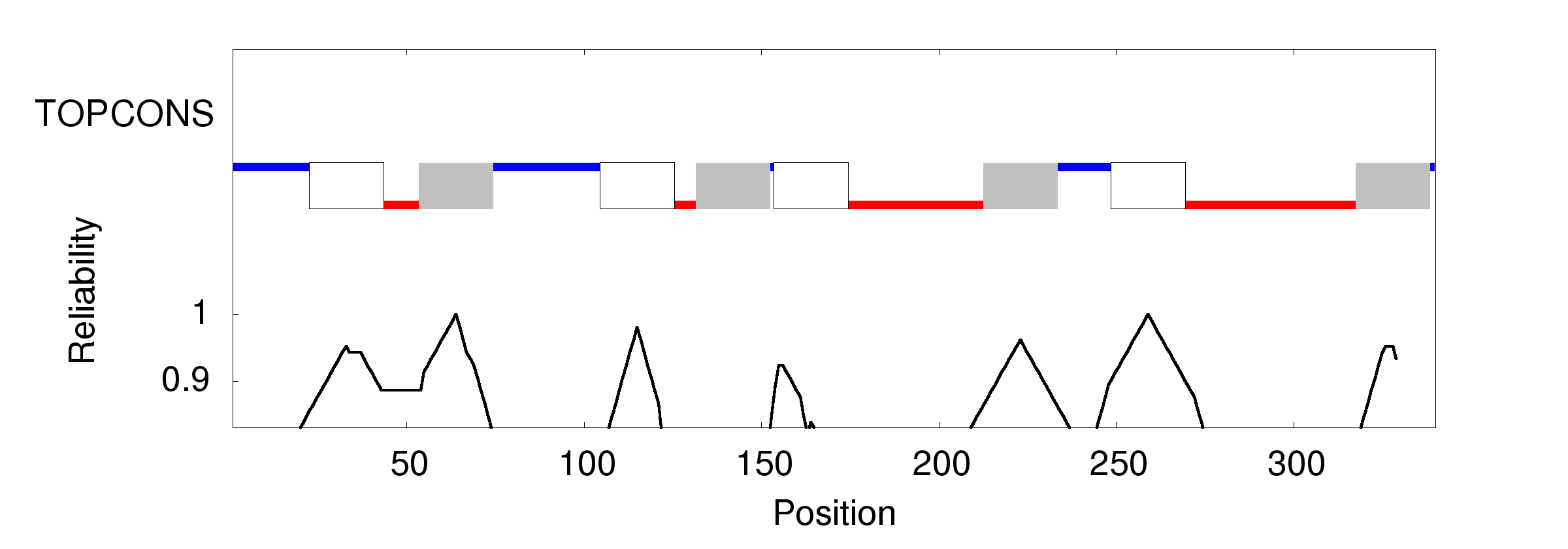

### topcons.large.png

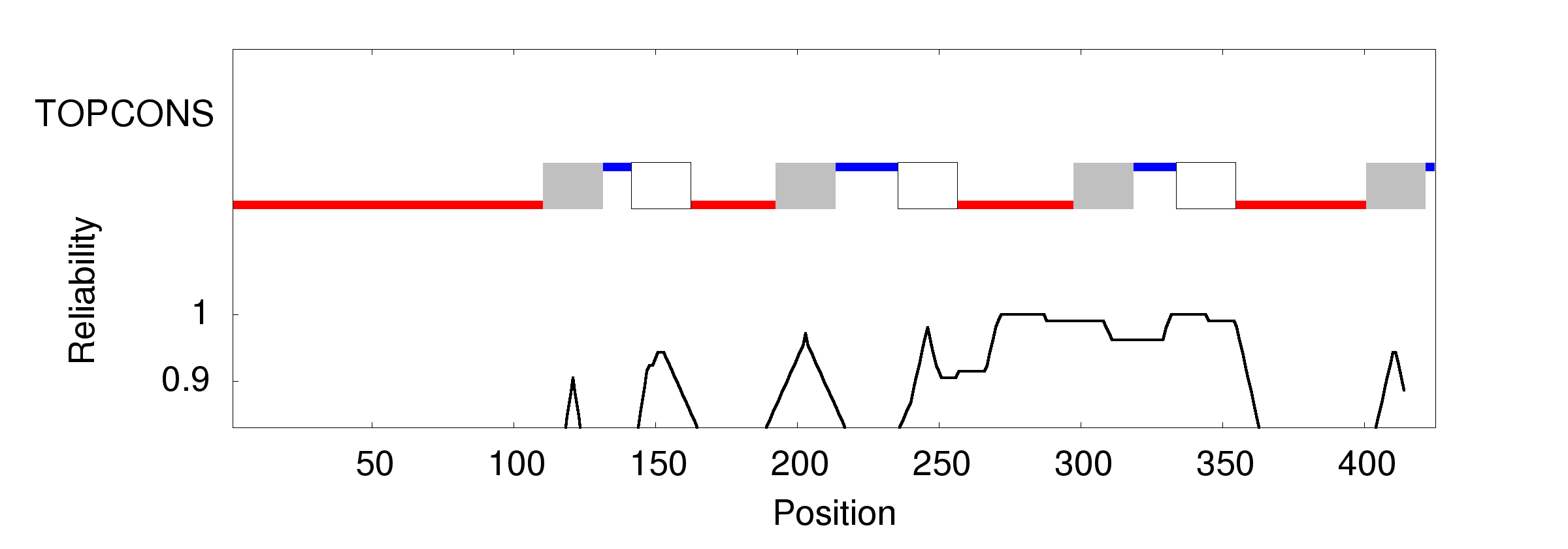

### topcons.png

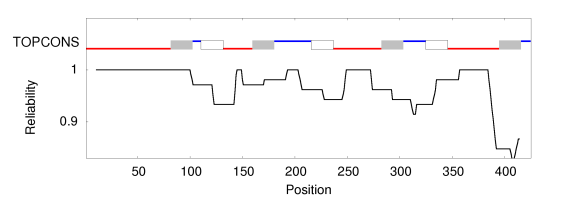

### topcons.png

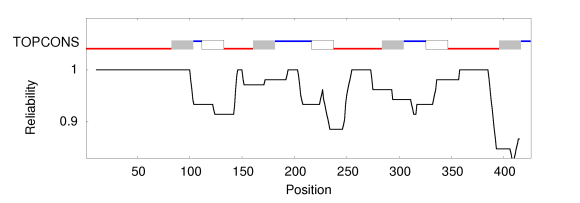

### topcons.png

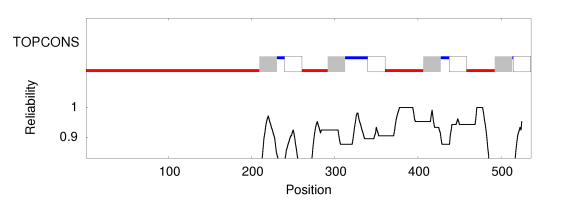

### topcons.png

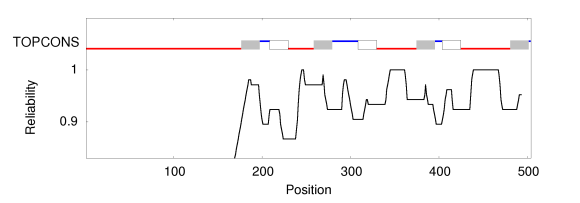

### topcons.png

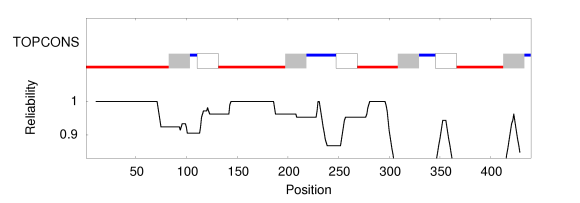

### topcons.png

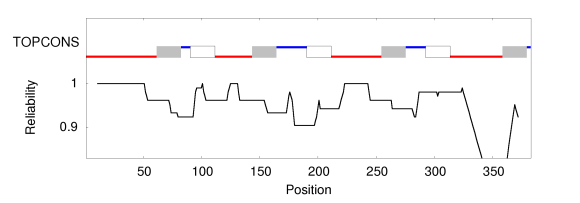

### topcons.png

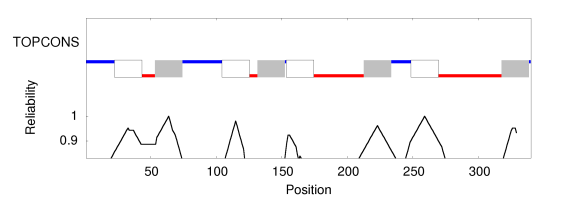

### topcons.png

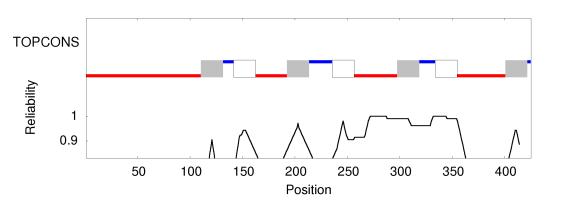

### total_image.large.png

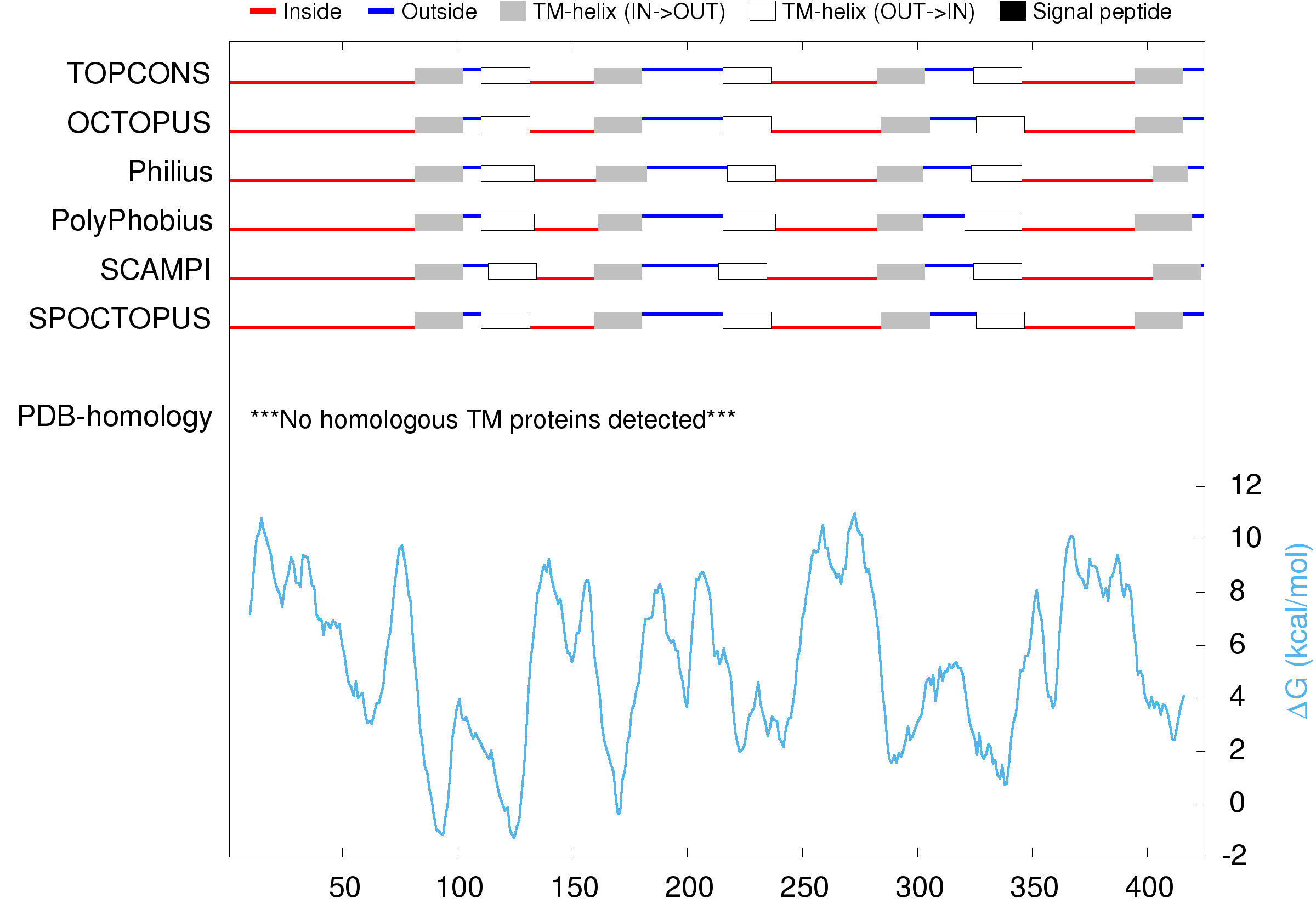

### total_image.large.png

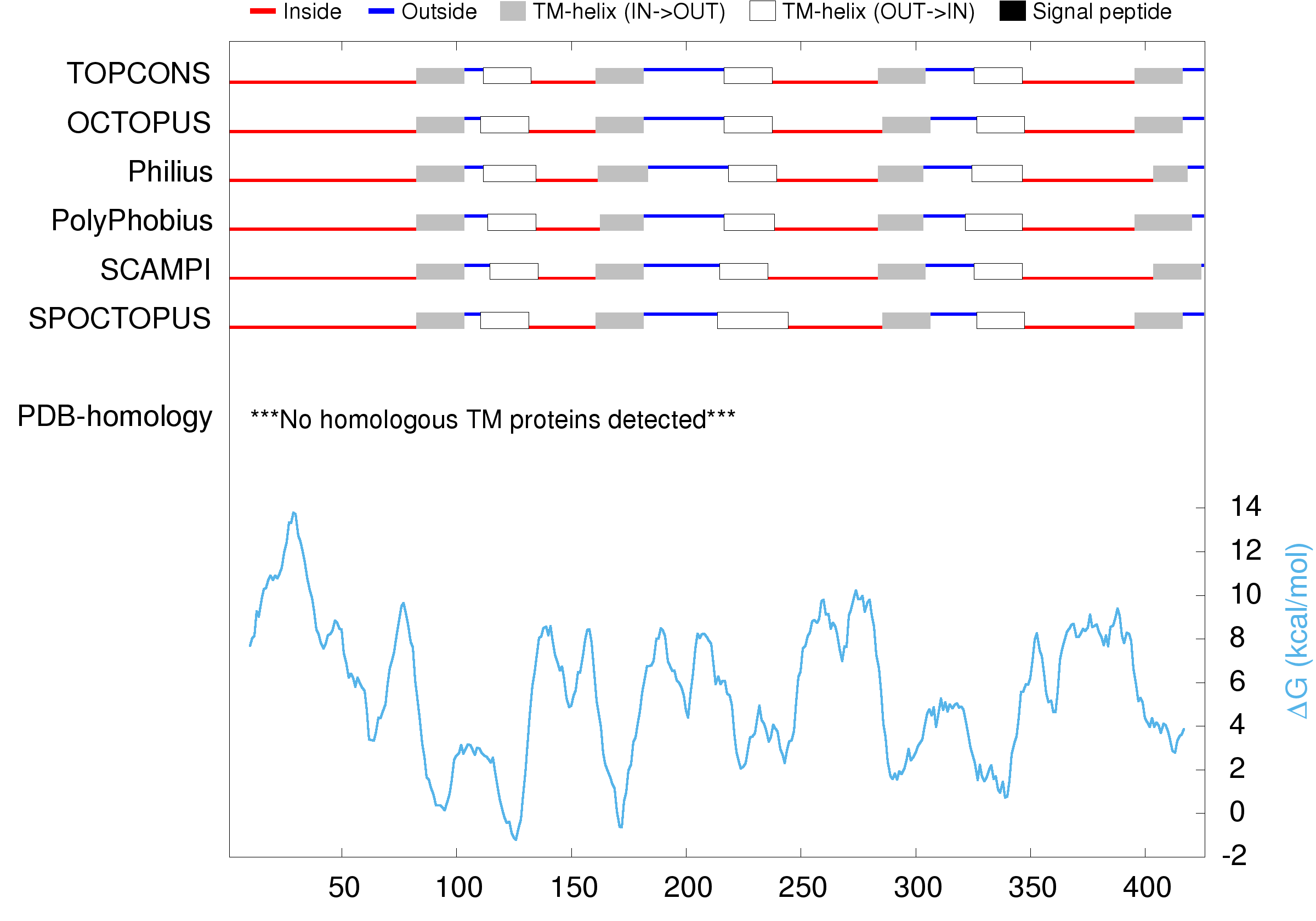

### total_image.large.png

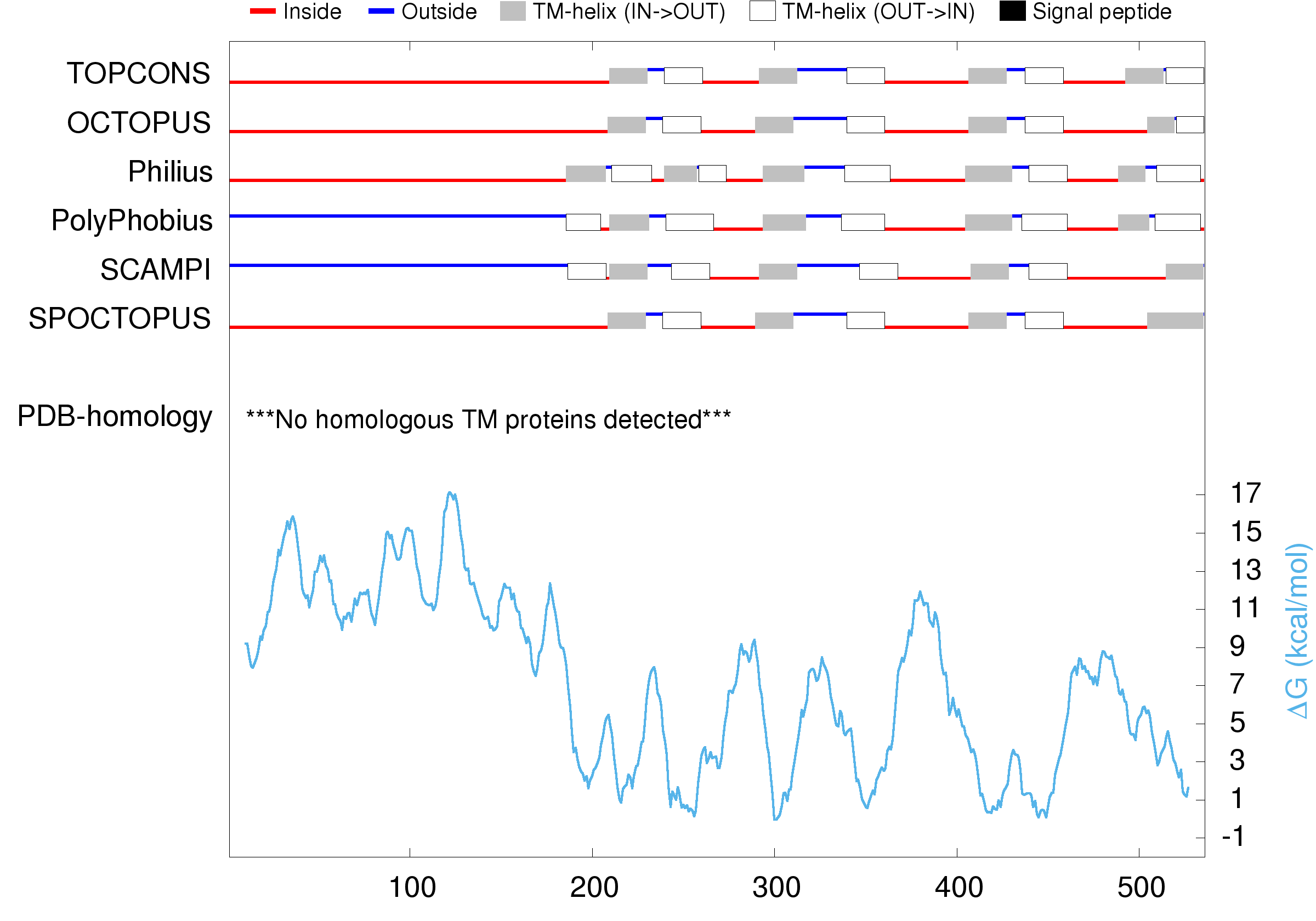

### total_image.large.png

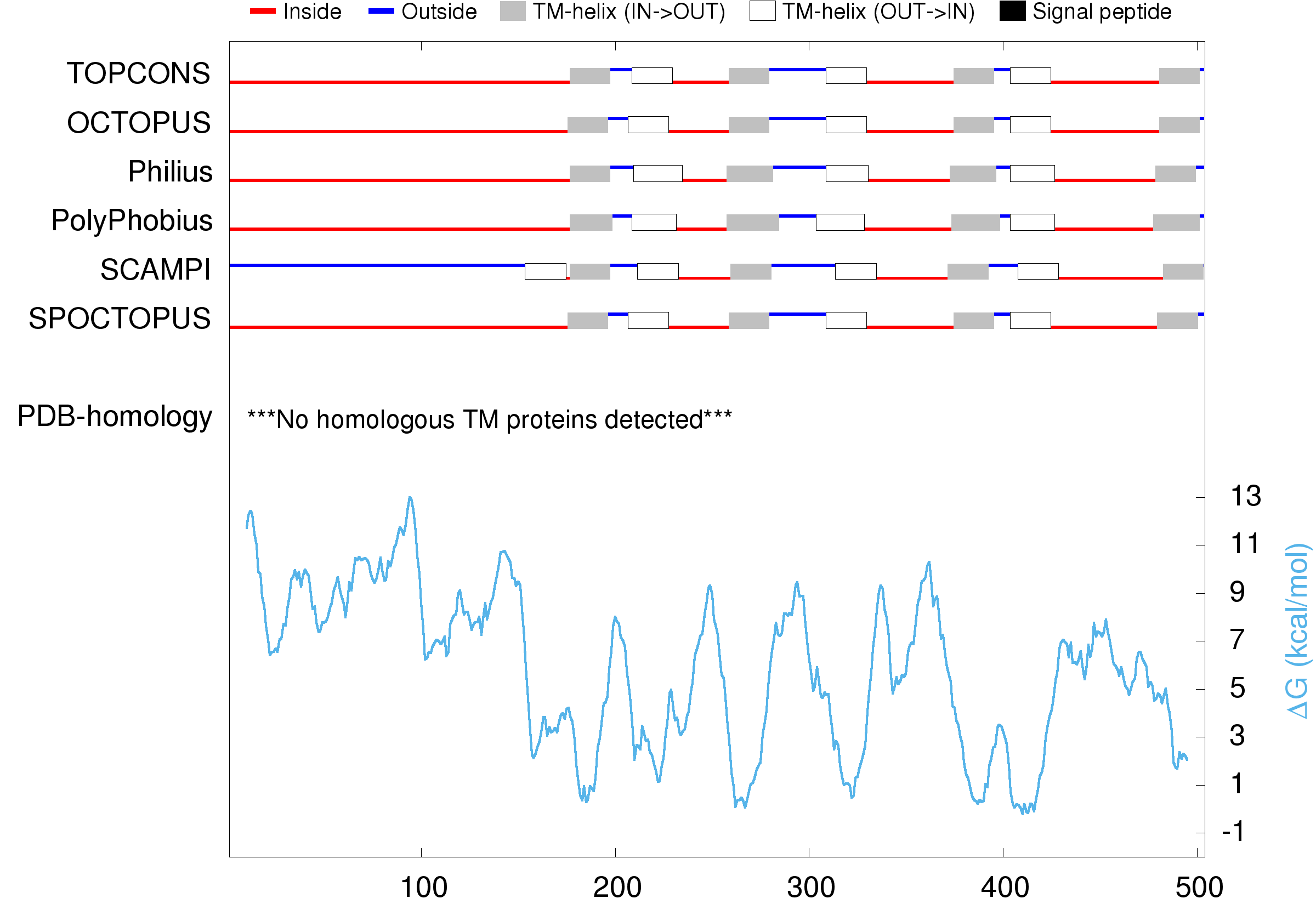

### total_image.large.png

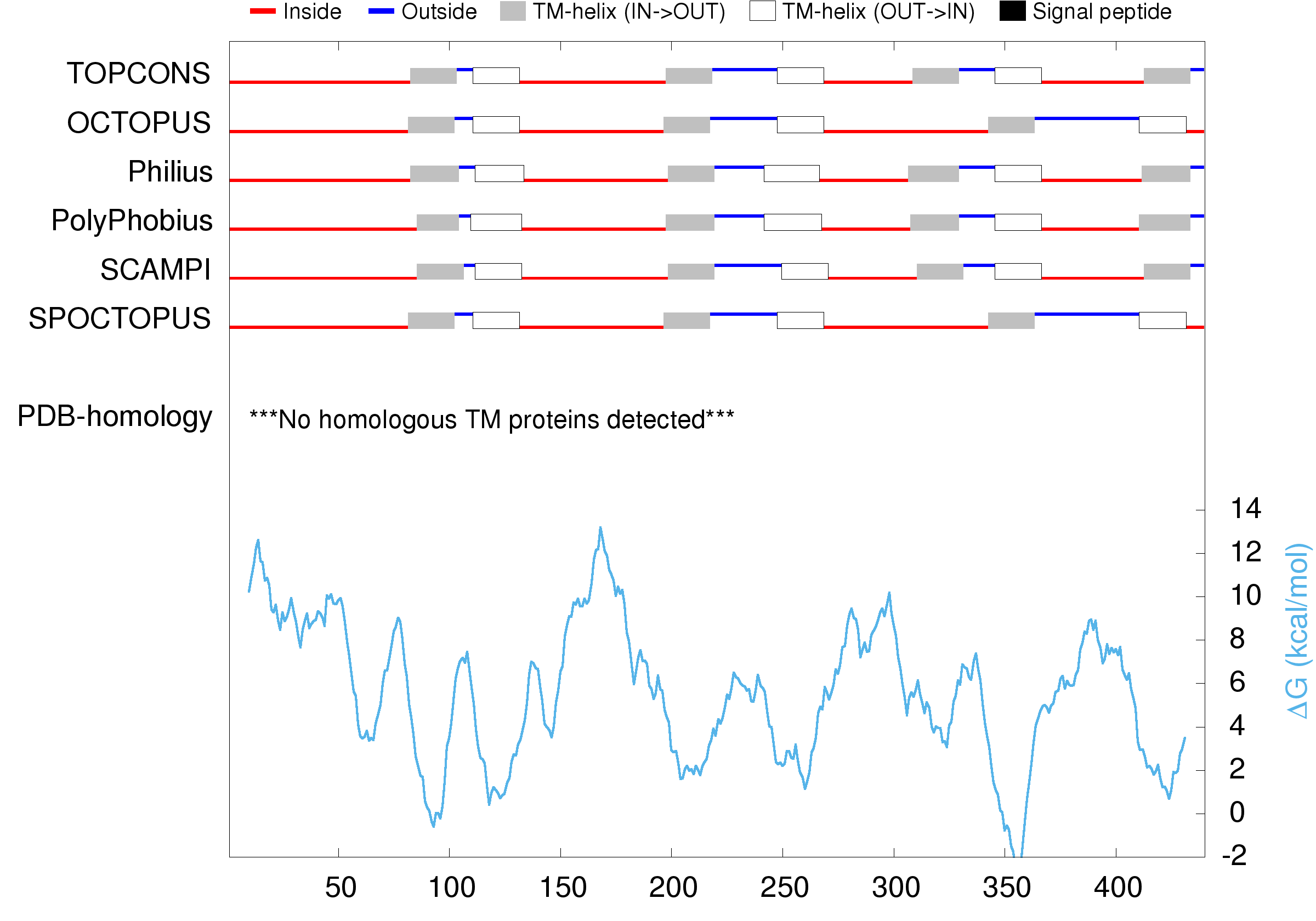

### total_image.large.png

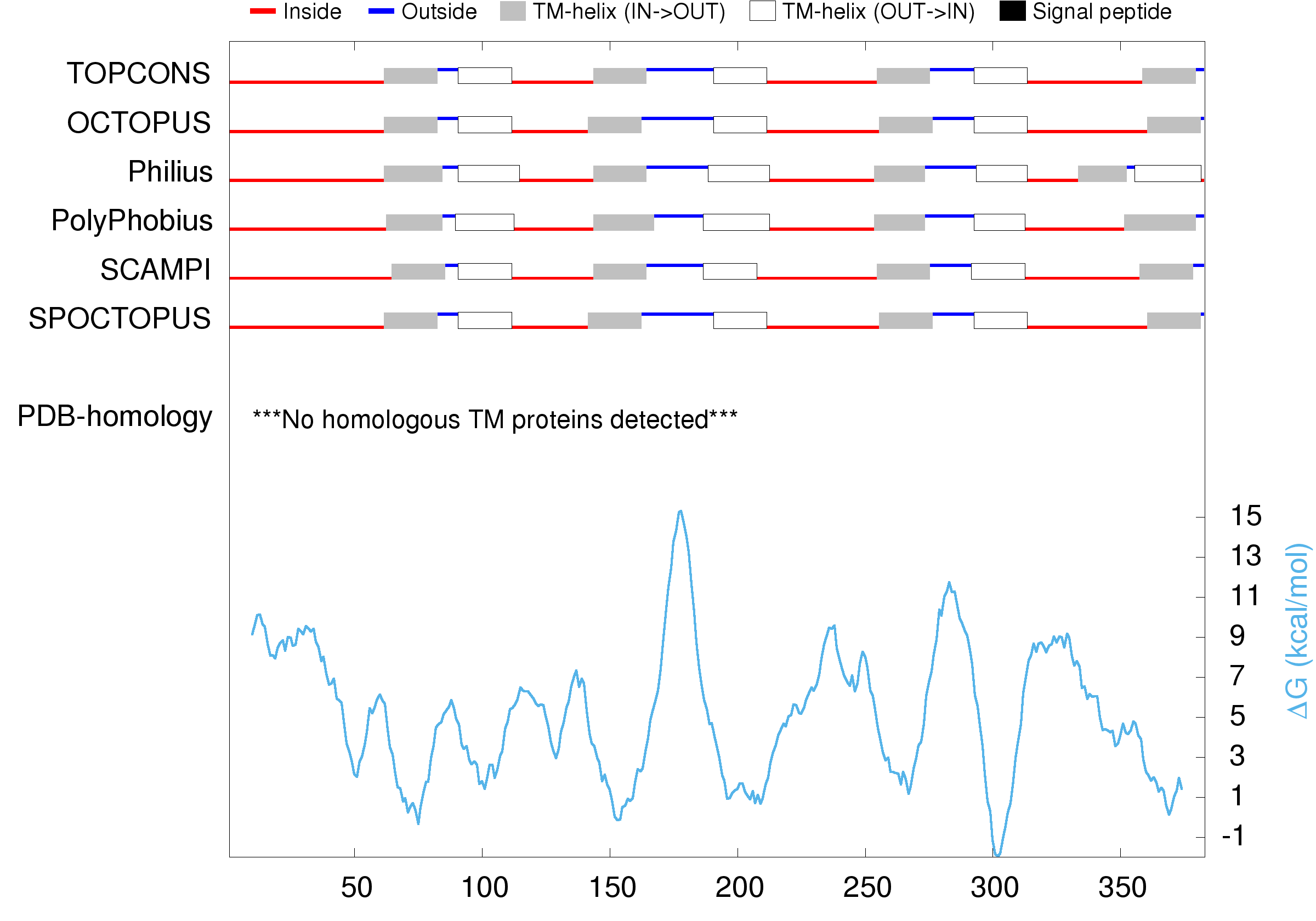

### total_image.large.png

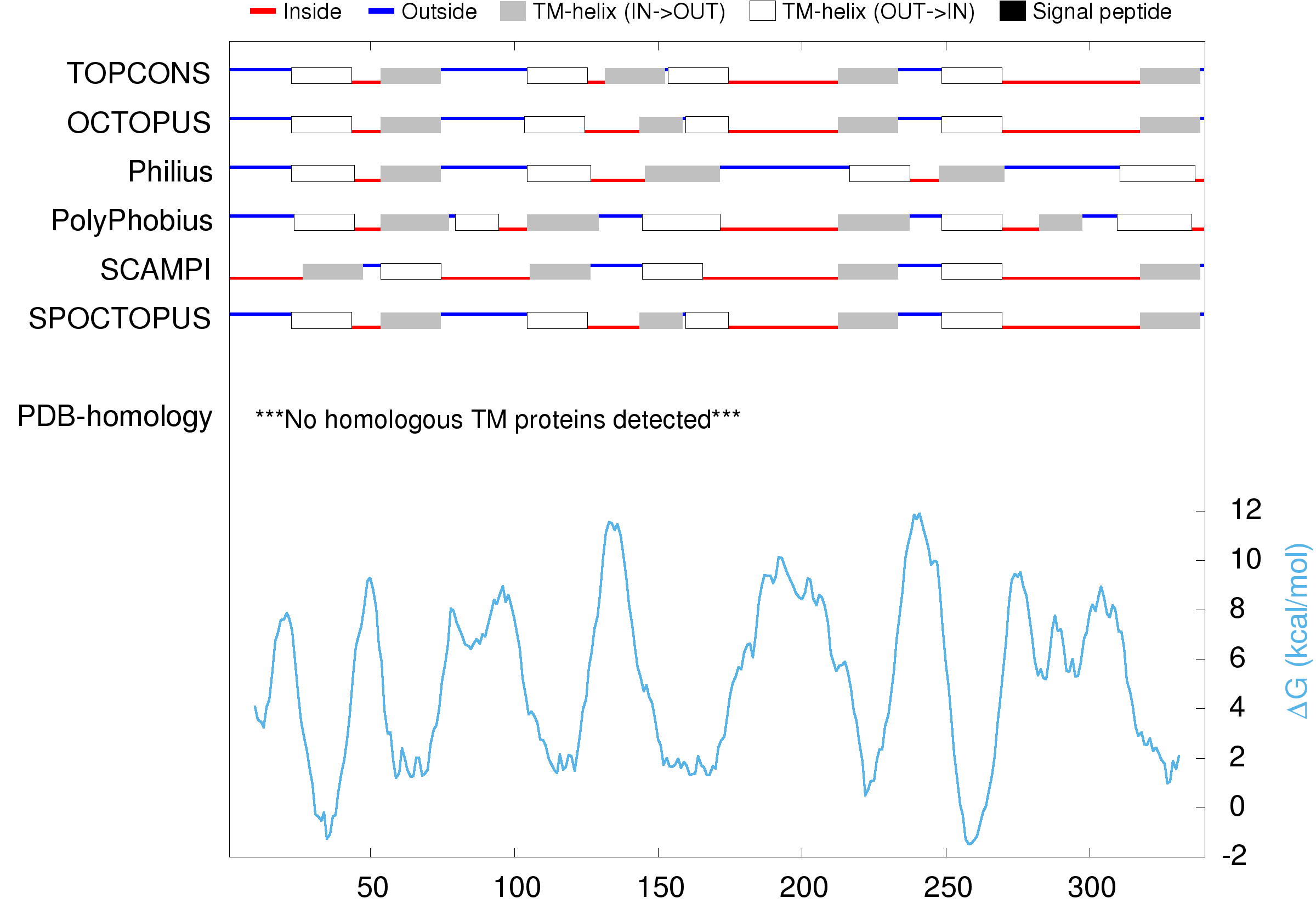

### total_image.large.png

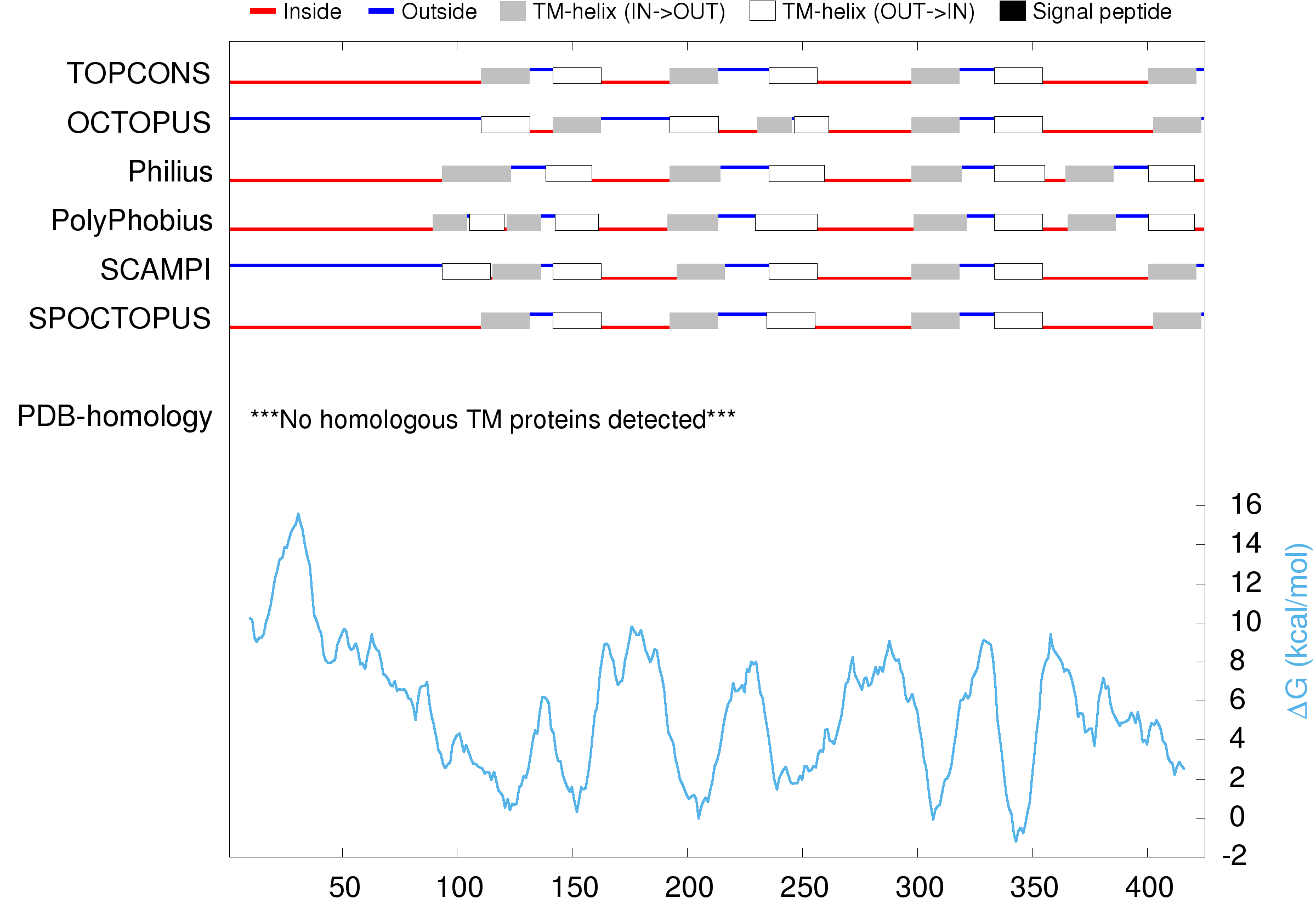

### total_image.png

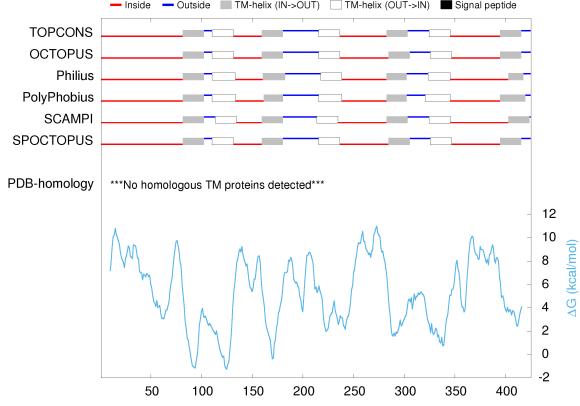

### total_image.png

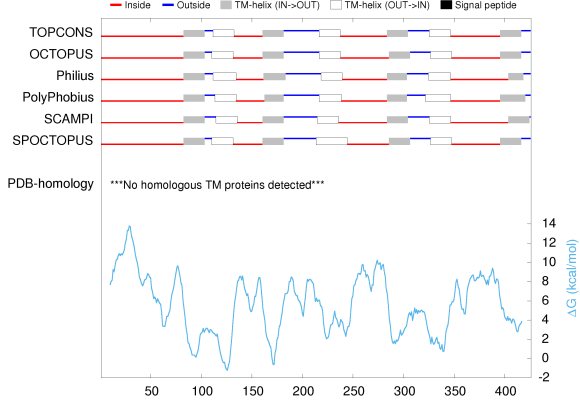

### total_image.png

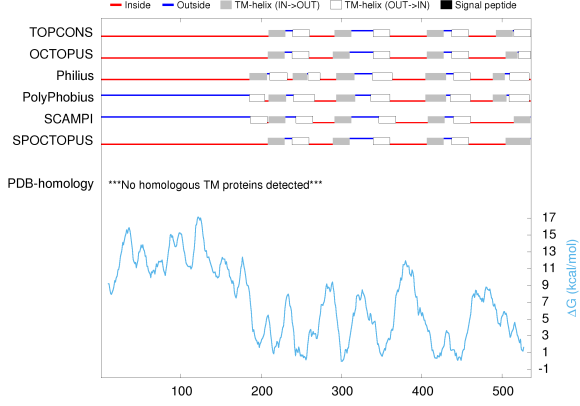

### total_image.png

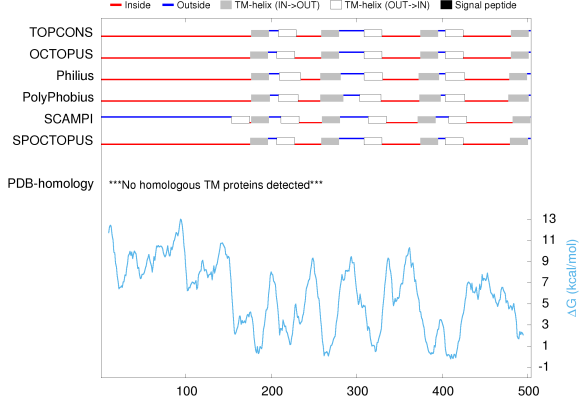

### total_image.png

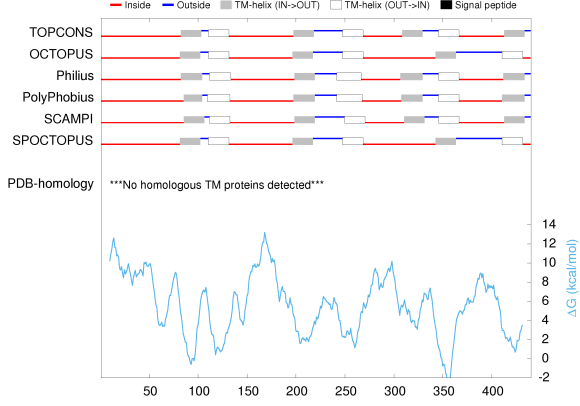

### total_image.png

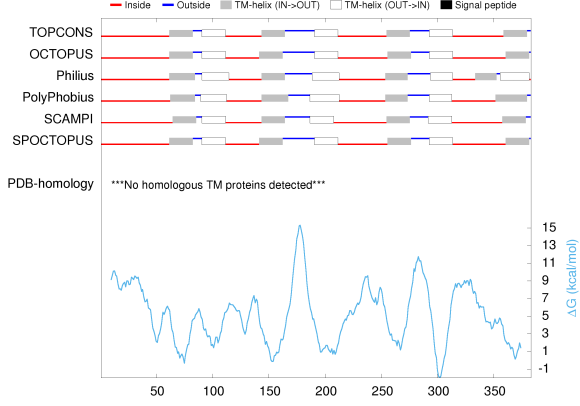

### total_image.png

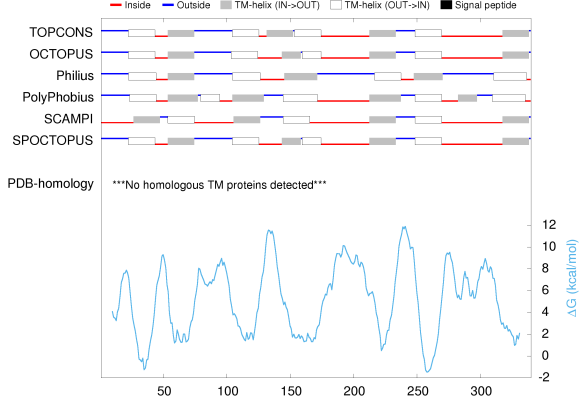

### total_image.png

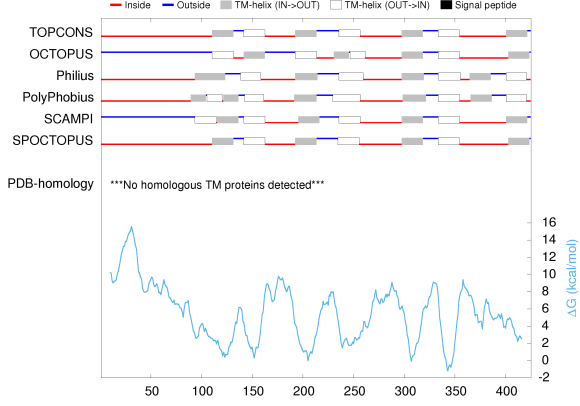
