## Supplementary material for "A putative origin of insect chemosensory receptors in the last common eukaryotic ancestor": Data S4: 200821_DataS4_code.html

hhblitsphyl


### Python notebook to obtain the HMM-based probability distance matrix¶

In [1]:

```
# load libraries
import pandas as pd
import hhsuitedb as hhdb
import glob, os, subprocess, shlex
from Bio import SeqIO
import csb
from csb.bio.io.hhpred import HHOutputParser
from matplotlib import pyplot as plt
import numpy as np
import networkx as nx
import pickle
import tempfile
from ete3 import PhyloTree
import seaborn as sns
```

In [3]:

```
# Path to the UniClust database
uniclust = '/home/cactuskid13/mntpt/HHBLITsdb/uniclust30_2018_08/uniclust30_2018_08'

qdir = 'orco/'
runName ='ORCO'
ncores = 8
# path to all query files
queries = glob.glob(qdir + 'queries/*.fasta')
```

In [4]:

```
# Functions to run programs from the HHsuite software
def runHHblits( aln , name, path , outdir, db , iterations , ncores , runName='' , SS= False  , ohhm = False , verbose = True , Z = 2000 , B = 2000 , xargs = ''):
    if verbose == True:
        print( [aln , name, path , outdir, db , iterations , ncores , runName] )
    
    outhhr= outdir+name+runName+".hhr"
    args = path + ' -cpu '+ str(ncores) +' -d ' + db + ' -i ' + aln  +' -o '+ outhhr + ' -n ' + str(iterations) + ' -B '+ str(B) + ' -Z ' + str(Z) +' '+ xargs 
    if SS == True:
         args += ' -ssm 2 -ssw .5 '
    
    if ohhm == True:
        outa3m = outdir+name+runName+'.hhm'
        args += ' -ohhm ' + outa3m
    else: 
        outa3m = None
    if verbose == True:
        print(args)
    
    args = shlex.split( args)
    p = subprocess.run( args )
    return p , [outhhr,outa3m]


def runHHmake( aln , name, path = 'hhmake' , outdir='./', verbose = False, SS = False):
    if verbose == True:
        print( [aln , name, path , outdir] )
    outhhm= outdir+name+".hhm"
    args = path + ' -i '+  aln  +' -o '+ outhhm + ' -M 50'
    if SS == True:
        #todo : make ss prediction here
        pass
    args = shlex.split(args)
    print(args)
    p = subprocess.Popen(args )
    return p , [outhhm]

    

def hhrparse(hhr , coverage , proba ):
    profile = HHOutputParser(alignments=False).parse_file(hhr)
    qname = profile.query_name
    for hit in profile:
        proba = hit.probability
        i = hit.id
```

In [8]:

```
# create output directories, as well as query files
queries = glob.glob(qdir + '*.fasta')
if not os.path.exists(qdir +'queries'):
    os.mkdir(qdir + 'queries')
if not os.path.exists(qdir +'HHfiles'):
    os.mkdir(qdir + 'HHfiles')
if not os.path.exists(qdir +'ALLVSALL'):
    os.mkdir(qdir + 'ALLVSALL')


for qfile in queries:
    print(qfile)
    fasta = SeqIO.parse(qfile, 'fasta')
    for s in fasta:
        SeqIO.write( [s]  ,  qdir + 'queries/' + s.id + '.fasta' , 'fasta' )
```

```
orco/orco.fasta
orco/orco_clst7.fasta
```

In [10]:

```
# run HHblits on each query
print(queries)
for q in queries:
    p,output = runHHblits(q , name = q.split('.')[0].split('/')[-1] + 'Profphylo' , path= 'hhblits ' , outdir = qdir+'HHfiles/' , db = uniclust , iterations= 3 , ncores = ncores , ohhm = True, verbose = True , runName=runName, xargs = ' -mact .5')
```

```
['orco/queries/AthaAT4G22270.fasta', 'orco/queries/AthaAT1G67570.fasta', 'orco/queries/DmelGr64a.fasta', 'orco/queries/MpusGRL1.fasta', 'orco/queries/VbraGRL5.fasta', 'orco/queries/TtraGRL4.fasta', 'orco/queries/TtraGRL6.fasta', 'orco/queries/SkowGRL1.fasta', 'orco/queries/TadhGRL1.fasta', 'orco/queries/AthaAT2G21080.fasta', 'orco/queries/VbraGRL4.fasta', 'orco/queries/AbakORCO.fasta', 'orco/queries/NvecGRL1.fasta', 'orco/queries/TtraGRL5.fasta', 'orco/queries/VbraGRL6.fasta', 'orco/queries/SpurGRL1.fasta', 'orco/queries/AthaAT4G03820.fasta', 'orco/queries/CpriGRL1.fasta', 'orco/queries/VbraGRL1.fasta', 'orco/queries/SpunGRL1.fasta', 'orco/queries/PfunGRL1.fasta', 'orco/queries/TtraGRL2.fasta', 'orco/queries/AthaAT1G50630.fasta', 'orco/queries/AthaAT3G20300.fasta', 'orco/queries/TtraGRL1.fasta', 'orco/queries/TtraGRL3.fasta', 'orco/queries/VbraGRL2.fasta', 'orco/queries/SpalGRL1.fasta', 'orco/queries/VbraGRL3.fasta']
['orco/queries/AthaAT4G22270.fasta', 'AthaAT4G22270Profphylo', 'hhblits ', 'orco/HHfiles/', '/home/cactuskid13/mntpt/HHBLITsdb/uniclust30_2018_08/uniclust30_2018_08', 3, 8, 'ORCO']
hhblits  -cpu 8 -d /home/cactuskid13/mntpt/HHBLITsdb/uniclust30_2018_08/uniclust30_2018_08 -i orco/queries/AthaAT4G22270.fasta -o orco/HHfiles/AthaAT4G22270ProfphyloORCO.hhr -n 3 -B 2000 -Z 2000  -mact .5 -ohhm orco/HHfiles/AthaAT4G22270ProfphyloORCO.hhm
['orco/queries/AthaAT1G67570.fasta', 'AthaAT1G67570Profphylo', 'hhblits ', 'orco/HHfiles/', '/home/cactuskid13/mntpt/HHBLITsdb/uniclust30_2018_08/uniclust30_2018_08', 3, 8, 'ORCO']
hhblits  -cpu 8 -d /home/cactuskid13/mntpt/HHBLITsdb/uniclust30_2018_08/uniclust30_2018_08 -i orco/queries/AthaAT1G67570.fasta -o orco/HHfiles/AthaAT1G67570ProfphyloORCO.hhr -n 3 -B 2000 -Z 2000  -mact .5 -ohhm orco/HHfiles/AthaAT1G67570ProfphyloORCO.hhm
['orco/queries/DmelGr64a.fasta', 'DmelGr64aProfphylo', 'hhblits ', 'orco/HHfiles/', '/home/cactuskid13/mntpt/HHBLITsdb/uniclust30_2018_08/uniclust30_2018_08', 3, 8, 'ORCO']
hhblits  -cpu 8 -d /home/cactuskid13/mntpt/HHBLITsdb/uniclust30_2018_08/uniclust30_2018_08 -i orco/queries/DmelGr64a.fasta -o orco/HHfiles/DmelGr64aProfphyloORCO.hhr -n 3 -B 2000 -Z 2000  -mact .5 -ohhm orco/HHfiles/DmelGr64aProfphyloORCO.hhm
['orco/queries/MpusGRL1.fasta', 'MpusGRL1Profphylo', 'hhblits ', 'orco/HHfiles/', '/home/cactuskid13/mntpt/HHBLITsdb/uniclust30_2018_08/uniclust30_2018_08', 3, 8, 'ORCO']
hhblits  -cpu 8 -d /home/cactuskid13/mntpt/HHBLITsdb/uniclust30_2018_08/uniclust30_2018_08 -i orco/queries/MpusGRL1.fasta -o orco/HHfiles/MpusGRL1ProfphyloORCO.hhr -n 3 -B 2000 -Z 2000  -mact .5 -ohhm orco/HHfiles/MpusGRL1ProfphyloORCO.hhm
['orco/queries/VbraGRL5.fasta', 'VbraGRL5Profphylo', 'hhblits ', 'orco/HHfiles/', '/home/cactuskid13/mntpt/HHBLITsdb/uniclust30_2018_08/uniclust30_2018_08', 3, 8, 'ORCO']
hhblits  -cpu 8 -d /home/cactuskid13/mntpt/HHBLITsdb/uniclust30_2018_08/uniclust30_2018_08 -i orco/queries/VbraGRL5.fasta -o orco/HHfiles/VbraGRL5ProfphyloORCO.hhr -n 3 -B 2000 -Z 2000  -mact .5 -ohhm orco/HHfiles/VbraGRL5ProfphyloORCO.hhm
['orco/queries/TtraGRL4.fasta', 'TtraGRL4Profphylo', 'hhblits ', 'orco/HHfiles/', '/home/cactuskid13/mntpt/HHBLITsdb/uniclust30_2018_08/uniclust30_2018_08', 3, 8, 'ORCO']
hhblits  -cpu 8 -d /home/cactuskid13/mntpt/HHBLITsdb/uniclust30_2018_08/uniclust30_2018_08 -i orco/queries/TtraGRL4.fasta -o orco/HHfiles/TtraGRL4ProfphyloORCO.hhr -n 3 -B 2000 -Z 2000  -mact .5 -ohhm orco/HHfiles/TtraGRL4ProfphyloORCO.hhm
['orco/queries/TtraGRL6.fasta', 'TtraGRL6Profphylo', 'hhblits ', 'orco/HHfiles/', '/home/cactuskid13/mntpt/HHBLITsdb/uniclust30_2018_08/uniclust30_2018_08', 3, 8, 'ORCO']
hhblits  -cpu 8 -d /home/cactuskid13/mntpt/HHBLITsdb/uniclust30_2018_08/uniclust30_2018_08 -i orco/queries/TtraGRL6.fasta -o orco/HHfiles/TtraGRL6ProfphyloORCO.hhr -n 3 -B 2000 -Z 2000  -mact .5 -ohhm orco/HHfiles/TtraGRL6ProfphyloORCO.hhm
['orco/queries/SkowGRL1.fasta', 'SkowGRL1Profphylo', 'hhblits ', 'orco/HHfiles/', '/home/cactuskid13/mntpt/HHBLITsdb/uniclust30_2018_08/uniclust30_2018_08', 3, 8, 'ORCO']
hhblits  -cpu 8 -d /home/cactuskid13/mntpt/HHBLITsdb/uniclust30_2018_08/uniclust30_2018_08 -i orco/queries/SkowGRL1.fasta -o orco/HHfiles/SkowGRL1ProfphyloORCO.hhr -n 3 -B 2000 -Z 2000  -mact .5 -ohhm orco/HHfiles/SkowGRL1ProfphyloORCO.hhm
['orco/queries/TadhGRL1.fasta', 'TadhGRL1Profphylo', 'hhblits ', 'orco/HHfiles/', '/home/cactuskid13/mntpt/HHBLITsdb/uniclust30_2018_08/uniclust30_2018_08', 3, 8, 'ORCO']
hhblits  -cpu 8 -d /home/cactuskid13/mntpt/HHBLITsdb/uniclust30_2018_08/uniclust30_2018_08 -i orco/queries/TadhGRL1.fasta -o orco/HHfiles/TadhGRL1ProfphyloORCO.hhr -n 3 -B 2000 -Z 2000  -mact .5 -ohhm orco/HHfiles/TadhGRL1ProfphyloORCO.hhm
['orco/queries/AthaAT2G21080.fasta', 'AthaAT2G21080Profphylo', 'hhblits ', 'orco/HHfiles/', '/home/cactuskid13/mntpt/HHBLITsdb/uniclust30_2018_08/uniclust30_2018_08', 3, 8, 'ORCO']
hhblits  -cpu 8 -d /home/cactuskid13/mntpt/HHBLITsdb/uniclust30_2018_08/uniclust30_2018_08 -i orco/queries/AthaAT2G21080.fasta -o orco/HHfiles/AthaAT2G21080ProfphyloORCO.hhr -n 3 -B 2000 -Z 2000  -mact .5 -ohhm orco/HHfiles/AthaAT2G21080ProfphyloORCO.hhm
['orco/queries/VbraGRL4.fasta', 'VbraGRL4Profphylo', 'hhblits ', 'orco/HHfiles/', '/home/cactuskid13/mntpt/HHBLITsdb/uniclust30_2018_08/uniclust30_2018_08', 3, 8, 'ORCO']
hhblits  -cpu 8 -d /home/cactuskid13/mntpt/HHBLITsdb/uniclust30_2018_08/uniclust30_2018_08 -i orco/queries/VbraGRL4.fasta -o orco/HHfiles/VbraGRL4ProfphyloORCO.hhr -n 3 -B 2000 -Z 2000  -mact .5 -ohhm orco/HHfiles/VbraGRL4ProfphyloORCO.hhm
['orco/queries/AbakORCO.fasta', 'AbakORCOProfphylo', 'hhblits ', 'orco/HHfiles/', '/home/cactuskid13/mntpt/HHBLITsdb/uniclust30_2018_08/uniclust30_2018_08', 3, 8, 'ORCO']
hhblits  -cpu 8 -d /home/cactuskid13/mntpt/HHBLITsdb/uniclust30_2018_08/uniclust30_2018_08 -i orco/queries/AbakORCO.fasta -o orco/HHfiles/AbakORCOProfphyloORCO.hhr -n 3 -B 2000 -Z 2000  -mact .5 -ohhm orco/HHfiles/AbakORCOProfphyloORCO.hhm
['orco/queries/NvecGRL1.fasta', 'NvecGRL1Profphylo', 'hhblits ', 'orco/HHfiles/', '/home/cactuskid13/mntpt/HHBLITsdb/uniclust30_2018_08/uniclust30_2018_08', 3, 8, 'ORCO']
hhblits  -cpu 8 -d /home/cactuskid13/mntpt/HHBLITsdb/uniclust30_2018_08/uniclust30_2018_08 -i orco/queries/NvecGRL1.fasta -o orco/HHfiles/NvecGRL1ProfphyloORCO.hhr -n 3 -B 2000 -Z 2000  -mact .5 -ohhm orco/HHfiles/NvecGRL1ProfphyloORCO.hhm
['orco/queries/TtraGRL5.fasta', 'TtraGRL5Profphylo', 'hhblits ', 'orco/HHfiles/', '/home/cactuskid13/mntpt/HHBLITsdb/uniclust30_2018_08/uniclust30_2018_08', 3, 8, 'ORCO']
hhblits  -cpu 8 -d /home/cactuskid13/mntpt/HHBLITsdb/uniclust30_2018_08/uniclust30_2018_08 -i orco/queries/TtraGRL5.fasta -o orco/HHfiles/TtraGRL5ProfphyloORCO.hhr -n 3 -B 2000 -Z 2000  -mact .5 -ohhm orco/HHfiles/TtraGRL5ProfphyloORCO.hhm
['orco/queries/VbraGRL6.fasta', 'VbraGRL6Profphylo', 'hhblits ', 'orco/HHfiles/', '/home/cactuskid13/mntpt/HHBLITsdb/uniclust30_2018_08/uniclust30_2018_08', 3, 8, 'ORCO']
hhblits  -cpu 8 -d /home/cactuskid13/mntpt/HHBLITsdb/uniclust30_2018_08/uniclust30_2018_08 -i orco/queries/VbraGRL6.fasta -o orco/HHfiles/VbraGRL6ProfphyloORCO.hhr -n 3 -B 2000 -Z 2000  -mact .5 -ohhm orco/HHfiles/VbraGRL6ProfphyloORCO.hhm
['orco/queries/SpurGRL1.fasta', 'SpurGRL1Profphylo', 'hhblits ', 'orco/HHfiles/', '/home/cactuskid13/mntpt/HHBLITsdb/uniclust30_2018_08/uniclust30_2018_08', 3, 8, 'ORCO']
hhblits  -cpu 8 -d /home/cactuskid13/mntpt/HHBLITsdb/uniclust30_2018_08/uniclust30_2018_08 -i orco/queries/SpurGRL1.fasta -o orco/HHfiles/SpurGRL1ProfphyloORCO.hhr -n 3 -B 2000 -Z 2000  -mact .5 -ohhm orco/HHfiles/SpurGRL1ProfphyloORCO.hhm
['orco/queries/AthaAT4G03820.fasta', 'AthaAT4G03820Profphylo', 'hhblits ', 'orco/HHfiles/', '/home/cactuskid13/mntpt/HHBLITsdb/uniclust30_2018_08/uniclust30_2018_08', 3, 8, 'ORCO']
hhblits  -cpu 8 -d /home/cactuskid13/mntpt/HHBLITsdb/uniclust30_2018_08/uniclust30_2018_08 -i orco/queries/AthaAT4G03820.fasta -o orco/HHfiles/AthaAT4G03820ProfphyloORCO.hhr -n 3 -B 2000 -Z 2000  -mact .5 -ohhm orco/HHfiles/AthaAT4G03820ProfphyloORCO.hhm
['orco/queries/CpriGRL1.fasta', 'CpriGRL1Profphylo', 'hhblits ', 'orco/HHfiles/', '/home/cactuskid13/mntpt/HHBLITsdb/uniclust30_2018_08/uniclust30_2018_08', 3, 8, 'ORCO']
hhblits  -cpu 8 -d /home/cactuskid13/mntpt/HHBLITsdb/uniclust30_2018_08/uniclust30_2018_08 -i orco/queries/CpriGRL1.fasta -o orco/HHfiles/CpriGRL1ProfphyloORCO.hhr -n 3 -B 2000 -Z 2000  -mact .5 -ohhm orco/HHfiles/CpriGRL1ProfphyloORCO.hhm
['orco/queries/VbraGRL1.fasta', 'VbraGRL1Profphylo', 'hhblits ', 'orco/HHfiles/', '/home/cactuskid13/mntpt/HHBLITsdb/uniclust30_2018_08/uniclust30_2018_08', 3, 8, 'ORCO']
hhblits  -cpu 8 -d /home/cactuskid13/mntpt/HHBLITsdb/uniclust30_2018_08/uniclust30_2018_08 -i orco/queries/VbraGRL1.fasta -o orco/HHfiles/VbraGRL1ProfphyloORCO.hhr -n 3 -B 2000 -Z 2000  -mact .5 -ohhm orco/HHfiles/VbraGRL1ProfphyloORCO.hhm
['orco/queries/SpunGRL1.fasta', 'SpunGRL1Profphylo', 'hhblits ', 'orco/HHfiles/', '/home/cactuskid13/mntpt/HHBLITsdb/uniclust30_2018_08/uniclust30_2018_08', 3, 8, 'ORCO']
hhblits  -cpu 8 -d /home/cactuskid13/mntpt/HHBLITsdb/uniclust30_2018_08/uniclust30_2018_08 -i orco/queries/SpunGRL1.fasta -o orco/HHfiles/SpunGRL1ProfphyloORCO.hhr -n 3 -B 2000 -Z 2000  -mact .5 -ohhm orco/HHfiles/SpunGRL1ProfphyloORCO.hhm
['orco/queries/PfunGRL1.fasta', 'PfunGRL1Profphylo', 'hhblits ', 'orco/HHfiles/', '/home/cactuskid13/mntpt/HHBLITsdb/uniclust30_2018_08/uniclust30_2018_08', 3, 8, 'ORCO']
hhblits  -cpu 8 -d /home/cactuskid13/mntpt/HHBLITsdb/uniclust30_2018_08/uniclust30_2018_08 -i orco/queries/PfunGRL1.fasta -o orco/HHfiles/PfunGRL1ProfphyloORCO.hhr -n 3 -B 2000 -Z 2000  -mact .5 -ohhm orco/HHfiles/PfunGRL1ProfphyloORCO.hhm
['orco/queries/TtraGRL2.fasta', 'TtraGRL2Profphylo', 'hhblits ', 'orco/HHfiles/', '/home/cactuskid13/mntpt/HHBLITsdb/uniclust30_2018_08/uniclust30_2018_08', 3, 8, 'ORCO']
hhblits  -cpu 8 -d /home/cactuskid13/mntpt/HHBLITsdb/uniclust30_2018_08/uniclust30_2018_08 -i orco/queries/TtraGRL2.fasta -o orco/HHfiles/TtraGRL2ProfphyloORCO.hhr -n 3 -B 2000 -Z 2000  -mact .5 -ohhm orco/HHfiles/TtraGRL2ProfphyloORCO.hhm
['orco/queries/AthaAT1G50630.fasta', 'AthaAT1G50630Profphylo', 'hhblits ', 'orco/HHfiles/', '/home/cactuskid13/mntpt/HHBLITsdb/uniclust30_2018_08/uniclust30_2018_08', 3, 8, 'ORCO']
hhblits  -cpu 8 -d /home/cactuskid13/mntpt/HHBLITsdb/uniclust30_2018_08/uniclust30_2018_08 -i orco/queries/AthaAT1G50630.fasta -o orco/HHfiles/AthaAT1G50630ProfphyloORCO.hhr -n 3 -B 2000 -Z 2000  -mact .5 -ohhm orco/HHfiles/AthaAT1G50630ProfphyloORCO.hhm
['orco/queries/AthaAT3G20300.fasta', 'AthaAT3G20300Profphylo', 'hhblits ', 'orco/HHfiles/', '/home/cactuskid13/mntpt/HHBLITsdb/uniclust30_2018_08/uniclust30_2018_08', 3, 8, 'ORCO']
hhblits  -cpu 8 -d /home/cactuskid13/mntpt/HHBLITsdb/uniclust30_2018_08/uniclust30_2018_08 -i orco/queries/AthaAT3G20300.fasta -o orco/HHfiles/AthaAT3G20300ProfphyloORCO.hhr -n 3 -B 2000 -Z 2000  -mact .5 -ohhm orco/HHfiles/AthaAT3G20300ProfphyloORCO.hhm
['orco/queries/TtraGRL1.fasta', 'TtraGRL1Profphylo', 'hhblits ', 'orco/HHfiles/', '/home/cactuskid13/mntpt/HHBLITsdb/uniclust30_2018_08/uniclust30_2018_08', 3, 8, 'ORCO']
hhblits  -cpu 8 -d /home/cactuskid13/mntpt/HHBLITsdb/uniclust30_2018_08/uniclust30_2018_08 -i orco/queries/TtraGRL1.fasta -o orco/HHfiles/TtraGRL1ProfphyloORCO.hhr -n 3 -B 2000 -Z 2000  -mact .5 -ohhm orco/HHfiles/TtraGRL1ProfphyloORCO.hhm
['orco/queries/TtraGRL3.fasta', 'TtraGRL3Profphylo', 'hhblits ', 'orco/HHfiles/', '/home/cactuskid13/mntpt/HHBLITsdb/uniclust30_2018_08/uniclust30_2018_08', 3, 8, 'ORCO']
hhblits  -cpu 8 -d /home/cactuskid13/mntpt/HHBLITsdb/uniclust30_2018_08/uniclust30_2018_08 -i orco/queries/TtraGRL3.fasta -o orco/HHfiles/TtraGRL3ProfphyloORCO.hhr -n 3 -B 2000 -Z 2000  -mact .5 -ohhm orco/HHfiles/TtraGRL3ProfphyloORCO.hhm
['orco/queries/VbraGRL2.fasta', 'VbraGRL2Profphylo', 'hhblits ', 'orco/HHfiles/', '/home/cactuskid13/mntpt/HHBLITsdb/uniclust30_2018_08/uniclust30_2018_08', 3, 8, 'ORCO']
hhblits  -cpu 8 -d /home/cactuskid13/mntpt/HHBLITsdb/uniclust30_2018_08/uniclust30_2018_08 -i orco/queries/VbraGRL2.fasta -o orco/HHfiles/VbraGRL2ProfphyloORCO.hhr -n 3 -B 2000 -Z 2000  -mact .5 -ohhm orco/HHfiles/VbraGRL2ProfphyloORCO.hhm
['orco/queries/SpalGRL1.fasta', 'SpalGRL1Profphylo', 'hhblits ', 'orco/HHfiles/', '/home/cactuskid13/mntpt/HHBLITsdb/uniclust30_2018_08/uniclust30_2018_08', 3, 8, 'ORCO']
hhblits  -cpu 8 -d /home/cactuskid13/mntpt/HHBLITsdb/uniclust30_2018_08/uniclust30_2018_08 -i orco/queries/SpalGRL1.fasta -o orco/HHfiles/SpalGRL1ProfphyloORCO.hhr -n 3 -B 2000 -Z 2000  -mact .5 -ohhm orco/HHfiles/SpalGRL1ProfphyloORCO.hhm
['orco/queries/VbraGRL3.fasta', 'VbraGRL3Profphylo', 'hhblits ', 'orco/HHfiles/', '/home/cactuskid13/mntpt/HHBLITsdb/uniclust30_2018_08/uniclust30_2018_08', 3, 8, 'ORCO']
hhblits  -cpu 8 -d /home/cactuskid13/mntpt/HHBLITsdb/uniclust30_2018_08/uniclust30_2018_08 -i orco/queries/VbraGRL3.fasta -o orco/HHfiles/VbraGRL3ProfphyloORCO.hhr -n 3 -B 2000 -Z 2000  -mact .5 -ohhm orco/HHfiles/VbraGRL3ProfphyloORCO.hhm
```

In [18]:

```
# We can assign an HHblits score to each hit

newhits = {}

hhfiles = glob.glob(qdir+'HHfiles/*.hhr')
print(hhfiles)

for file in hhfiles:
    results = HHOutputParser(alignments=False).parse_file(file)
    for hit in results:
        newhits.update( { hit.id.split('|')[1].strip() : { 'prob':hit.probability, 'len' : hit.length , 'query':results.query_name }}  )
df = pd.DataFrame.from_dict(newhits, orient='index')                                                      

#adhoc score = len * proba
df['score'] = df['prob'] * df['len']

#sort the data frame and print
print(df.sort_values( 'score' , ascending = False) )
df.to_csv( './orco/orco_hhblits_results.csv' )
```

```
['orco/HHfiles/NvecGRL1ProfphyloORCO.hhr', 'orco/HHfiles/AthaAT4G22270ProfphyloORCO.hhr', 'orco/HHfiles/SpurGRL1ProfphyloORCO.hhr', 'orco/HHfiles/VbraGRL6ProfphyloORCO.hhr', 'orco/HHfiles/VbraGRL2ProfphyloORCO.hhr', 'orco/HHfiles/TtraGRL5ProfphyloORCO.hhr', 'orco/HHfiles/AthaAT1G50630ProfphyloORCO.hhr', 'orco/HHfiles/AbakORCOProfphyloORCO.hhr', 'orco/HHfiles/TadhGRL1ProfphyloORCO.hhr', 'orco/HHfiles/CpriGRL1ProfphyloORCO.hhr', 'orco/HHfiles/VbraGRL1ProfphyloORCO.hhr', 'orco/HHfiles/TtraGRL2ProfphyloORCO.hhr', 'orco/HHfiles/VbraGRL5ProfphyloORCO.hhr', 'orco/HHfiles/MpusGRL1ProfphyloORCO.hhr', 'orco/HHfiles/SkowGRL1ProfphyloORCO.hhr', 'orco/HHfiles/VbraGRL4ProfphyloORCO.hhr', 'orco/HHfiles/PfunGRL1ProfphyloORCO.hhr', 'orco/HHfiles/TtraGRL3ProfphyloORCO.hhr', 'orco/HHfiles/SpalGRL1ProfphyloORCO.hhr', 'orco/HHfiles/AthaAT2G21080ProfphyloORCO.hhr', 'orco/HHfiles/TtraGRL4ProfphyloORCO.hhr', 'orco/HHfiles/AthaAT1G67570ProfphyloORCO.hhr', 'orco/HHfiles/AthaAT3G20300ProfphyloORCO.hhr', 'orco/HHfiles/SpunGRL1ProfphyloORCO.hhr', 'orco/HHfiles/TtraGRL1ProfphyloORCO.hhr', 'orco/HHfiles/TtraGRL6ProfphyloORCO.hhr', 'orco/HHfiles/AthaAT4G03820ProfphyloORCO.hhr', 'orco/HHfiles/DmelGr64aProfphyloORCO.hhr', 'orco/HHfiles/VbraGRL3ProfphyloORCO.hhr']
             prob    len                                              query  \
A0A0G4FWI7  1.000    730  VbraGRL3 CEM19221.1 unnamed protein product [V...   
A0A0L0DUY0  1.000    536  TtraGRL1 gi|923135227|ref|XP_013761079.1| hypo...   
B3NG15      1.000    450  DmelGr64a NP_728920.1 gustatory receptor 64a [...   
A0A0L0D5B5  1.000    437  TtraGRL3 gi|923132535|ref|XP_013759733.1| hypo...   
A0A194YRL2  1.000    431  AthaAT4G03820 Athaliana|AT4G03820|AT4G03820.1/...   
...           ...    ...                                                ...   
A0A0L8H0D7  0.227      6  AthaAT2G21080 Athaliana|AT2G21080|AT2G21080.1/...   
A0A2V6BG25  0.216      6  VbraGRL5 CEM10760.1 unnamed protein product [V...   
S8EV43      0.216      6  VbraGRL3 CEM19221.1 unnamed protein product [V...   
A0A0V0YA82  0.215      6  AthaAT1G67570 Athaliana|AT1G67570|AT1G67570.1/...   
K8F3H1_0    0.205 -12659  VbraGRL3 CEM19221.1 unnamed protein product [V...   

               score  
A0A0G4FWI7   730.000  
A0A0L0DUY0   536.000  
B3NG15       450.000  
A0A0L0D5B5   437.000  
A0A194YRL2   431.000  
...              ...  
A0A0L8H0D7     1.362  
A0A2V6BG25     1.296  
S8EV43         1.296  
A0A0V0YA82     1.290  
K8F3H1_0   -2595.095  

[7377 rows x 4 columns]
```

In [29]:

```
#plot the scores and lengths of hits
newhitsdf = pd.DataFrame.from_dict(newhits, orient= 'index')
newhitsdf.hist()
plt.show()

#filter out poor matches
filterdf = newhitsdf[newhitsdf.prob >.9]
filterdf = filterdf[newhitsdf.len >200]

filterdf.hist()
plt.show()

print(filterdf)
```

```
/home/cactuskid13/miniconda3/envs/pyprofiler3/lib/python3.7/site-packages/ipykernel_launcher.py:9: UserWarning: Boolean Series key will be reindexed to match DataFrame index.
  if __name__ == '__main__':
```

```
             prob  len                                              query
A0A0C5GSD5  0.992  205  DmelGr64a NP_728920.1 gustatory receptor 64a [...
A0A2B4RET4  0.991  204  DmelGr64a NP_728920.1 gustatory receptor 64a [...
B3NG15      1.000  450  DmelGr64a NP_728920.1 gustatory receptor 64a [...
J9LZ04      1.000  359  DmelGr64a NP_728920.1 gustatory receptor 64a [...
A0A1W4WHQ3  1.000  292  DmelGr64a NP_728920.1 gustatory receptor 64a [...
...           ...  ...                                                ...
A0A151JNA6  0.983  204   AbakORCO tr|B0FAQ4|B0FAQ4_APOBA Odorant receptor
A0A1I8NAF8  0.983  210   AbakORCO tr|B0FAQ4|B0FAQ4_APOBA Odorant receptor
A0A0L0D5B5  1.000  437  TtraGRL3 gi|923132535|ref|XP_013759733.1| hypo...
A0A2P6P0A7  1.000  410  PfunGRL1 Protostelium aurantium var. fungivoru...
A0A0L0HDK0  1.000  425  SpunGRL1 gi|907093037|gb|KNC99049.1| hypotheti...

[383 rows x 3 columns]
```

In [7]:

```
#make a db with HMMS

if os.path.exists(qdir+runName+'profilephylo_v2_hhm_db*'):
    os.rm(qdir+runName+'profilephylo_v2_hhm_db*')

import hhsuitedb
hhsuitedb.add_new_files(  qdir+'HHfiles/*.hhm' , "hhm" ,  qdir+runName+'profilephylo_v2_hhm_db')
```

In [8]:

```
#run all v all comparison
hhms = glob.glob(  qdir+'HHfiles/*.hhm' )
print(hhms)
for q in hhms:
    p,output = runHHblits(q , name = q.split('.')[0].split('/')[-1] + 'allvall' , path= 'hhsearch ' , outdir = qdir+'ALLVSALL/' , db = qdir+runName+'profilephylo_v2_hhm_db' , iterations= 1 , ncores = ncores , ohhm = False, verbose = True , runName= 'test' , xargs = ' -mact .3')
```

```
['orco/HHfiles/VbraGRL1ProfphyloORCO.hhm', 'orco/HHfiles/NvecGRL1ProfphyloORCO.hhm', 'orco/HHfiles/AbakORCOProfphyloORCO.hhm', 'orco/HHfiles/VbraGRL2ProfphyloORCO.hhm', 'orco/HHfiles/AthaAT1G67570ProfphyloORCO.hhm', 'orco/HHfiles/TtraGRL1ProfphyloORCO.hhm', 'orco/HHfiles/CpriGRL1ProfphyloORCO.hhm', 'orco/HHfiles/AthaAT4G22270ProfphyloORCO.hhm', 'orco/HHfiles/TtraGRL6ProfphyloORCO.hhm', 'orco/HHfiles/AthaAT1G50630ProfphyloORCO.hhm', 'orco/HHfiles/TtraGRL2ProfphyloORCO.hhm', 'orco/HHfiles/TtraGRL5ProfphyloORCO.hhm', 'orco/HHfiles/VbraGRL5ProfphyloORCO.hhm', 'orco/HHfiles/MpusGRL1ProfphyloORCO.hhm', 'orco/HHfiles/AthaAT2G21080ProfphyloORCO.hhm', 'orco/HHfiles/VbraGRL3ProfphyloORCO.hhm', 'orco/HHfiles/AthaAT4G03820ProfphyloORCO.hhm', 'orco/HHfiles/VbraGRL6ProfphyloORCO.hhm', 'orco/HHfiles/SpurGRL1ProfphyloORCO.hhm', 'orco/HHfiles/SpunGRL1ProfphyloORCO.hhm', 'orco/HHfiles/SpalGRL1ProfphyloORCO.hhm', 'orco/HHfiles/VbraGRL4ProfphyloORCO.hhm', 'orco/HHfiles/TadhGRL1ProfphyloORCO.hhm', 'orco/HHfiles/DmelGr64aProfphyloORCO.hhm', 'orco/HHfiles/AthaAT3G20300ProfphyloORCO.hhm', 'orco/HHfiles/TtraGRL3ProfphyloORCO.hhm', 'orco/HHfiles/PfunGRL1ProfphyloORCO.hhm', 'orco/HHfiles/TtraGRL4ProfphyloORCO.hhm', 'orco/HHfiles/SkowGRL1ProfphyloORCO.hhm']
['orco/HHfiles/VbraGRL1ProfphyloORCO.hhm', 'VbraGRL1ProfphyloORCOallvall', 'hhsearch ', 'orco/ALLVSALL/', 'orco/ORCOprofilephylo_v2_hhm_db', 1, 8, 'test']
hhsearch  -cpu 8 -d orco/ORCOprofilephylo_v2_hhm_db -i orco/HHfiles/VbraGRL1ProfphyloORCO.hhm -o orco/ALLVSALL/VbraGRL1ProfphyloORCOallvalltest.hhr -n 1 -B 2000 -Z 2000  -mact .3
['orco/HHfiles/NvecGRL1ProfphyloORCO.hhm', 'NvecGRL1ProfphyloORCOallvall', 'hhsearch ', 'orco/ALLVSALL/', 'orco/ORCOprofilephylo_v2_hhm_db', 1, 8, 'test']
hhsearch  -cpu 8 -d orco/ORCOprofilephylo_v2_hhm_db -i orco/HHfiles/NvecGRL1ProfphyloORCO.hhm -o orco/ALLVSALL/NvecGRL1ProfphyloORCOallvalltest.hhr -n 1 -B 2000 -Z 2000  -mact .3
['orco/HHfiles/AbakORCOProfphyloORCO.hhm', 'AbakORCOProfphyloORCOallvall', 'hhsearch ', 'orco/ALLVSALL/', 'orco/ORCOprofilephylo_v2_hhm_db', 1, 8, 'test']
hhsearch  -cpu 8 -d orco/ORCOprofilephylo_v2_hhm_db -i orco/HHfiles/AbakORCOProfphyloORCO.hhm -o orco/ALLVSALL/AbakORCOProfphyloORCOallvalltest.hhr -n 1 -B 2000 -Z 2000  -mact .3
['orco/HHfiles/VbraGRL2ProfphyloORCO.hhm', 'VbraGRL2ProfphyloORCOallvall', 'hhsearch ', 'orco/ALLVSALL/', 'orco/ORCOprofilephylo_v2_hhm_db', 1, 8, 'test']
hhsearch  -cpu 8 -d orco/ORCOprofilephylo_v2_hhm_db -i orco/HHfiles/VbraGRL2ProfphyloORCO.hhm -o orco/ALLVSALL/VbraGRL2ProfphyloORCOallvalltest.hhr -n 1 -B 2000 -Z 2000  -mact .3
['orco/HHfiles/AthaAT1G67570ProfphyloORCO.hhm', 'AthaAT1G67570ProfphyloORCOallvall', 'hhsearch ', 'orco/ALLVSALL/', 'orco/ORCOprofilephylo_v2_hhm_db', 1, 8, 'test']
hhsearch  -cpu 8 -d orco/ORCOprofilephylo_v2_hhm_db -i orco/HHfiles/AthaAT1G67570ProfphyloORCO.hhm -o orco/ALLVSALL/AthaAT1G67570ProfphyloORCOallvalltest.hhr -n 1 -B 2000 -Z 2000  -mact .3
['orco/HHfiles/TtraGRL1ProfphyloORCO.hhm', 'TtraGRL1ProfphyloORCOallvall', 'hhsearch ', 'orco/ALLVSALL/', 'orco/ORCOprofilephylo_v2_hhm_db', 1, 8, 'test']
hhsearch  -cpu 8 -d orco/ORCOprofilephylo_v2_hhm_db -i orco/HHfiles/TtraGRL1ProfphyloORCO.hhm -o orco/ALLVSALL/TtraGRL1ProfphyloORCOallvalltest.hhr -n 1 -B 2000 -Z 2000  -mact .3
['orco/HHfiles/CpriGRL1ProfphyloORCO.hhm', 'CpriGRL1ProfphyloORCOallvall', 'hhsearch ', 'orco/ALLVSALL/', 'orco/ORCOprofilephylo_v2_hhm_db', 1, 8, 'test']
hhsearch  -cpu 8 -d orco/ORCOprofilephylo_v2_hhm_db -i orco/HHfiles/CpriGRL1ProfphyloORCO.hhm -o orco/ALLVSALL/CpriGRL1ProfphyloORCOallvalltest.hhr -n 1 -B 2000 -Z 2000  -mact .3
['orco/HHfiles/AthaAT4G22270ProfphyloORCO.hhm', 'AthaAT4G22270ProfphyloORCOallvall', 'hhsearch ', 'orco/ALLVSALL/', 'orco/ORCOprofilephylo_v2_hhm_db', 1, 8, 'test']
hhsearch  -cpu 8 -d orco/ORCOprofilephylo_v2_hhm_db -i orco/HHfiles/AthaAT4G22270ProfphyloORCO.hhm -o orco/ALLVSALL/AthaAT4G22270ProfphyloORCOallvalltest.hhr -n 1 -B 2000 -Z 2000  -mact .3
['orco/HHfiles/TtraGRL6ProfphyloORCO.hhm', 'TtraGRL6ProfphyloORCOallvall', 'hhsearch ', 'orco/ALLVSALL/', 'orco/ORCOprofilephylo_v2_hhm_db', 1, 8, 'test']
hhsearch  -cpu 8 -d orco/ORCOprofilephylo_v2_hhm_db -i orco/HHfiles/TtraGRL6ProfphyloORCO.hhm -o orco/ALLVSALL/TtraGRL6ProfphyloORCOallvalltest.hhr -n 1 -B 2000 -Z 2000  -mact .3
['orco/HHfiles/AthaAT1G50630ProfphyloORCO.hhm', 'AthaAT1G50630ProfphyloORCOallvall', 'hhsearch ', 'orco/ALLVSALL/', 'orco/ORCOprofilephylo_v2_hhm_db', 1, 8, 'test']
hhsearch  -cpu 8 -d orco/ORCOprofilephylo_v2_hhm_db -i orco/HHfiles/AthaAT1G50630ProfphyloORCO.hhm -o orco/ALLVSALL/AthaAT1G50630ProfphyloORCOallvalltest.hhr -n 1 -B 2000 -Z 2000  -mact .3
['orco/HHfiles/TtraGRL2ProfphyloORCO.hhm', 'TtraGRL2ProfphyloORCOallvall', 'hhsearch ', 'orco/ALLVSALL/', 'orco/ORCOprofilephylo_v2_hhm_db', 1, 8, 'test']
hhsearch  -cpu 8 -d orco/ORCOprofilephylo_v2_hhm_db -i orco/HHfiles/TtraGRL2ProfphyloORCO.hhm -o orco/ALLVSALL/TtraGRL2ProfphyloORCOallvalltest.hhr -n 1 -B 2000 -Z 2000  -mact .3
['orco/HHfiles/TtraGRL5ProfphyloORCO.hhm', 'TtraGRL5ProfphyloORCOallvall', 'hhsearch ', 'orco/ALLVSALL/', 'orco/ORCOprofilephylo_v2_hhm_db', 1, 8, 'test']
hhsearch  -cpu 8 -d orco/ORCOprofilephylo_v2_hhm_db -i orco/HHfiles/TtraGRL5ProfphyloORCO.hhm -o orco/ALLVSALL/TtraGRL5ProfphyloORCOallvalltest.hhr -n 1 -B 2000 -Z 2000  -mact .3
['orco/HHfiles/VbraGRL5ProfphyloORCO.hhm', 'VbraGRL5ProfphyloORCOallvall', 'hhsearch ', 'orco/ALLVSALL/', 'orco/ORCOprofilephylo_v2_hhm_db', 1, 8, 'test']
hhsearch  -cpu 8 -d orco/ORCOprofilephylo_v2_hhm_db -i orco/HHfiles/VbraGRL5ProfphyloORCO.hhm -o orco/ALLVSALL/VbraGRL5ProfphyloORCOallvalltest.hhr -n 1 -B 2000 -Z 2000  -mact .3
['orco/HHfiles/MpusGRL1ProfphyloORCO.hhm', 'MpusGRL1ProfphyloORCOallvall', 'hhsearch ', 'orco/ALLVSALL/', 'orco/ORCOprofilephylo_v2_hhm_db', 1, 8, 'test']
hhsearch  -cpu 8 -d orco/ORCOprofilephylo_v2_hhm_db -i orco/HHfiles/MpusGRL1ProfphyloORCO.hhm -o orco/ALLVSALL/MpusGRL1ProfphyloORCOallvalltest.hhr -n 1 -B 2000 -Z 2000  -mact .3
['orco/HHfiles/AthaAT2G21080ProfphyloORCO.hhm', 'AthaAT2G21080ProfphyloORCOallvall', 'hhsearch ', 'orco/ALLVSALL/', 'orco/ORCOprofilephylo_v2_hhm_db', 1, 8, 'test']
hhsearch  -cpu 8 -d orco/ORCOprofilephylo_v2_hhm_db -i orco/HHfiles/AthaAT2G21080ProfphyloORCO.hhm -o orco/ALLVSALL/AthaAT2G21080ProfphyloORCOallvalltest.hhr -n 1 -B 2000 -Z 2000  -mact .3
['orco/HHfiles/VbraGRL3ProfphyloORCO.hhm', 'VbraGRL3ProfphyloORCOallvall', 'hhsearch ', 'orco/ALLVSALL/', 'orco/ORCOprofilephylo_v2_hhm_db', 1, 8, 'test']
hhsearch  -cpu 8 -d orco/ORCOprofilephylo_v2_hhm_db -i orco/HHfiles/VbraGRL3ProfphyloORCO.hhm -o orco/ALLVSALL/VbraGRL3ProfphyloORCOallvalltest.hhr -n 1 -B 2000 -Z 2000  -mact .3
['orco/HHfiles/AthaAT4G03820ProfphyloORCO.hhm', 'AthaAT4G03820ProfphyloORCOallvall', 'hhsearch ', 'orco/ALLVSALL/', 'orco/ORCOprofilephylo_v2_hhm_db', 1, 8, 'test']
hhsearch  -cpu 8 -d orco/ORCOprofilephylo_v2_hhm_db -i orco/HHfiles/AthaAT4G03820ProfphyloORCO.hhm -o orco/ALLVSALL/AthaAT4G03820ProfphyloORCOallvalltest.hhr -n 1 -B 2000 -Z 2000  -mact .3
['orco/HHfiles/VbraGRL6ProfphyloORCO.hhm', 'VbraGRL6ProfphyloORCOallvall', 'hhsearch ', 'orco/ALLVSALL/', 'orco/ORCOprofilephylo_v2_hhm_db', 1, 8, 'test']
hhsearch  -cpu 8 -d orco/ORCOprofilephylo_v2_hhm_db -i orco/HHfiles/VbraGRL6ProfphyloORCO.hhm -o orco/ALLVSALL/VbraGRL6ProfphyloORCOallvalltest.hhr -n 1 -B 2000 -Z 2000  -mact .3
['orco/HHfiles/SpurGRL1ProfphyloORCO.hhm', 'SpurGRL1ProfphyloORCOallvall', 'hhsearch ', 'orco/ALLVSALL/', 'orco/ORCOprofilephylo_v2_hhm_db', 1, 8, 'test']
hhsearch  -cpu 8 -d orco/ORCOprofilephylo_v2_hhm_db -i orco/HHfiles/SpurGRL1ProfphyloORCO.hhm -o orco/ALLVSALL/SpurGRL1ProfphyloORCOallvalltest.hhr -n 1 -B 2000 -Z 2000  -mact .3
['orco/HHfiles/SpunGRL1ProfphyloORCO.hhm', 'SpunGRL1ProfphyloORCOallvall', 'hhsearch ', 'orco/ALLVSALL/', 'orco/ORCOprofilephylo_v2_hhm_db', 1, 8, 'test']
hhsearch  -cpu 8 -d orco/ORCOprofilephylo_v2_hhm_db -i orco/HHfiles/SpunGRL1ProfphyloORCO.hhm -o orco/ALLVSALL/SpunGRL1ProfphyloORCOallvalltest.hhr -n 1 -B 2000 -Z 2000  -mact .3
['orco/HHfiles/SpalGRL1ProfphyloORCO.hhm', 'SpalGRL1ProfphyloORCOallvall', 'hhsearch ', 'orco/ALLVSALL/', 'orco/ORCOprofilephylo_v2_hhm_db', 1, 8, 'test']
hhsearch  -cpu 8 -d orco/ORCOprofilephylo_v2_hhm_db -i orco/HHfiles/SpalGRL1ProfphyloORCO.hhm -o orco/ALLVSALL/SpalGRL1ProfphyloORCOallvalltest.hhr -n 1 -B 2000 -Z 2000  -mact .3
['orco/HHfiles/VbraGRL4ProfphyloORCO.hhm', 'VbraGRL4ProfphyloORCOallvall', 'hhsearch ', 'orco/ALLVSALL/', 'orco/ORCOprofilephylo_v2_hhm_db', 1, 8, 'test']
hhsearch  -cpu 8 -d orco/ORCOprofilephylo_v2_hhm_db -i orco/HHfiles/VbraGRL4ProfphyloORCO.hhm -o orco/ALLVSALL/VbraGRL4ProfphyloORCOallvalltest.hhr -n 1 -B 2000 -Z 2000  -mact .3
['orco/HHfiles/TadhGRL1ProfphyloORCO.hhm', 'TadhGRL1ProfphyloORCOallvall', 'hhsearch ', 'orco/ALLVSALL/', 'orco/ORCOprofilephylo_v2_hhm_db', 1, 8, 'test']
hhsearch  -cpu 8 -d orco/ORCOprofilephylo_v2_hhm_db -i orco/HHfiles/TadhGRL1ProfphyloORCO.hhm -o orco/ALLVSALL/TadhGRL1ProfphyloORCOallvalltest.hhr -n 1 -B 2000 -Z 2000  -mact .3
['orco/HHfiles/DmelGr64aProfphyloORCO.hhm', 'DmelGr64aProfphyloORCOallvall', 'hhsearch ', 'orco/ALLVSALL/', 'orco/ORCOprofilephylo_v2_hhm_db', 1, 8, 'test']
hhsearch  -cpu 8 -d orco/ORCOprofilephylo_v2_hhm_db -i orco/HHfiles/DmelGr64aProfphyloORCO.hhm -o orco/ALLVSALL/DmelGr64aProfphyloORCOallvalltest.hhr -n 1 -B 2000 -Z 2000  -mact .3
['orco/HHfiles/AthaAT3G20300ProfphyloORCO.hhm', 'AthaAT3G20300ProfphyloORCOallvall', 'hhsearch ', 'orco/ALLVSALL/', 'orco/ORCOprofilephylo_v2_hhm_db', 1, 8, 'test']
hhsearch  -cpu 8 -d orco/ORCOprofilephylo_v2_hhm_db -i orco/HHfiles/AthaAT3G20300ProfphyloORCO.hhm -o orco/ALLVSALL/AthaAT3G20300ProfphyloORCOallvalltest.hhr -n 1 -B 2000 -Z 2000  -mact .3
['orco/HHfiles/TtraGRL3ProfphyloORCO.hhm', 'TtraGRL3ProfphyloORCOallvall', 'hhsearch ', 'orco/ALLVSALL/', 'orco/ORCOprofilephylo_v2_hhm_db', 1, 8, 'test']
hhsearch  -cpu 8 -d orco/ORCOprofilephylo_v2_hhm_db -i orco/HHfiles/TtraGRL3ProfphyloORCO.hhm -o orco/ALLVSALL/TtraGRL3ProfphyloORCOallvalltest.hhr -n 1 -B 2000 -Z 2000  -mact .3
['orco/HHfiles/PfunGRL1ProfphyloORCO.hhm', 'PfunGRL1ProfphyloORCOallvall', 'hhsearch ', 'orco/ALLVSALL/', 'orco/ORCOprofilephylo_v2_hhm_db', 1, 8, 'test']
hhsearch  -cpu 8 -d orco/ORCOprofilephylo_v2_hhm_db -i orco/HHfiles/PfunGRL1ProfphyloORCO.hhm -o orco/ALLVSALL/PfunGRL1ProfphyloORCOallvalltest.hhr -n 1 -B 2000 -Z 2000  -mact .3
['orco/HHfiles/TtraGRL4ProfphyloORCO.hhm', 'TtraGRL4ProfphyloORCOallvall', 'hhsearch ', 'orco/ALLVSALL/', 'orco/ORCOprofilephylo_v2_hhm_db', 1, 8, 'test']
hhsearch  -cpu 8 -d orco/ORCOprofilephylo_v2_hhm_db -i orco/HHfiles/TtraGRL4ProfphyloORCO.hhm -o orco/ALLVSALL/TtraGRL4ProfphyloORCOallvalltest.hhr -n 1 -B 2000 -Z 2000  -mact .3
['orco/HHfiles/SkowGRL1ProfphyloORCO.hhm', 'SkowGRL1ProfphyloORCOallvall', 'hhsearch ', 'orco/ALLVSALL/', 'orco/ORCOprofilephylo_v2_hhm_db', 1, 8, 'test']
hhsearch  -cpu 8 -d orco/ORCOprofilephylo_v2_hhm_db -i orco/HHfiles/SkowGRL1ProfphyloORCO.hhm -o orco/ALLVSALL/SkowGRL1ProfphyloORCOallvalltest.hhr -n 1 -B 2000 -Z 2000  -mact .3
```

In [9]:

```
# parse hhr files and make dist kernel

def cleanID(ID):
    
    if '|PDBID' in ID:
        ID = ID.split('|PDBID')[0]
    
    elif '|' in ID:
        ID = ID.split('|')[0].split()[0]
    elif '.' in ID:
        ID = ID.split('.')[0]
    elif '/' in ID:
        ID = ID.split('/')[0]
    elif 'hmmercut' in ID:
        ID = ID.split('hmmercut')[0]
    ID = ID.split()[0]
    ID = ID.strip()
    return ID


def HHSearch_parseTo_DMandNX(hhrs):
    clusternames = []
        
    for i,hhr in enumerate(hhrs):
        print(hhr)
        profile = HHOutputParser(alignments=False).parse_file(hhr)
        if cleanID(profile.query_name) not in clusternames:
            print(cleanID(profile.query_name))
            clusternames.append(cleanID(profile.query_name))
         
    print(clusternames)
    evalDM = np.ones( (len(clusternames),len(clusternames) ))
    pvalDM = np.ones( (len(clusternames),len(clusternames) ))
    scoreDM = np.zeros( (len(clusternames),len(clusternames) ))
    SSDM = np.zeros( (len(clusternames),len(clusternames) ))
    probaDM = np.zeros( (len(clusternames),len(clusternames) ))
    lenDM =  np.ones( (len(clusternames),len(clusternames) ))
    
    NX = nx.Graph()
    
    for i,hhr in enumerate(hhrs):
        protlist = []
        profile = HHOutputParser(alignments=False).parse_file(hhr)
        for hit in profile:
            DMscore = float(hit.evalue)
            proba = hit.probability
            if 'anchor' not in hit.id and 'anchor' not in profile.query_name:
                i = clusternames.index(cleanID(hit.id))
                j = clusternames.index(cleanID(profile.query_name))

                if hit.evalue < evalDM[i,j]:
                    evalDM[i,j] = hit.evalue
                    evalDM[j,i] = evalDM[i,j]

                if hit.pvalue < pvalDM[i,j]:
                    pvalDM[i,j] = hit.pvalue
                    pvalDM[j,i] = pvalDM[i,j]

                if scoreDM[i,j] < hit.score:
                    scoreDM[i,j] = hit.score
                    scoreDM[j,i] = scoreDM[i,j]

                if SSDM[i,j] < hit.ss_score:
                    SSDM[i,j] = hit.ss_score
                    SSDM[j,i] = SSDM[i,j]


                if probaDM[i,j] < hit.probability:
                    probaDM[i,j] = hit.probability
                    probaDM[j,i] = probaDM[i,j]

                #use smallest of the two prots
                if lenDM[i,j] == 1 or lenDM[i,j] > hit.qlength:
                    lenDM[i,j] = hit.qlength
                    lenDM[j,i] = lenDM[i,j]

            if hit.id != profile.query_name :
                NX.add_edge( hit.id , profile.query_name )
                NX[hit.id][profile.query_name]['score']= hit.score
    return probaDM, evalDM ,pvalDM,  lenDM , scoreDM, SSDM, NX , clusternames
```

```
/home/cactuskid13/miniconda3/envs/pyprofiler3/lib/python3.7/site-packages/matplotlib/__init__.py:886: MatplotlibDeprecationWarning: 
examples.directory is deprecated; in the future, examples will be found relative to the 'datapath' directory.
  "found relative to the 'datapath' directory.".format(key))
/home/cactuskid13/miniconda3/envs/pyprofiler3/lib/python3.7/site-packages/statsmodels/tools/_testing.py:19: FutureWarning: pandas.util.testing is deprecated. Use the functions in the public API at pandas.testing instead.
  import pandas.util.testing as tm
```

In [14]:

```
print(glob.glob(qdir+'ALLVSALL/*.hhr'))
probaDM, evalDM ,pvalDM,  lenDM , scoreDM, SSDM, NX , clusternames = HHSearch_parseTo_DMandNX(glob.glob(qdir+ 'ALLVSALL/*.hhr'))
```

```
['orco/ALLVSALL/AthaAT2G21080ProfphyloORCOallvalltest.hhr', 'orco/ALLVSALL/AthaAT1G67570ProfphyloORCOallvalltest.hhr', 'orco/ALLVSALL/VbraGRL5ProfphyloORCOallvalltest.hhr', 'orco/ALLVSALL/PfunGRL1ProfphyloORCOallvalltest.hhr', 'orco/ALLVSALL/NvecGRL1ProfphyloORCOallvalltest.hhr', 'orco/ALLVSALL/DmelGr64aProfphyloORCOallvalltest.hhr', 'orco/ALLVSALL/TtraGRL5ProfphyloORCOallvalltest.hhr', 'orco/ALLVSALL/MpusGRL1ProfphyloORCOallvalltest.hhr', 'orco/ALLVSALL/SpalGRL1ProfphyloORCOallvalltest.hhr', 'orco/ALLVSALL/VbraGRL1ProfphyloORCOallvalltest.hhr', 'orco/ALLVSALL/SkowGRL1ProfphyloORCOallvalltest.hhr', 'orco/ALLVSALL/AthaAT3G20300ProfphyloORCOallvalltest.hhr', 'orco/ALLVSALL/SpunGRL1ProfphyloORCOallvalltest.hhr', 'orco/ALLVSALL/TtraGRL3ProfphyloORCOallvalltest.hhr', 'orco/ALLVSALL/AthaAT4G22270ProfphyloORCOallvalltest.hhr', 'orco/ALLVSALL/SpurGRL1ProfphyloORCOallvalltest.hhr', 'orco/ALLVSALL/AthaAT4G03820ProfphyloORCOallvalltest.hhr', 'orco/ALLVSALL/TtraGRL6ProfphyloORCOallvalltest.hhr', 'orco/ALLVSALL/TtraGRL1ProfphyloORCOallvalltest.hhr', 'orco/ALLVSALL/AthaAT1G50630ProfphyloORCOallvalltest.hhr', 'orco/ALLVSALL/TadhGRL1ProfphyloORCOallvalltest.hhr', 'orco/ALLVSALL/VbraGRL6ProfphyloORCOallvalltest.hhr', 'orco/ALLVSALL/AbakORCOProfphyloORCOallvalltest.hhr', 'orco/ALLVSALL/TtraGRL4ProfphyloORCOallvalltest.hhr', 'orco/ALLVSALL/VbraGRL4ProfphyloORCOallvalltest.hhr', 'orco/ALLVSALL/VbraGRL3ProfphyloORCOallvalltest.hhr', 'orco/ALLVSALL/CpriGRL1ProfphyloORCOallvalltest.hhr', 'orco/ALLVSALL/TtraGRL2ProfphyloORCOallvalltest.hhr', 'orco/ALLVSALL/VbraGRL2ProfphyloORCOallvalltest.hhr']
orco/ALLVSALL/AthaAT2G21080ProfphyloORCOallvalltest.hhr
AthaAT2G21080
orco/ALLVSALL/AthaAT1G67570ProfphyloORCOallvalltest.hhr
AthaAT1G67570
orco/ALLVSALL/VbraGRL5ProfphyloORCOallvalltest.hhr
VbraGRL5
orco/ALLVSALL/PfunGRL1ProfphyloORCOallvalltest.hhr
PfunGRL1
orco/ALLVSALL/NvecGRL1ProfphyloORCOallvalltest.hhr
NvecGRL1
orco/ALLVSALL/DmelGr64aProfphyloORCOallvalltest.hhr
DmelGr64a
orco/ALLVSALL/TtraGRL5ProfphyloORCOallvalltest.hhr
TtraGRL5
orco/ALLVSALL/MpusGRL1ProfphyloORCOallvalltest.hhr
MpusGRL1
orco/ALLVSALL/SpalGRL1ProfphyloORCOallvalltest.hhr
SpalGRL1
orco/ALLVSALL/VbraGRL1ProfphyloORCOallvalltest.hhr
VbraGRL1
orco/ALLVSALL/SkowGRL1ProfphyloORCOallvalltest.hhr
SkowGRL1
orco/ALLVSALL/AthaAT3G20300ProfphyloORCOallvalltest.hhr
AthaAT3G20300
orco/ALLVSALL/SpunGRL1ProfphyloORCOallvalltest.hhr
SpunGRL1
orco/ALLVSALL/TtraGRL3ProfphyloORCOallvalltest.hhr
TtraGRL3
orco/ALLVSALL/AthaAT4G22270ProfphyloORCOallvalltest.hhr
AthaAT4G22270
orco/ALLVSALL/SpurGRL1ProfphyloORCOallvalltest.hhr
SpurGRL1
orco/ALLVSALL/AthaAT4G03820ProfphyloORCOallvalltest.hhr
AthaAT4G03820
orco/ALLVSALL/TtraGRL6ProfphyloORCOallvalltest.hhr
TtraGRL6
orco/ALLVSALL/TtraGRL1ProfphyloORCOallvalltest.hhr
TtraGRL1
orco/ALLVSALL/AthaAT1G50630ProfphyloORCOallvalltest.hhr
AthaAT1G50630
orco/ALLVSALL/TadhGRL1ProfphyloORCOallvalltest.hhr
TadhGRL1
orco/ALLVSALL/VbraGRL6ProfphyloORCOallvalltest.hhr
VbraGRL6
orco/ALLVSALL/AbakORCOProfphyloORCOallvalltest.hhr
AbakORCO
orco/ALLVSALL/TtraGRL4ProfphyloORCOallvalltest.hhr
TtraGRL4
orco/ALLVSALL/VbraGRL4ProfphyloORCOallvalltest.hhr
VbraGRL4
orco/ALLVSALL/VbraGRL3ProfphyloORCOallvalltest.hhr
VbraGRL3
orco/ALLVSALL/CpriGRL1ProfphyloORCOallvalltest.hhr
CpriGRL1
orco/ALLVSALL/TtraGRL2ProfphyloORCOallvalltest.hhr
TtraGRL2
orco/ALLVSALL/VbraGRL2ProfphyloORCOallvalltest.hhr
VbraGRL2
['AthaAT2G21080', 'AthaAT1G67570', 'VbraGRL5', 'PfunGRL1', 'NvecGRL1', 'DmelGr64a', 'TtraGRL5', 'MpusGRL1', 'SpalGRL1', 'VbraGRL1', 'SkowGRL1', 'AthaAT3G20300', 'SpunGRL1', 'TtraGRL3', 'AthaAT4G22270', 'SpurGRL1', 'AthaAT4G03820', 'TtraGRL6', 'TtraGRL1', 'AthaAT1G50630', 'TadhGRL1', 'VbraGRL6', 'AbakORCO', 'TtraGRL4', 'VbraGRL4', 'VbraGRL3', 'CpriGRL1', 'TtraGRL2', 'VbraGRL2']
```

In [21]:

```
g = sns.clustermap( probaDM , xticklabels=clusternames  , yticklabels=clusternames , figsize = (20,20) )
```
